# Complement activation: in maternal and placental pathology of preeclampsia

**DOI:** 10.1101/2025.10.03.680407

**Authors:** Manu T Banadakoppa, Madhulatha S Chauhan, Moises J Tacam, Allyson W Nevins, Antonio H Ruano, Simone H Ruano, Jon A Fuson, Chandra Yallampalli

## Abstract

Preeclampsia is a multifactorial, pregnancy-related disorder characterized by new-onset hypertension and proteinuria, with distinct early- and late-onset forms linked to varying placental and maternal pathologies. While complement system dysregulation has been implicated in the pathogenesis of preeclampsia, its causal relationship and molecular mechanisms remain unclear. In mice, complement receptor 1-related protein y (Crry) functions as a critical complement regulator at the fetal-maternal interface, essential for early embryonic survival. Complete Crry deficiency is embryonically lethal, complicating in vivo studies of complement activation in pregnancy. Using an alternative strategy, we developed a placenta-specific, doxycycline-inducible shRNA mouse model utilizing Cyp19-driven Cre recombinase and a Tet-On system to downregulate Crry in a dose-dependent manner 9.5 days post coitus, effectively restricting complement activation to the placenta.

Using this model, we demonstrate that early gestation but sustained placental complement activation impairs maternal heart and liver adaptation, reduces placental efficiency, and causes fetal growth restriction, mimicking early-onset preeclampsia. Conversely, delayed complement activation induces a phenotype more consistent with late-onset preeclampsia features without placental pathology. In early-onset preeclampsia-like phenotype, the fetal growth restriction is accompanied by placental glycogen storage deficiency, and impaired hormonal function. Maternal glucose metabolism is not affected but compensatory adaptations in lipid metabolism occur although insufficient to offset fetal growth restriction. This novel model reveals that the timing of placental complement activation dictates the spectrum of preeclampsia-like pathology, providing mechanistic insights into the pathophysiology of preeclampsia.

## INTRODUCTION

Emerging evidence implicates dysregulation of the complement system-a key component of innate immunity-in the pathogenesis of preeclampsia. The products of complement activation are increased in the circulation of pregnant women (1). These products are further increased in preeclampsia patients suggesting an association between preeclampsia and increased complement activation (2–6). In preeclampsia patients, complement split products are deposited on the placenta suggesting an increased local complement activation at the fetal-maternal interface (7;8).

Circumstantial evidence obtained from various animal models supports a pathophysiological role for complement activation in preeclampsia. In mice, treatment with a site-specific complement inhibitor abrogates placental and fetal preeclampsia like-phenotypes of CBA/J females mated with DBA/2 males which are otherwise hypertensive (9). Site-specific complement inhibition during pregnancy abrogates fetal and placental preeclampsia-like phenotypes in mildly hypertensive BPH/5 mice (10). In human angiotensin II autoantibody induced preeclampsia mouse model, inhibition of C3a-C3aR signaling axis with a C3aR antagonist prevents hypertension and feto-placental preeclampsia-like phenotype (11).

Although implicated in pathophysiology, weather complement activation has a causal effect on the development of preeclampsia is not known. In a recent study using a lentiviral mediated shRNA mouse model, we have reported that downregulation of complement inhibitor Crry leads to preeclampsia-like phenotype (12). Here we have extended this study using a more vigorous genetically modified mouse model of Crry downregulation and show that placenta-specific increase in complement activation is a common etiologic factor for early- and late-onset types of preeclampsia.

## RESULTS

### Opposing Effects of Crry Suppression and Feedback Upregulation Shape Complement Activation on placenta

Rodent-specific complement pathway regulator Crry has similarity in function with more general complement regulators-decay accelerating factor (DAF, CD55) and membrane cofactor protein (MCP, CD46) (13). In rodents, MCP is predominantly expressed only in testes (14). In mouse placenta, DAF expression is detectable at around 10.5 days post coitus (dpc), much later than the expression of Crry (15;16). Consequently, Crry is very critical for complement regulation during early embryonic development. Complete deletion of Crry leads to embryo demise by 9.5 dpc (17). To test if increased complement activation leads to preeclampsia phenotype in mouse, we used shRNA approach to downregulate Crry expression in a placenta-specific manner, simultaneously avoiding lethal phenotype. To restrict complement activation to the placenta, we employed a mouse model based on Cre-Lox*P* technology. In this mouse model, Cre recombinase expression was initiated by the human Cyp19 promoter (18). The Tet-On system with doxycycline dependent high-efficiency reverse tetracycline transactivator (rtTA) (19) was used to induce Crry shRNA in a dose-dependent manner, and after 9.5 dpc. Since maternal inheritance provides strongest and consistent expression through Cyp19 promoter (18), we mated female Cyp19^Cre/Cre^ mice with male rtTA^+/+^Crry^LSL-RNAi^ to obtain timed pregnancies.

Since variable Cre recombination efficiency and weak to moderate Cre expression throughout the internal tissues of some fetuses (≤ 4.5%) were previously reported in Cyp19-Cre transgenic mice (20), (18), we examined recombination efficiency in placentas and various internal organs of offspring from Cyp19^Cre/Cre^ female x rtTA^+/+^Crry^LSL-RNAi^ male mating. Recombination was tested using primers designed to bind outside the Lox*P* sites **(Figure S1A)**. Cre recombination occurred in approximately 80% (27 out of 34 from 4 dams) of placentas. In offspring, Cre recombination was less than 2% throughout the internal organs which is consistent with the previous report (18).

An individual’s intrinsic complement activation potential depends on expression levels and inherited variations in function of complement proteins resulting in population-wide heterogeneity in complement activation when triggered (21;22). This heterogeneity of complement activation exists in pregnant women so that there is a possibility of different degrees of complement activation increase during pregnancy. Therefore, we set out to examine how two different levels of complement activation increase affect pregnancy outcomes using 37.5 µg/mL (low-dose) and 75 µg/mL (high-dose) doxycycline to induce Crry shRNA starting from gestation day (GD) 10.5 based on our previous study (23). We compared the Crry protein levels in placentas from the low- and the high-dose group with the control group by Western blot analysis at different time points after the doxycycline treatment **(Figure 1A)**. After 6 hours of doxycycline treatment, placental Crry protein levels decreased by about 80% in the low- and the high dose groups compared to control group **(Figure 1B)**. At GD 11.5, placental Crry protein levels decreased by about 60% in the low-dose group compared to the control group. There was no difference in placental Crry levels between the high-dose and the control groups **(Figure 1C)**. At GD13.5, placental Crry levels decreased by about 50% and 30% in the low- and the high-dose groups respectively, compared to the control group **(Figure 1D)**. At GD17.5, placental Crry level were not different between the low-dose and the control groups but decreased by about 60% in the high-dose group compared to the control group **(Figure 1E)**. Upregulation of Crry expression in the low-dose group continued at GD17.5 while there was no difference in its expression between the control and the high-dose groups. These data suggested that a consistent feedback upregulation response occurs after initial downregulation of Crry in doxycycline-treated groups. Therefore, the net Crry mRNA levels throughout the gestation were determined by a balance between knockdown effect of shRNA and feedback upregulation response. These opposing suppression and feedback upregulation effects were reflected in Crry protein levels. Placental Crry protein levels were decreased by approximately 50% and 35% in the low- and the high-dose groups respectively compared to the control group at GD13.5 **(Figure 1 C, D)**. At GD 17.5, placental Crry protein levels were not significantly different between the control and the low-dose groups but decreased by more than 50% in the high-dose group compared to the control **(Figure 1 E, F)**.

The central reaction in the complement activation cascade is the conversion of C3 protein into C3b and C3a fragments resulting in the deposition of C3b on nearby cell surfaces. Surface bound C3b molecules are rapidly inactivated into iC3b by fragmentation through various regulators including Factor I, CD46 and Factor H proteins (24). Initially, we performed Western blot analysis to identify the type of fragment detectable in tissue lysates from mouse placentas by comparing with purified mouse proteins. We detected the presence of iC3b in placental tissue lysates but could not detect C3b, possibly due to complete degradation into iC3b. We were able to detect intact C3 protein also in these placentas **(Figure S1B)**. We then compared iC3b levels in placentas from the three groups by Western blot analysis at different time points during the gestation **(Figure 2A)**. Six hours after the doxycycline treatment and at GD11.5, placental iC3b levels were not significantly different between the groups **(Figure 2 B, C).** At GD13.5. iC3b levels in placentas from the low-dose group increased by approximately 200% compared to the control and the high-dose groups consistent with decreased Crry protein levels. There was no difference in iC3b levels between the high-dose and the control groups. **(Figure 2 D)**. At GD17.5, iC3b levels in placentas from the high-dose group increased by approximately 80% compared to the control and the low-dose groups consistent with decreased Crry levels. There was no difference in iC3b levels between the low-dose and the control groups **(Figure 2 E)**. When iC3b protein levels were compared between GD13.5 and GD17.5 for each group **(Figure 2F)**, there was approximately 200% and 400% increase in the control and the high-dose groups respectively without any changes in the low-dose group **(Figure 2 G-I)**. These data suggested that, in the low-dose group, complement activation substantially increased at GD13.5 and remained elevated as gestation progressed to 17.5 days. In the control and the high-dose groups, complement activation did not increase substantially at GD13.5, but modest and large increases occurred in the control and the low-dose groups respectively as gestation progressed from 13.5 to 17.5 days.

The C3 protein in circulation is the major source providing C3 for the central reaction in the complement pathway (25). Sine placenta can also produce C3 locally (26;27), we sought to evaluate relative contribution from these two sources to the increased complement activation by measuring serum C3 levels and assessing C3 protein in placentas. At GD 13.5, the mean serum C3 concentrations were decreased by approximately 0.15 mg/mL in the low- and the high-dose groups compared to the control group, but the differences were not statistically significant **(Figure 2 J)**. However, at GD 17.5, the mean serum C3 concentration was significantly decreased in the high-dose group compared to the control group **(Figure 2K)**. When the mean serum C3 concentrations were compared between the two gestational time points, we observed that C3 concentration was decreased by 0.1 mg/mL from GD17.5 to GD13.5 in the control group though the difference was not statistically significant **(Figure 2L)**. In the low-dose group, mean serum C3 concentrations were similar at GD13.5 and 17.5 **(Figure 2M)**. In the high-dose group, the mean serum C3 concentration significantly decreased from GD13.5 to GD17.5 **(Figure 2N)**. Western blot analysis revealed that the C3 protein levels in the placentas from the high-dose group were significantly increased at GD13.5 compared to the control and the low-dose groups **(Figure 2 O, P)**. At GD17.5, C3 protein levels in placentas from the low-dose group significantly increased compared to the control group **(Figure 2 Q, R)**. When placental C3 levels were compared between GD13.5 and GD17.5, significant decrease occurred only in the control group **(Figure 2 S-V)**. These data suggested that, at GD13.5, C3 protein from maternal circulation was used for complement activation in the low- and the high-dose groups. Additionally, C3 protein from placenta was also used for complement activation in the low-dose group. As gestation progressed, C3 from circulation was used significantly only in the high-dose group while C3 from placenta was exclusively used in the control group for complement activation.

Taken together, these data suggested that in the low- and the high-dose groups, the trajectories of Crry downregulation and complement activation were different. An early, and moderate but sustained complement activation occurred in the low-dose group whereas, late but high-level complement activation occurred in the high-dose group. An initial downregulation of Crry was opposed by a feedback upregulation. The net effect of shRNA-mediated downregulation and opposing feedback upregulation was decreased Crry protein levels at GD13.5 in both the low- and the high-dose groups. The complement activation increased moderately at GD13.5 only in the low-dose group indicating that Crry protein levels were still sufficient to prevent the complement activation in the high-dose group though its levels decreased. As gestation progressed, feedback upregulation was able to partially restore Crry protein levels in the low-dose group. However, in the high dose group, continuous high-level suppression caused further decrease in the Crry protein levels by GD17.5 leading to substantial increase in complement activation.

### Increased Complement Activation Impairs Cardiovascular Adaptation and Elevates Blood Pressure During Pregnancy

Litter size, fetal reabsorption percentages and body weights of dams at GD13.5 and GD17.5 were not significantly different between the groups **(Figure S2 A-F)**. Non-invasive tail-cuff measurements of blood pressure in pregnant mice revealed that systolic, diastolic, and mean arterial pressure were significantly increased in the low- and the high-dose doxycycline groups compared to the controls. Within the doxycycline treated groups, blood pressure was significantly higher in the high-dose group compared to the low-dose group **(Figure 3 A-C).** Since uteroplacental blood circulation is impaired in women with preeclampsia (28), and in mice models that show preeclampsia phenotype (29), we measured uterine artery blood flow using non-invasive ultrasound and Doppler imaging **(Figure 3D)**. The uterine artery blood flow patterns at GD15.5 were altered in both the low- and the high-dose groups compared to the control animals as indicated by increase in uterine artery resistive index (UARI) (**Figure 3E).** However, UARI was not different between the low- and the high-dose groups. Preeclampsia is associated with endothelial injury due to the deficiency of endothelial nitric oxide synthase enzyme (eNOS) (30). Immunostaining of mesenteric arteries for eNOS revealed that eNOS protein expression was similar in dams from three groups at GD13.5. However, as gestation progressed to 17.5 days, eNOS protein expression in mesenteric arteries from the control mice increased abundantly. In mice from the low- and the high-dose groups, mesenteric artery eNOS protein levels did not increase as gestation progressed to GD17.5 **(Figure 3F)**. Normal pregnancy is associated with hypertrophic growth in the heart to compensate for increased cardiac output (31;32). We found that the heart weight increased in mice from the control group as pregnancy progressed from 13.5 to 17.5 days. However, the heart weight regressed both in the low- and the high-dose dams compared to the control group with the pregnancy progression **(Figure 3 G, H)** suggesting that the normal adaptation of heart to pregnancy failed in those animals. These data indicated that although complement activation followed different trajectories with the low- and the high shRNA levels, the increase in blood pressure and impaired cardiovascular adaptation were common in both the conditions. However, the magnitude of increase in blood pressure was apparently dependent on the magnitude of increase in the complement activation.

### Hepatic Adaptation is Impaired and Glomerular Endotheliosis Occurred Depending on the Trajectory of Complement Activation During Pregnancy

During normal pregnancy the liver undergoes enhanced growth to adapt to the increasing energy demand from the developing fetus and for the detoxification of fetal metabolites (33;34). As expected, the liver weight increased from GD13.5 to GD17.5 in the control dams. On the other hand, the liver weight decreased from GD13.5 to GD17.5 in the low-dose dams. Further, the mean of the liver weight was significantly reduced in the low dose group compared to the control and the high-dose groups at GD17.5. In the high-dose group, the liver weight slightly regressed from GD13.5 to GD17.5 but was not significantly different from the control group **(Figure 4 A, B)**. These data suggested that normal adaptation of the liver to pregnancy was impaired in the mice treated with the low-dose doxycycline. However, serum aspartate transferase (ALT), serum alanine transaminase (AST) and serum lactate dehydrogenase levels were not different between the three groups, suggesting that extensive liver damage was not present in mice with increased complement activation **(Figure S3 A-C)**.

During normal pregnancy, the spleen also transiently enlarges due to proliferation of erythroid cells and leukocytes (35–37). Pregnancy associated spleen enlargement peaks around GD13 in mice and then regresses to non-pregnant weight before parturition (37). As expected, the spleen weight decreased from GD13.5 to GD17.5 in all three groups. However, there was no significant change in the spleen weight between the groups **(Figure S3 D, E)**. Further, there was no significant difference in the weights of the lungs and the kidneys between the groups **(Figure S3 F-I)**. However, bright field microscopy of hematoxylin and eosin-stained kidney sections revealed that glomerular endotheliosis was present in the kidneys from the low- and the high-dose dams **(Figure 4C).** Concurrently, urine albumin to creatinine ratio was significantly increased in the high-dose group compared to the control and the low-dose groups suggesting impaired kidney function only in mice from the high-dose group **(Figure 4D)**.

### Early and Sustained Increase in Complement Activation During Pregnancy Impairs Placental Endocrine Function and Reduces Fetal Growth

The mean weight of the fetuses from the low-dose group increased by 8% and 9% compared to the control and the high-dose groups respectively at GD13.5. Placental weights were not significantly different between the groups at GD13.5. Placental efficiency was significantly increased in the low-dose group compared to the control and the high-dose groups at GD13.5 **(Figure 5 A-C)**. However, at GD17.5, the mean weight of the fetuses from the low-dose group reduced by 5% compared to the control group. Placental weights were not different between the groups at GD17.5. But the placental efficiency was reduced in the low-dose group compared to the control **(Figure 5 D-F)**. These data indicated that fetal growth restriction occurred apparently due to reduced placenta efficiency in the low-dose group.

Several placental factors such as defects in labyrinth and junctional zones, and placental hormone dysfunction are known to be strongly associated with fetal growth restriction (38–41). Therefore, we first analyzed hematoxylin and eosin (H&E) stained sections of the placentas to find out if the increase in complement activation induced any structural defects in the placenta **(Figure 5 G, H)**. At GD 13.5, there were no differences in the sizes of either labyrinth or junctional zones between the three groups. The labyrinth zone area as a percentage of total area of the placenta was not different between the groups at GD17.5 **(Figure 5 I)**. The junctional zone area percentage at GD17.5 was significantly reduced in the low-dose group in comparison to the control group **(Figure 5 J)**. The junctional zone produces several prolactin hormones such as placental lactogen 1 (prl3d1) and placental lactogen 2 (prl3b1) (42). We assessed the expression levels of prl3d1 and prl3b1 in the placentas. The mRNA level of prl3d1 was significantly decreased in the low-dose group compared to the high-dose group at GD 13.5. By GD17.5, prl3d1 mRNA level in the low-dose group was significantly increased compared to the control and the high-dose groups **(Figure 5 K, L)**. The mRNA level of prl3b1 was significantly increased in the low-dose group in comparison with the control and the high-dose groups at both GD 13.5 and GD 17.5 **(Figure 5 M, N)**. These data suggested that the fetal growth restriction in the low-dose group was associated with reduced junctional zone and altered endocrine function in the placenta.

### Glycogen Storage Deficiency in the Placenta Underlies Fetal Growth Restriction Independent of Maternal Glucose Homeostasis

Placental glycogen store provides a source of glucose to the fetus in late gestation (43) suggesting that glycogen storage deficiency may play a role in fetal growth restriction. In several gene deletion experiments in mice, placental glycogen store deficiency is frequently associated with fetal growth restriction (44). Therefore, we measured the placental glycogen content using periodic acid-Schiff staining method **(Figure 6 A, B)** to assess if placental glycogen storage deficiency underlies the fetal growth restriction in the low-dose group. We observed slightly increased placental glycogen levels at GD17.5 compared to GD13.5 in all three groups. However, placental glycogen levels were significantly reduced in the low-dose group in comparison with the control and the high-dose groups at GD17.5 **(Figure 6 C, D)**. To assess if changes in the maternal glucose metabolism induced the glycogen storage deficiency in the low-dose group, we measured insulin levels and performed glucose tolerance (GTT) and insulin tolerance tests (ITT). The fasting glucose levels were not different between the groups. Glucose tolerance and area under the curve for GTT were not different between the groups **(Figure 6 E, F)**. As expected during pregnancy, the dams from all three groups showed insulin intolerance. However, insulin tolerance and area under the curve for ITT were not different between the groups **(Figure 6 G, H)**. There were no differences in serum insulin levels between the groups at GD13.5 and 17.5 **(Figure 6 I, J)**. Taken together, these results indicated that glycogen storage deficiency occurred in the low-dose doxycycline group. The glucose metabolism was not impaired in dams from the low-dose group in comparison with the dams from the control and the high-dose groups suggesting that placental glycogen storage deficiency was independent of maternal glucose metabolism.

### Placental Adaptation via Lipid Transporter Upregulation in the Context of Complement-Associated Glycogen Deficiency

To examine if changes in the maternal lipid metabolism triggered placental glycogen deficiency and fetal growth restriction, we analyzed circulating lipid levels in dams. The serum triglyceride and leptin levels were not different in the dams from three groups at GD 17.5 **(Figure 7 A, B)**. Concurrently, ex vivo analysis of lipolysis in chunks of perigonadal fat tissues obtained from these dams revealed that the rate of lipolysis was not different between the three groups **(Figure 7 C, D)**. We then analyzed if expression levels of fatty acid transporters (FATs) and fatty acid binding proteins (FABPs) which play a vital role in transportation of lipids across the placenta, lipid storage and mobilization (45;46) were altered. Among FATs, fatty acid translocase (fat) and fatty acid transport protein 6 (fatp6) mRNA levels increased in the low-dose group at GD 13.5 compared to the control and the high-dose groups **(Figure 7 E, F)**. Among FABPs, the mRNA levels of FABP4 increased in the low-dose group at GD 17.5 compared to the control group **(Figure 7 G)**. The mRNA levels of fabp5 in the low-dose group increased at 1GD 3.5 compared to the control group and increased at GD 17.5 compared to both the control and the high-dose groups **(Figure 7 H, I)**. Taken together, these data suggested that lipid transportation and mobilization increased in placentas from the low-dose group possibly to compensate for glycogen deficiency.

## DISCUSSION

For a long time, preeclampsia has been known to be a multifactorial syndrome mainly manifested as new onset of hypertension along with proteinuria (47). Preeclampsia is often associated with fetal growth restriction especially in case of early-onset severe preeclampsia (48). Early-onset preeclampsia comprises placenta pathology whereas, late-onset preeclampsia frequently leads to mild symptoms with less involvement of placenta pathology and fewer adverse fetal outcomes (49–51). From an etiological point of view, early- and late-onset preeclampsia are considered as two distinct entities with slightly overlapping pathophysiology (52). Several experimental animal models have been developed to study the putative pathophysiological aspects of preeclampsia. These include placental ischemia (RUPP rat model), angiogenic imbalance (sFLT1 overexpression model), maternal immune activation (AT1-AA model) and genetic models such as BPH/5, STOX1 overexpression etc., (53). To our knowledge, an animal model that exhibits early- and late-onset preeclampsia phenotypes with a common etiology has not been demonstrated. In this study, we report that placenta-specific increase in complement activation leads to early- or late-onset preeclampsia depending on the trajectory of increase.

Precise regulation of complement activation is critical during pregnancy to protect the fetus from immune-mediated injury while maintaining maternal immune surveillance. In rodents, the complement regulatory protein Crry plays a central role in this balance, compensating for the delayed placental expression of decay accelerating factor (DAF) and the tissue-restricted expression of membrane cofactor protein (CD46) (14;15;54). Complete deletion of Crry results in embryonic lethality by GD 9.5 (17), highlighting its indispensable role in early gestation. To circumvent early lethality and assess how varying degrees of placental complement dysregulation affect pregnancy outcomes, we developed a placenta-specific, doxycycline-inducible, Crry targeted shRNA mouse model. Using the Cyp19a-Cre transgene for trophoblast-specific recombination and a Tet-On system to achieve dose-dependent induction post-GD9.5, we achieved effective and placenta-restricted Crry knockdown. Recombination efficiency analysis confirmed approximately 80% placental targeting with minimal activity in fetal tissues validating the specificity of our model. To address the differences in the magnitude of complement activation in different individuals based on their complotype (21;22), we sought to understand how different complement activation levels affect the maternal and fetal outcomes in this mouse model.

Our data revealed that the low- and the high-dose doxycycline administration produced distinct trajectories of complement activation and associated maternal-fetal phenotypes. Immediately after the shRNA was induced, placental Crry protein levels were downregulated in the low- and the high-dose groups. However, in the low-dose group, a compensatory feedback response reduced the effect of shRNA gradually so that towards the end of the gestation, the Crry levels were equal to the control group. On the other hand, the compensatory feedback response was stronger in the high-dose group causing substantial upregulation of Crry protein soon after the shRNA induction followed by significant downregulation towards the end of gestation. As a result, a moderate and sustained level of complement activation occurred in the low-dose group, whereas excess complement activation occurred in the high-dose group only near the end of the gestation.

The maternal consequences of these complement alterations were dose dependent. In the low-dose group, maternal liver weights failed to increase with advancing gestation, indicating impaired hepatic adaptation, a finding not accompanied by overt liver injury as serum AST, ALT, and LDH levels remained normal. Moreover, glomerular endotheliosis was observed in both the low- and the high-dose groups, but only the high-dose animals exhibited proteinuria.

Fetal outcomes also diverged. Fetuses in the low-dose group initially exhibited enhanced growth and placental efficiency at GD13.5, but by GD17.5, they showed fetal growth restriction and reduced placental efficiency. This shift was associated with a specific reduction in the junctional zone and altered expression of key placental hormones—prl3d1 and prl3b1—suggesting impaired endocrine support to the fetus.

Additionally, we found that glycogen storage was significantly reduced in the low-dose placentas at GD17.5, despite normal maternal glucose metabolism, insulin levels, and glucose/insulin tolerance. This suggests a placenta-intrinsic defect in energy storage contributing to fetal growth restriction. Interestingly, expression of lipid transporters and binding proteins such as FATP6, FABP4, and FABP5 were elevated in the low-dose placentas likely as a compensatory mechanism for deficient glycogen reserves. However, these adaptations were insufficient to sustain fetal growth, underscoring the critical role of placental endocrine and metabolic functions in supporting fetal development.

In contrast, the high-dose animals showed preserved fetal growth despite significant late-gestational complement activation and maternal kidney injury, suggesting a more consistent phenotype with late-onset preeclampsia, where placental involvement is limited, and fetal outcomes are relatively spared.

Taken together, our findings provide strong evidence that the magnitude and timing of complement activation in the placenta determines whether maternal, fetal, or both compartments are affected. Moderate and sustained complement activation impairs placental function and fetal growth, while higher, delayed activation preferentially affects maternal organs. This dose-dependent divergence recapitulates key features of early- and late-onset preeclampsia in humans (49;52), providing a unified mechanistic framework for understanding their etiology.

Moreover, the balance observed between shRNA-mediated knockdown and feedback upregulation of Crry illustrates the dynamic regulation of complement homeostasis during pregnancy. However, extrapolating the findings from this study to the human pregnancy is complicated by the absence of Crry in humans and expression of other complement inhibitory proteins such as CD46 and CD55 in human placenta. However, recent studies have indicated that CD55 is upregulated as a feedback response to increased complement activation in different tissues (55–58). On this basis we speculate that identical to the findings in this study, complement activation follows different trajectories depending on the activation and feedback inhibition balance at the human fetal-maternal interface. This dynamic balance is dictated by pregnancy “complotype” (21;22).

This model serves as a valuable experimental platform to explore the pathogenesis of preeclampsia, investigate biomarker dynamics, and test therapeutic strategies aimed at restoring complement balance at the fetal-maternal interface.

## MATERIALS and METHODS

### Animals

To achieve placenta specific complement activation, we used a tet-inducible promoter (TRE) to drive the expression of a miRE-based shRNA targeting Crry in the presence of reverse tet-transactivator (rtTA3). To avoid leakiness of the TRE promoter a loxP-Stop-loxP cassette was used between TRE promoter and GFP-miRE expression. With this approach founder transgenic mice, Cr1l.634shRNA were generated by Mirimus. Male Cr1l.634shRNA mice were then crossed with female B6. Cg-Gt (ROSA)26Sortm1(rtTA*M2) Jae/J to obtain Cr1l^+/+^RosartTA^+/+^ mice. These mice were of C57BL/6 genetic background. To obtain placenta specific shRNA induction, we used Cyp19-Cre mice (Kind gift from Dr. Gustavo Leone) in which Cre is under Cyp19 promoter and hence expressed only in placenta. Female Cyp19-Cre mice were of FVB and C57BL/6 mixed genetic background. Female Cyp19-Cre mice were then mated with male Cr1l^+/+^RosartTA^+/+^ mice. To induce shRNA, doxycycline in drinking water (75 or 35 µg/mL) was provided to pregnant Cyp19-Cre mice from 10.5dpc to 17.5dpc.

### RNA extraction and real-tie quantitative PCR

The total RNA was extracted from frozen placental tissues using Aurum™ Total RNA Fatty and Fibrous Tissue Kit (Bio-Rad, 7326830) following manufacturer’s instructions, and their concentrations were determined using a NanoDrop (Thermo Fisher Scientific) spectrophotometer. cDNA was synthesized using iScript™ Advanced cDNA Synthesis Kit (Bio-Rad, 1725038). Following cDNA synthesis, quantitative real time PCR was performed on CFX Opus 96 thermocycler using iTaq™ Universal SYBR® Green Supermix (Bio-Rad, 1725125) with the primers listed in the following table. Relative mRNA expression was calculated using the 2^-ΔΔCT^ method, normalized to the average of three housekeeping genes, Hsp90ab1, β2-microglobulin, and β-actin. Fold change was calculated to respective control group.

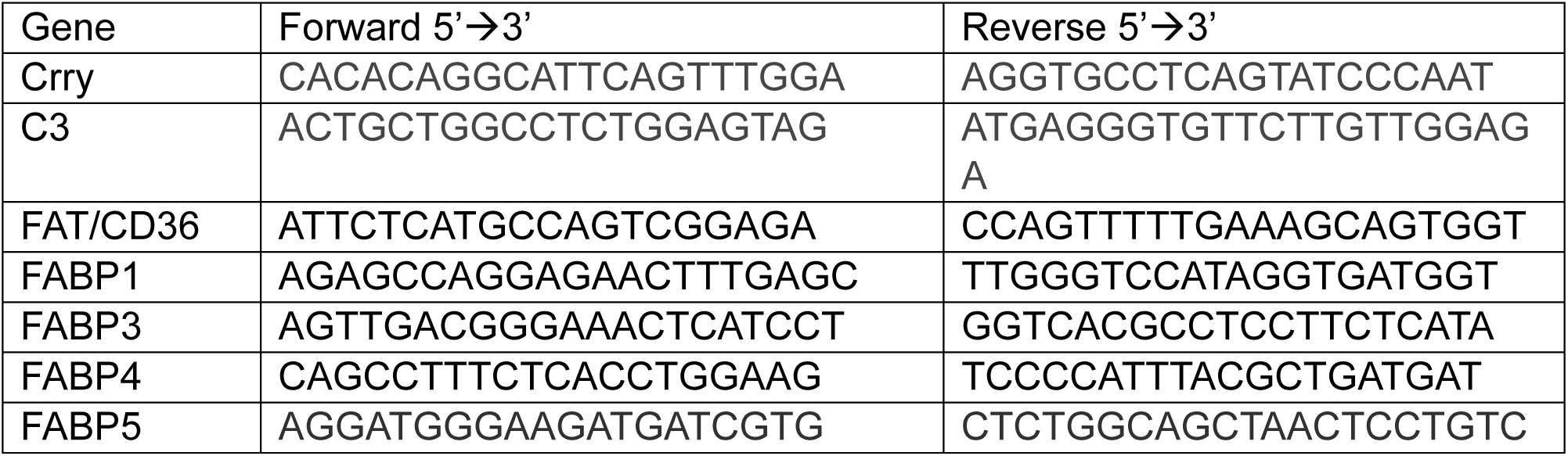
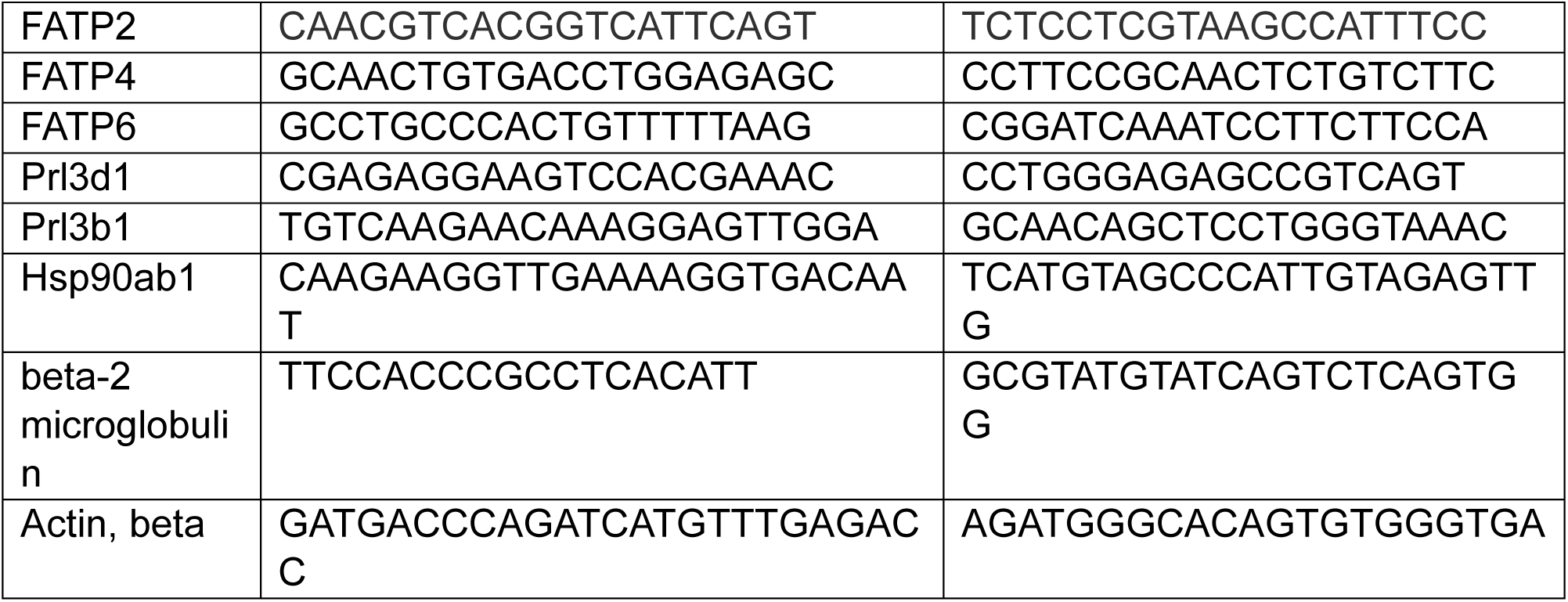

### Protein extraction and Western blot analysis

Protein was extracted from snap frozen placentas using RIPA buffer (Cell signaling, 9806S) supplemented with PMSF (Cell signaling, 8553S), Protease inhibitor cocktail (Selleckchem, B14001), Phosphatase inhibitor cocktail 2 (Sigma, P5726) and Phosphatase inhibitor cocktail 3 (Sigma, P0044). Protein concentrations were determined using Pierce™ BCA Protein Assay Kits (Thermo Scientific, 23225). 20µg of protein extracted or positive protein controls were run on 10% Mini-PROTEAN® TGX Stain-Free™ Protein Gels (Bio-Rad) and transferred onto immune-blot PVDF membrane (Bio-Rad, 1620177). Membranes were blocked in 5% fetal bovine serum (FBS) then incubated in the primary antibodies listed in the following table in 5% FBS overnight at 4°C. Secondary antibodies (1:5000; SouthernBiotech 1030-05 or 4050-05; anti-mouse or anti-rabbit IgG, HRP linked) were added for 1.5 hours. After washing, Clarity Western ECL substrate (Bio-Rad, 1705061) was added to the membranes and developed in Odyssey FC Imaging System (LI-COR). Image Studio Software was used to quantitate the density of the bands. β-actin was used as protein loading control, and samples were normalized to control groups in each blot.

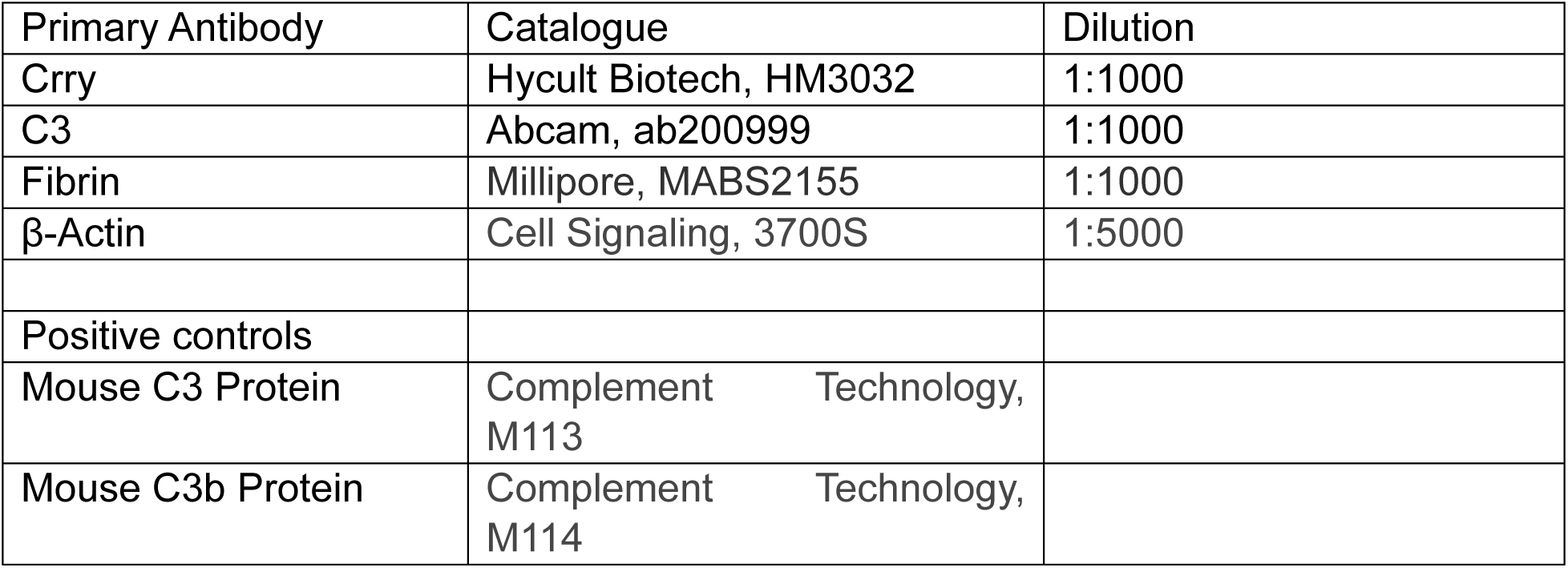
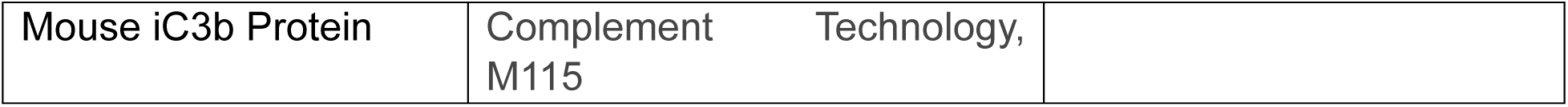

### Serum C3 concentration measurement

Serum C3 levels were measured using Mouse Complement C3 ELISA Kit (Abcam, ab157711) according to manufacturer’s instructions.

### Maternal preeclampsia phenotype

Systolic, diastolic mean arterial blood pressures were measured by tail cuff method in non-anesthetized mice using 8-channel CODA system (Kent Scientific Corporation). During BP measurements mice were restrained in the nose cone holders with unrestricted breathing and clear visibility. An occlusion tail cuff and volume pressure recording (VPR) sensor were gently inserted onto the tails of mice placed on a pre-warmed (37^0^C) platform. Recordings were performed in three sessions with 20 cycles in each cycle. The consistent readings from the three sessions were averaged to get the final reading.

### Uterine artery doppler study

Doppler ultrasounds were performed at the Small Animal Imaging Core Facility at Baylor College of Medicine on pregnant mice under room air using a 30-MHz pulsed Doppler system (Vevo 2100, VisualSonics) equipped with an MX550D probe. The uterine artery was identified at the anatomical location where it crosses over the external iliac artery. Mice were anesthetized with 1.0–2.0% isoflurane (Baxter Healthcare Corporation) delivered with 1.5 L/min oxygen. Animals were placed supine on a heated imaging platform (Vevo 2100 Imaging Station, VisualSonics) and secured gently with adhesive tape. Abdominal hair was removed using depilatory gel to optimize acoustic coupling. Throughout the procedure, body temperature was maintained at 37°C, and heart rate was continuously monitored to ensure animals remained within safe physiological limits. Uterine artery blood flow was assessed using pulsed-wave Doppler. The MX550D probe was positioned over the lower abdomen with ultrasound coupling gel, as previously described (59;60). Doppler recordings were acquired from both right and left uterine arteries, near the bladder or fetuses. After acquisition, mice were removed from anesthesia and returned to their home cages for recovery. From each Doppler waveform, peak systolic velocity (PSV), end-diastolic velocity (EDV), and mean velocity were measured. Average PSV and EDV were calculated per mouse, and the following vascular index was derived, Uterine Artery Resistive Index (UARI) = (PSV − EDV) / PSV.

### Immunofluorescence imaging (IF)

After euthanasia tissues were embedded in OCT compound (Fisher Scientific, 4585), snap frozen immediately and stored at −80^0^C until used for IF. Tissue sections were cut at 8µm and fixed with 4% paraformaldehyde. Sections were permeabilized with 0.25% Triton X-100 for 10 minutes, blocked in 1% BSA and stained overnight at 4°C with respective primary antibodies listed in the following table. Secondary antibodies and EverBrite mounting media with DAPI (Biotium, 23002) were added to visualize immuno-staining and nuclei respectively, in tissue sections. Image acquisition was performed with an Olympus IX73 microscope and analyzed by using cellSens software (Olympus Scientific, Walthan MA, USA).

| Primary Antibody | Catalogue | Dilution |
| --- | --- | --- |
| Fibrin | Millipore, MABS2155 | 1:200 |
| eNOS/NOS Type III | BD Biosciences, 610297 | 1:200 |
| Secondary Antibody |  |  |
| Goat anti-Mouse, Alexa Fluor 594 | Invitrogen, A-11005 |  |

### Hematoxylin & eosin (H&E) staining and Periodic Acid-Schiff staining (PAS)

Paraformaldehyde-fixed tissues were processed, embedded, and sectioned by the Human Tissue Acquisition and Pathology (HTAP) core at Baylor College of Medicine. H&E staining was performed by the HTAP core at Baylor College of Medicine. Periodic Acid-Schiff staining was performed using Periodic Acid-Schiff (PAS) Kit (Epredia, 87007) following the manufacturer’s instructions. Imaging was performed by the RNA In Situ Hybridization Core at Baylor College of Medicine. Placental zone areas and PAS staining area analysis were performed using Image J software.

### Proteinuria measurement

Urine was collected at GD17.5. Urine albumin and creatinine were measured using Mouse Albumin ELISA Kit (Abcam, ab108792, dilution 1:1000) and Creatinine Assay Kit (Colorimetric) (Abcam, ab204537, dilution 1:10) according to manufacturer’s instructions.

### Glucose Tolerance Test (GTT)

Pregnant dams were fasted for 4 hours on GD15.5, and then a baseline fasting blood glucose was obtained from the venous tail sample. An intraperitoneal injection of glucose (1mg/g) was given. Blood glucose levels were obtained at 15, 30, 60, 120 minutes post injection by removing the tail scab for blood collection. All blood glucose measurements were performed in duplicates using two AimStrip Plus Blood Glucose Meters (Germaine™ Laboratories, 37321).

### Insulin Tolerance Test (ITT)

Pregnant dams were fasted for 4 hours on GD15.5, and then a baseline fasting blood glucose was obtained from a venous tail sample. An intraperitoneal injection of insulin (0.5U/kg) was given, as previously described (61). Blood glucose levels were obtained at 15, 30, 45, 60, 90, 120 minutes post injection by removing the tail scab for blood collection. All blood glucose measurements were performed in duplicates using two AimStrip Plus Blood Glucose Meters (Germaine™ Laboratories, 37321).

### Fasting serum insulin and leptin levels

Pregnant dams were fasted for a duration of 4 hours on GD13.5 or GD17.5. Serum was collected after euthanasia. Serum leptin and insulin were measured using a Mouse Leptin ELISA (EMD Millipore, EZML-82K) and a Rat/Mouse Insulin ELISA (EMD Millipore, EZRMI-13K) according to the manufacturer’s instructions.

### Fasting Serum Triglyceride

Pregnant dams were fasted for a duration of 4 hours on GD17.5. Serum was collected after euthanasia. Triglycerides were measured using Triglyceride Colorimetric Assay Kit (Cayman Chemical, 10010303) according to manufacturer’s instructions.

### Ex vivo Lipolysis assay

Pregnant dams were fasted for a duration of 4 hours on GD17.5. Visceral fat was collected after euthanasia. In a 24-well plate, 30mg of fat was added in triplicate wells for control and stimulation media. Control media was composed of 2% BSA (Sigma-Aldrich, A1595) with vehicle control in serum free DMEM media (Corning, 10-013-CV) with 1% Antibiotic-Antimycotic Solution (Corning, 30-004-CI). Stimulation media had the addition of 10uM Isoproterenol. Exactly 500uL of medium was added at the start of the experiment. Exactly 200uL of medium was collected and replaced with 200uL of respective fresh control or stimulation media at time t=0, 30min, 1hour, and 2hours. Glycerol levels were measured using Free Glycerol Reagent (Sigma-Aldrich, F6428) according to manufacturer’s instructions. Calculations were performed as previously described (62).

### Genotyping

DNA Extraction Solution for EZ Fast PCR Genotyping Kit (EZ Bioresearch LLC, G1001) was used to extract DNA from fetal tissues.

Primers for Crry^LSL-RNAi^ were Cola1a Forward 5’-AATCATCCCAGGTGCACAGCATTGCG −3’; Col1a1 Reverse 5’-CTTTGAGGGCTCATGAACCTCCCAGG −3’; and SAdPa Reverse 5’-AAGACCGCGAAGAGTTTGTC −3’. PCR was run using DreamTaq PCR Master Mixes (2X) (Thermo Scientific, K1082) on T100 Thermal Cycler (Bio-Rad). Homozygous Crry^LSL-RNAi^ produces a single band at 353bp, meanwhile homozygous negative (wild type) produces a single band at 245bp. Heterozygous produces a band at 353bp and a band at 245bp.

Primers for rtTA mice are oIMR8052 5’-GCGAAGAGTTTGTCCTCAACC −3’; oIMR8545 5’-AAAGTCGCTCTGAGTTGTTAT −3’; and oIMR8546 5’-GGAGCGGGAGAAATGGATATG −3’. PCR was run using KAPA2G Fast HotStart (Roche Diagnostics, KK5503) on T100 Thermal Cycler (Bio-Rad). Homozygous rtTA^+/+^ mice produces a single band at ∼300bp. Homozygous negative (wild type) produces a single band at 603bp. Heterozygous produces both bands.

Primers for Cyp19-Cre are PW51 5’-GACCTTGCTGAGATTAGATC −3’ and PW22 5’-GACGATGAAGCATGTTTAGCTGGCC −3’. PCR was run using DreamTaq PCR Master Mixes (2X) (Thermo Scientific, K1082) on T100 Thermal Cycler (Bio-Rad). Positive band appeared at ∼545bp. Genotype only showed if it was present but couldn’t distinguish homozygous versus heterozygous. Cyp19-Cre mice were crossed with up to 3 wild type (-/-) mice, and their litter was tested to determine if parent was homozygous (+/+) or heterozygous (+/-).

### Recombination efficiency test

Genomic DNA was extracted from fetal tail, placenta, and offspring tissues using QIAamp DNA Mini Kit (Qiagen, 51304). Genomic DNA (100ng) was amplified using PrimeSTAR GXL Premix (Takara Bio, R051A) with the following primers: GM119 5’-GGGACCGATCCGTCGAGC −3’; GM43 5’-CTTGCCGGTGGTGCAGATGA −3’; and GM115 5’-CATGGTGAGCAAGGGCGAG −3’.

GM119 and GM43 produce a 250bp product when recombination occurs. GM115 and GM43 are used as control that the shRNA cassette is present producing a 160bp product.

---

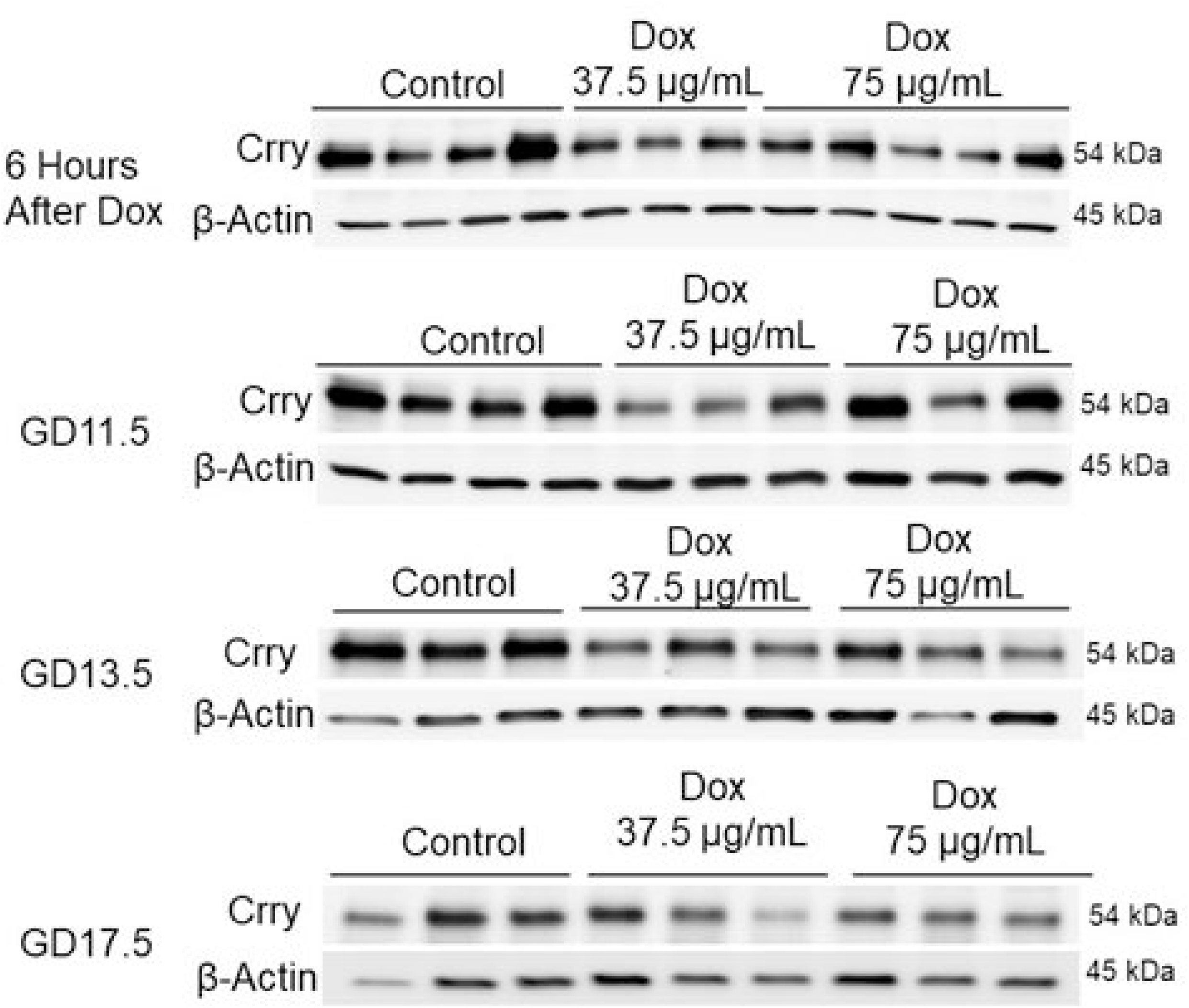
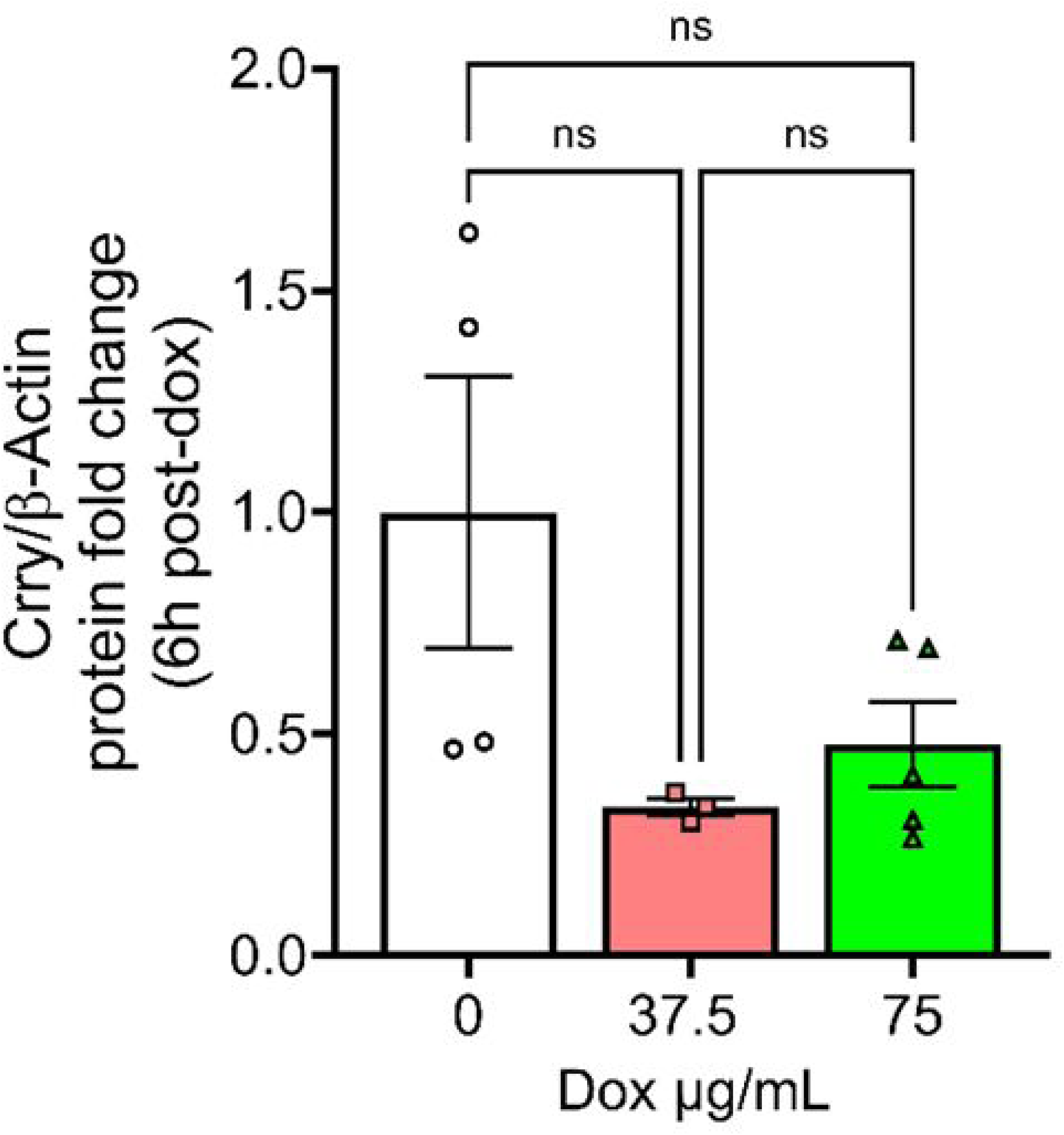
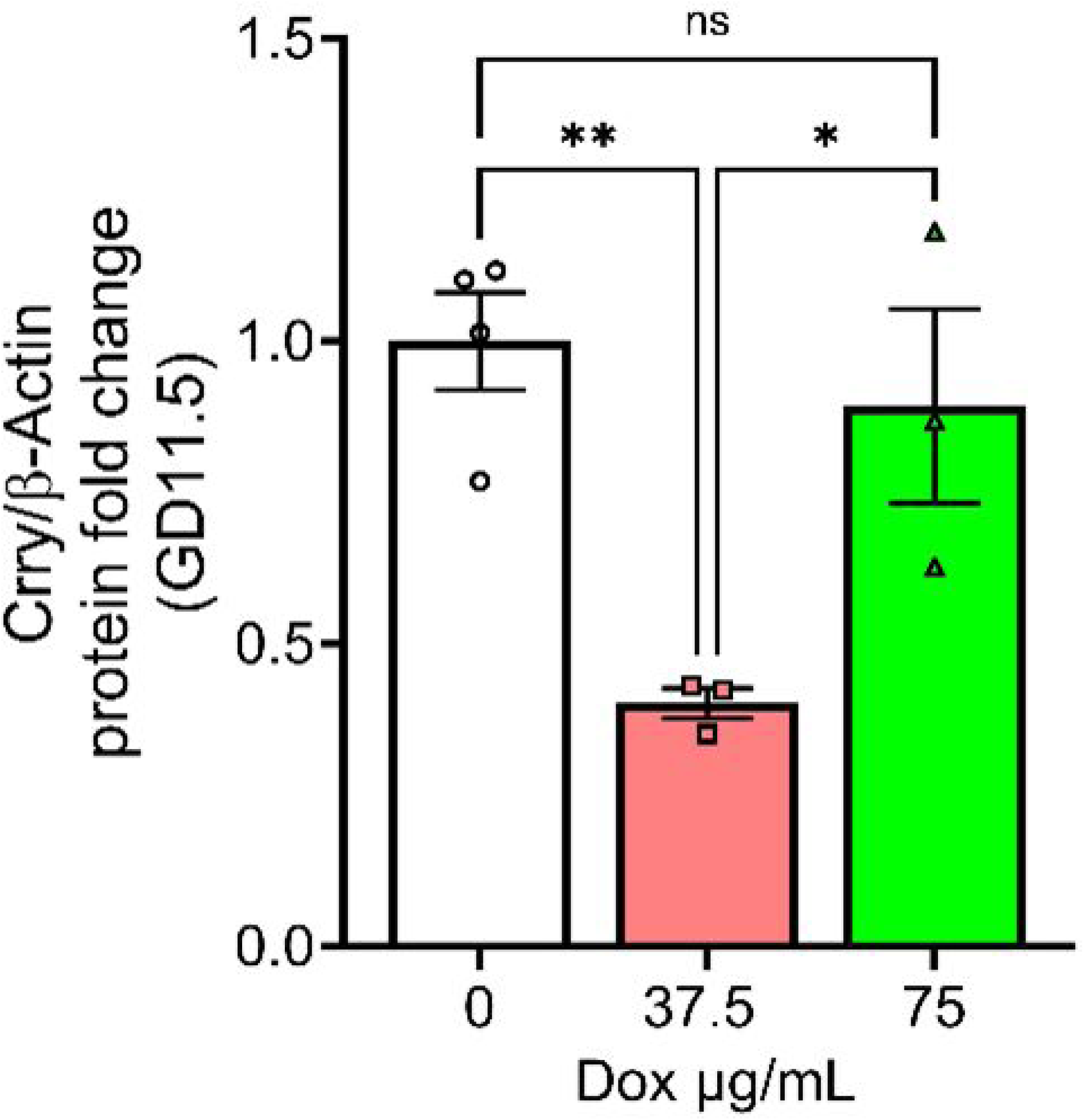
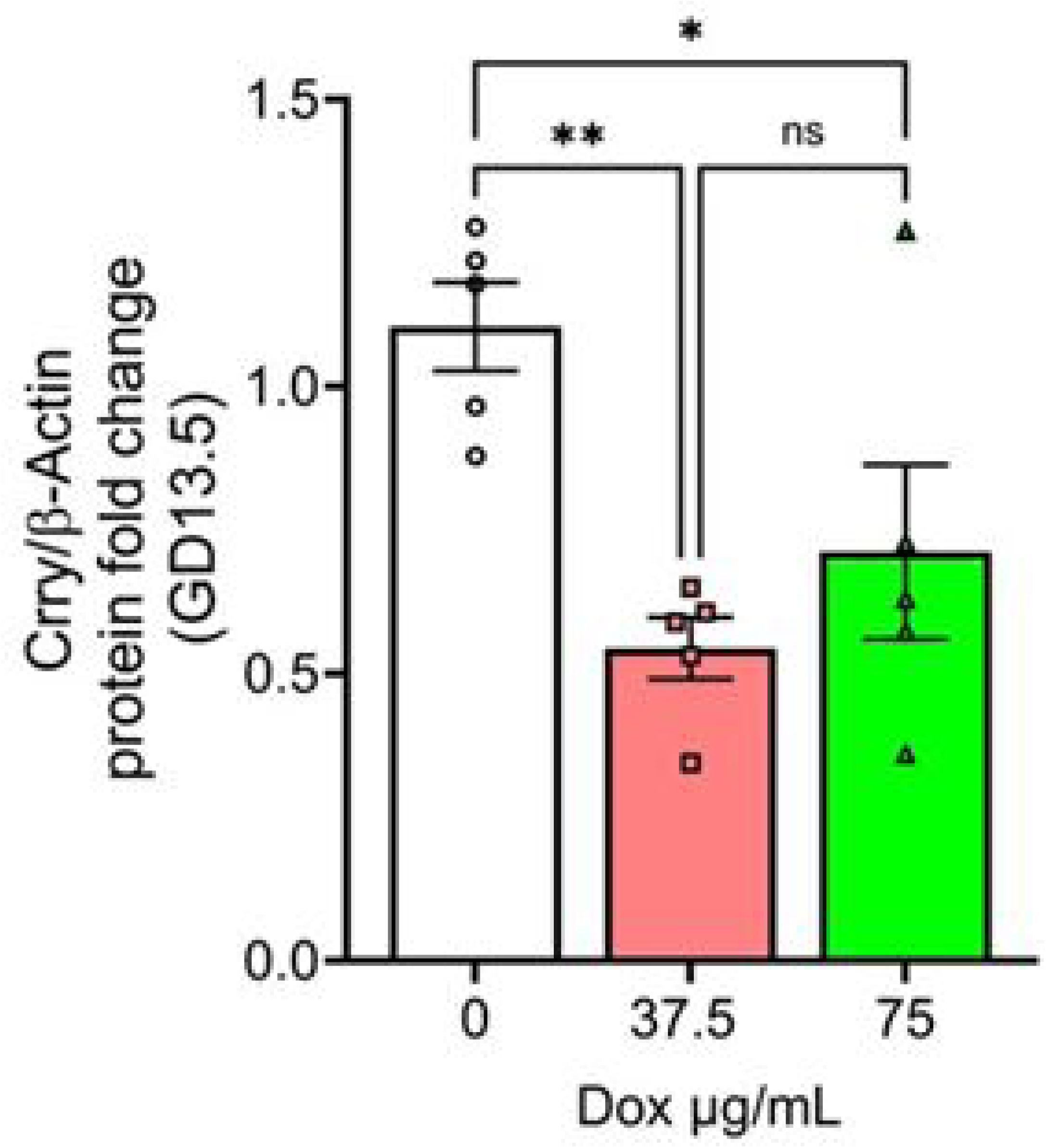
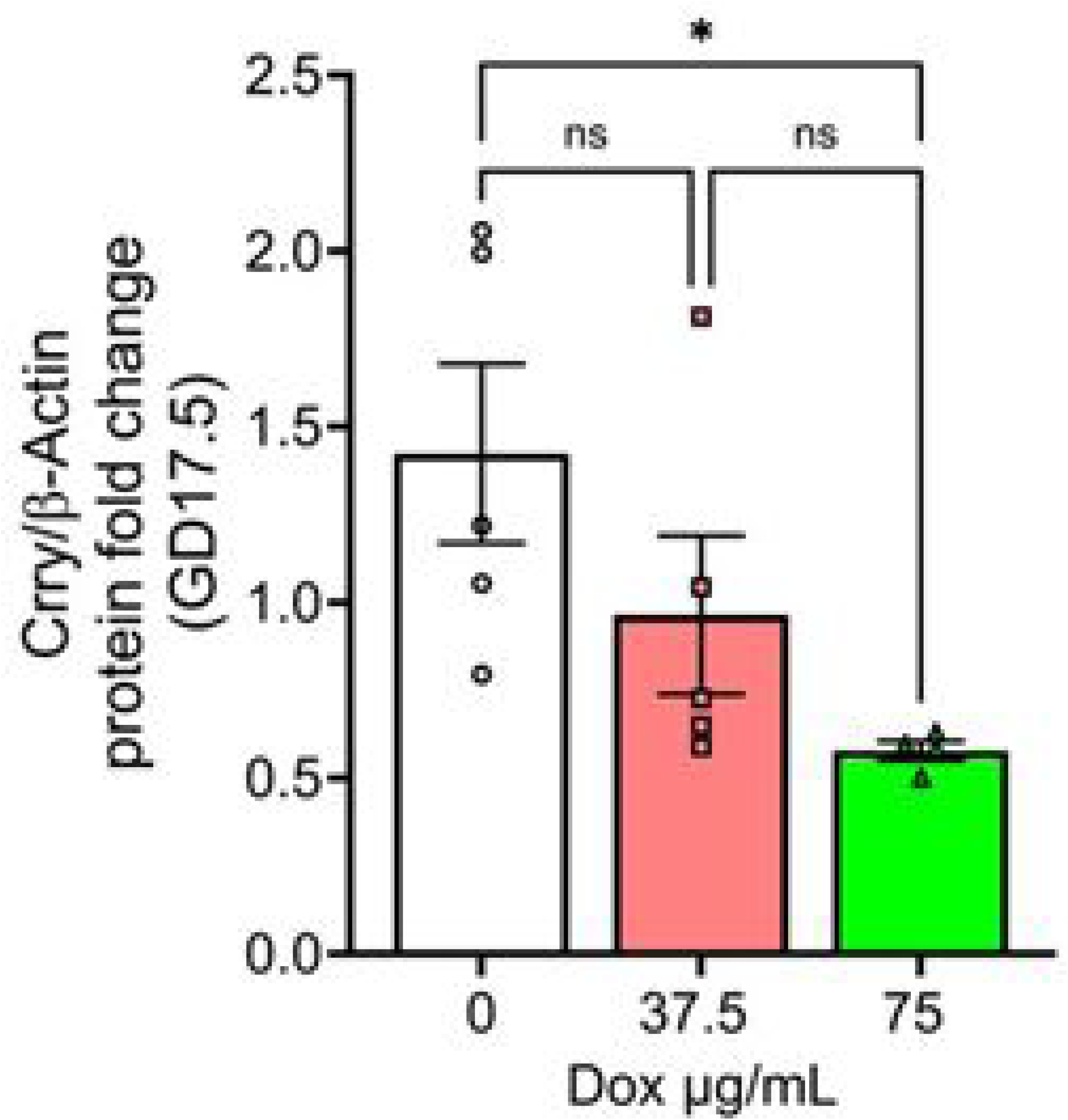
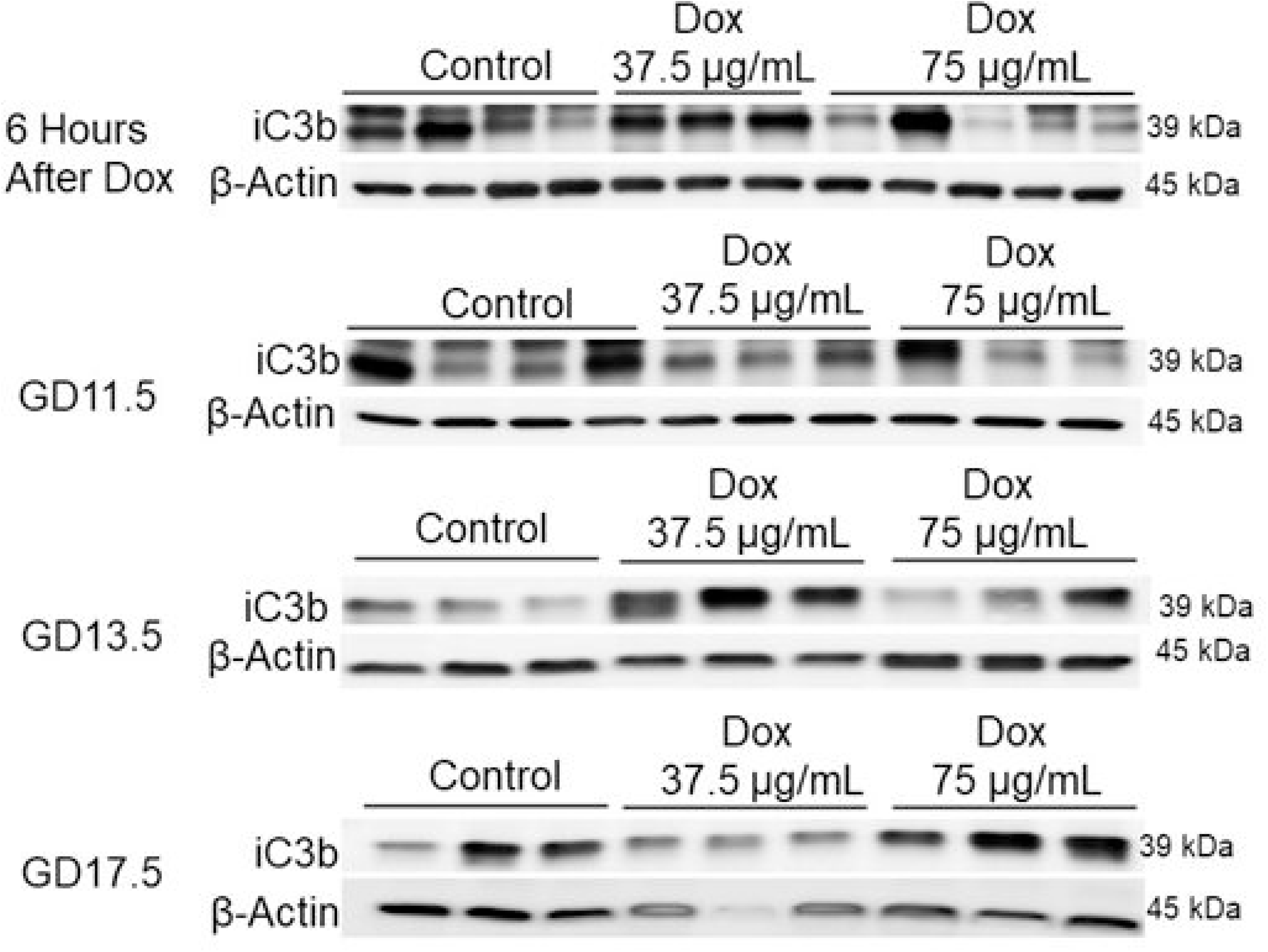
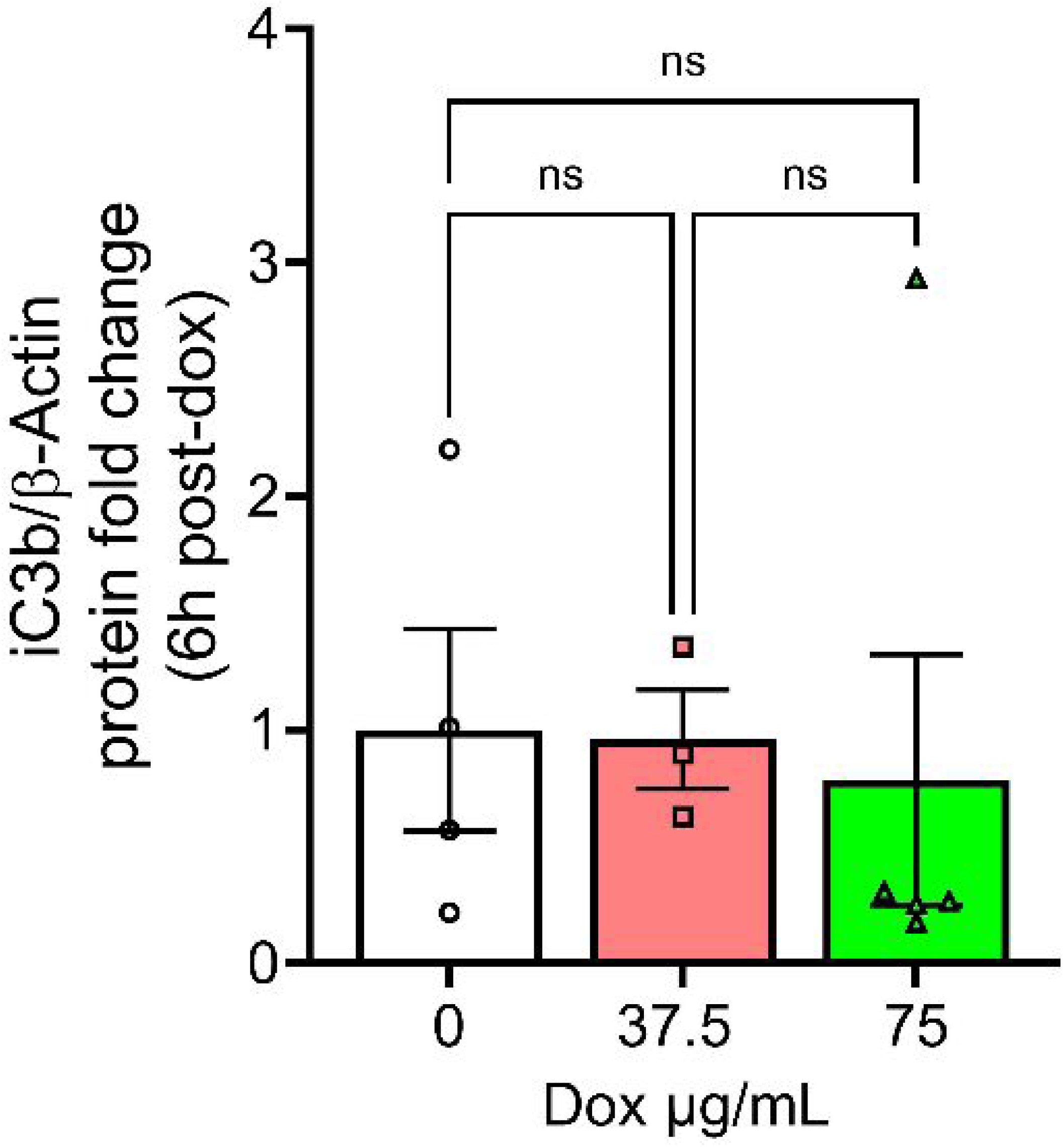
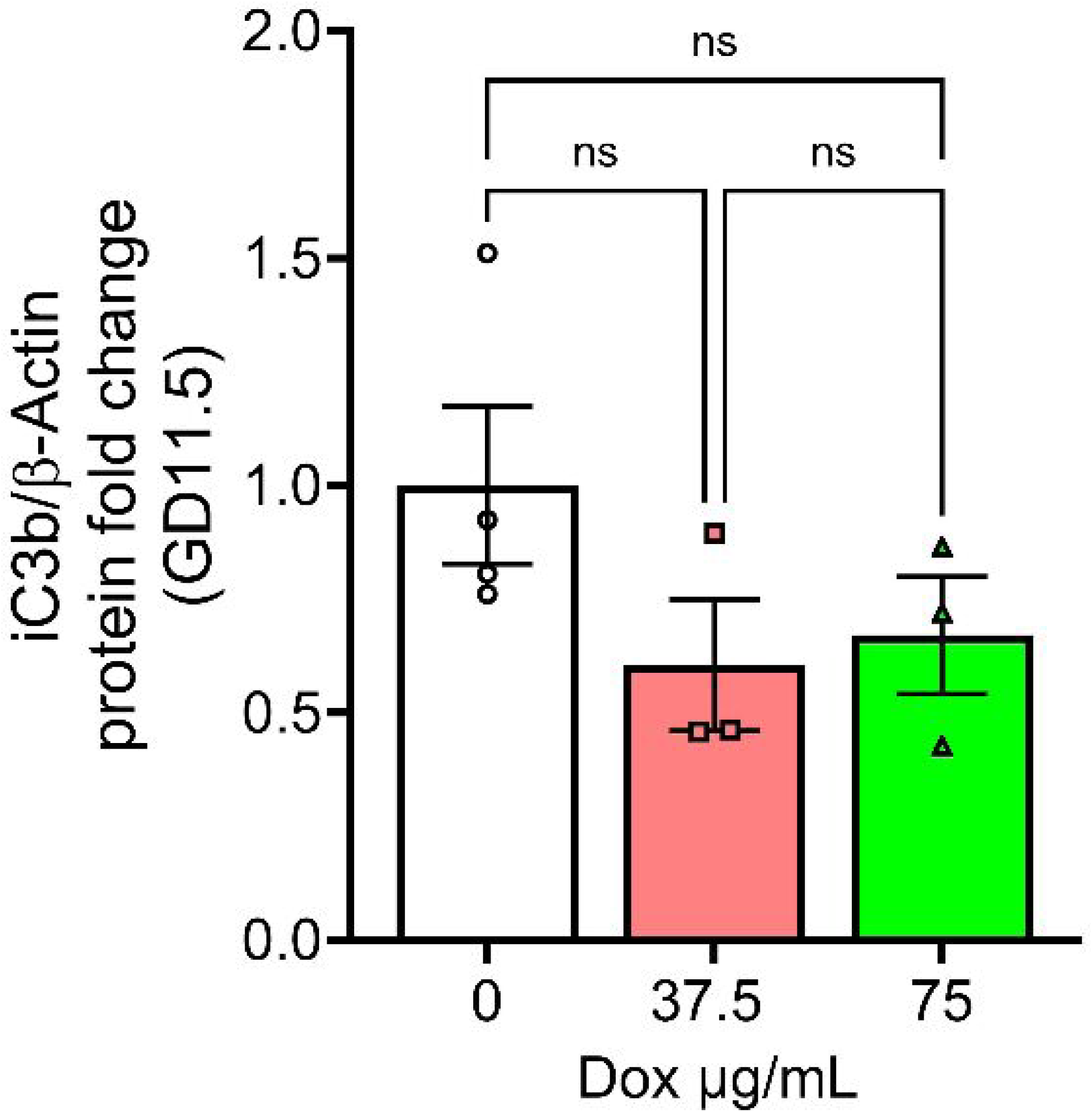
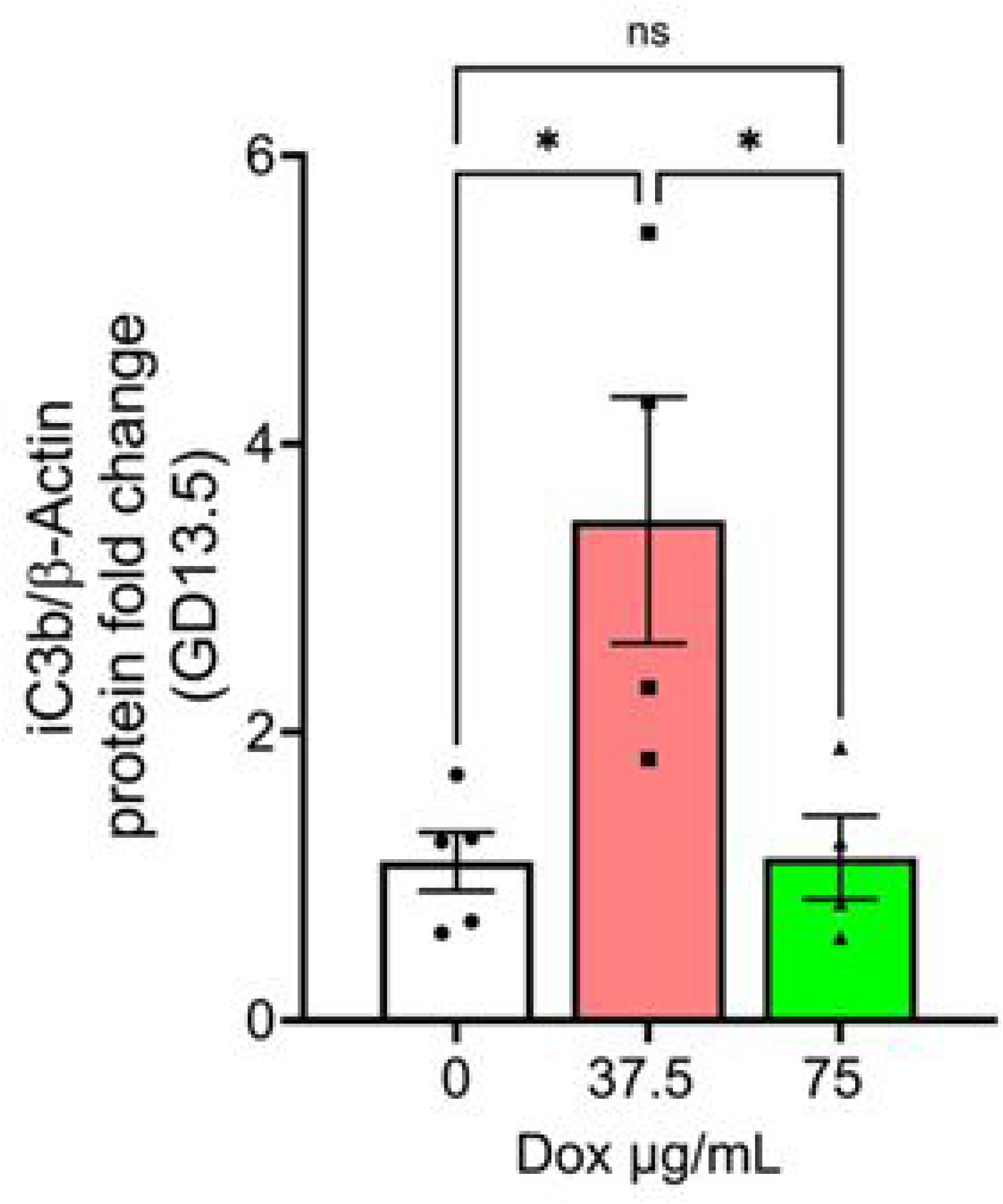
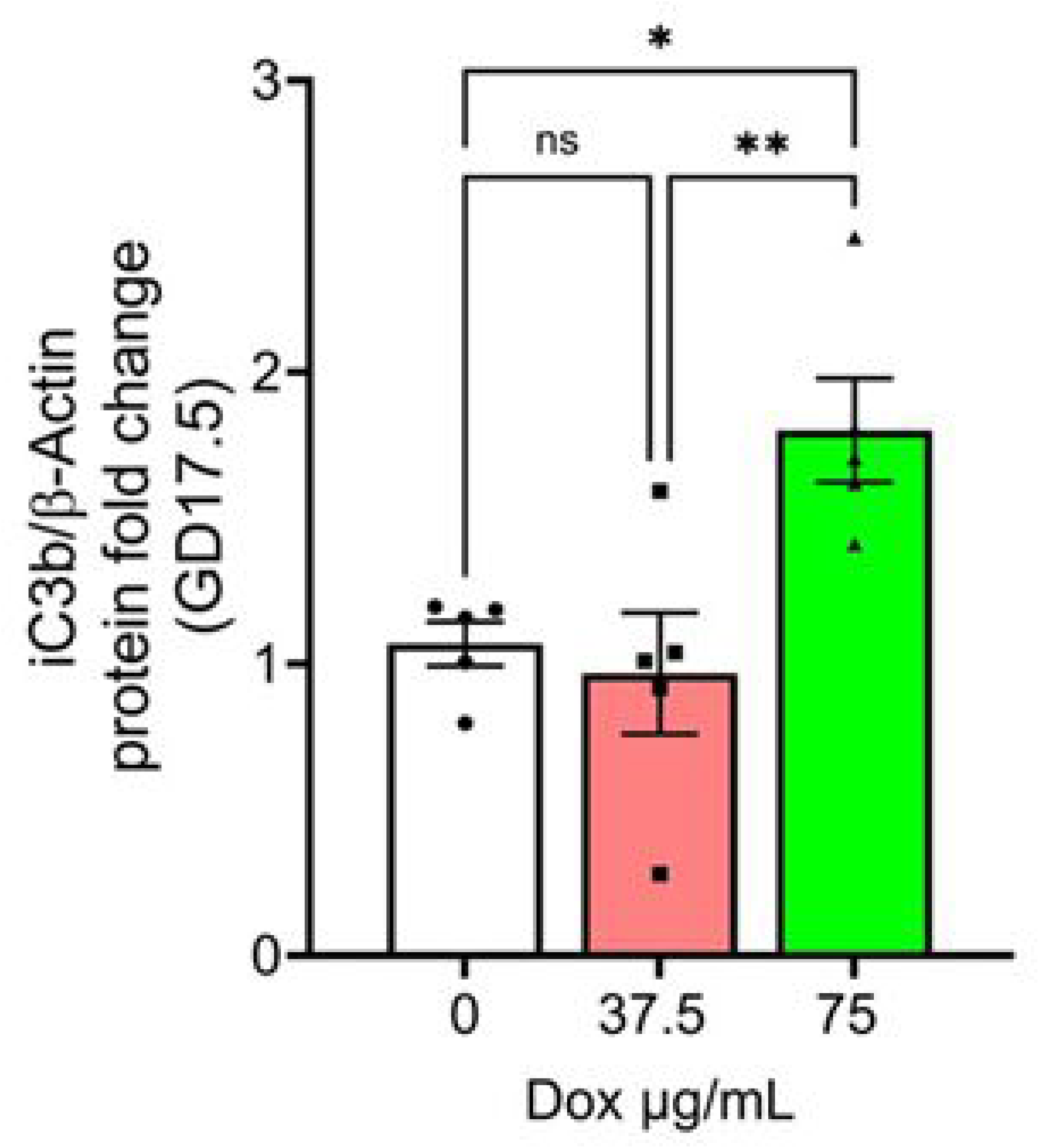
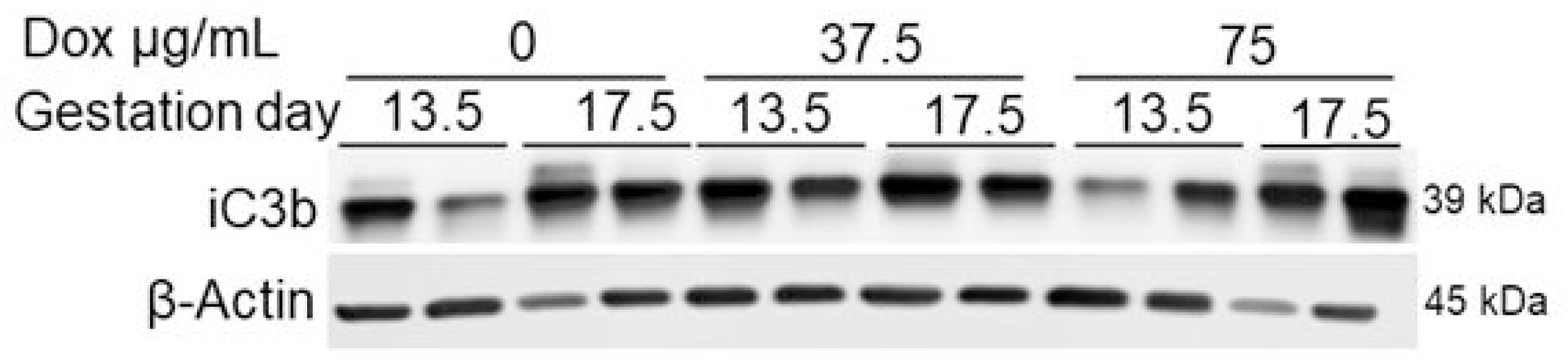
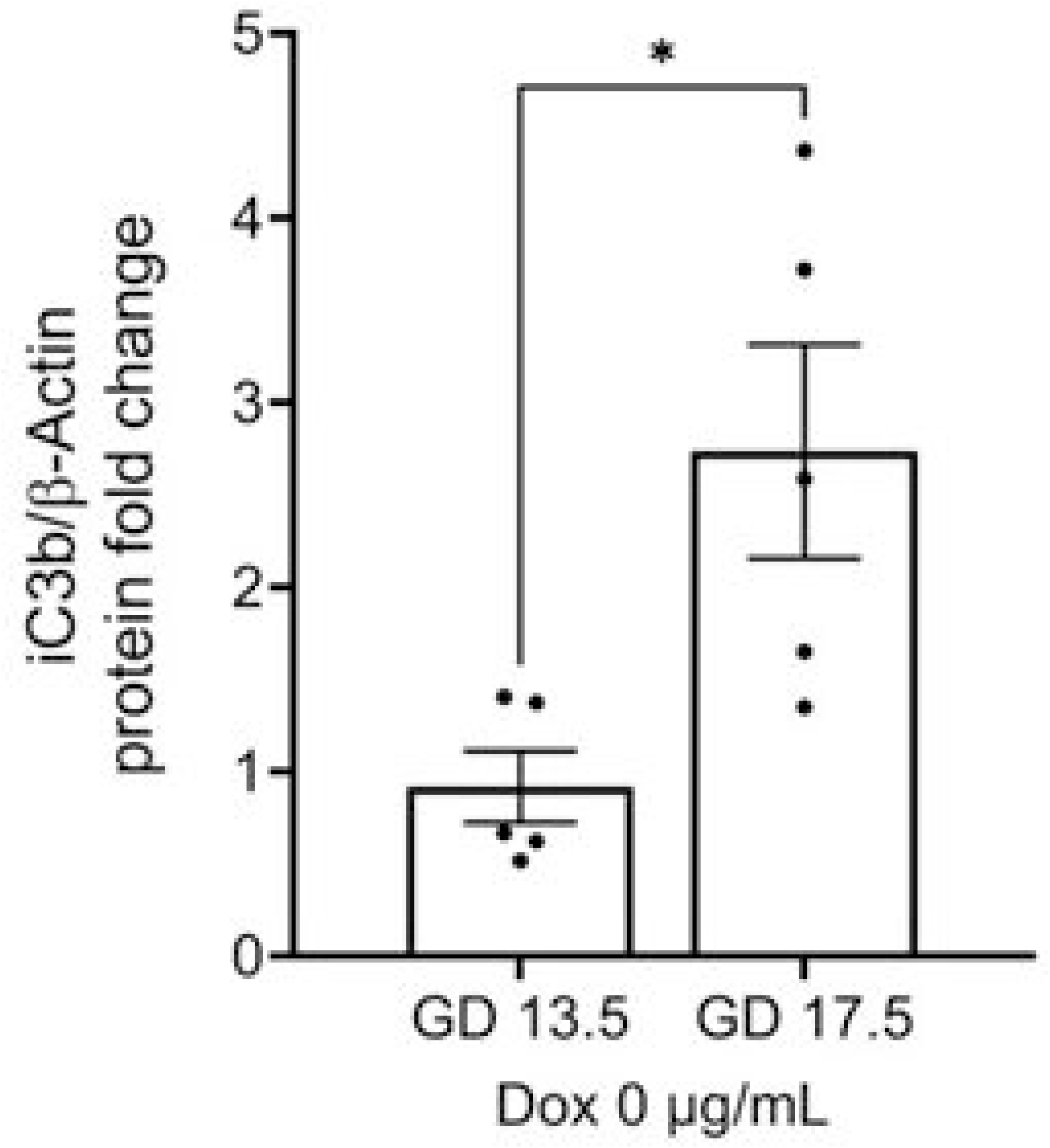
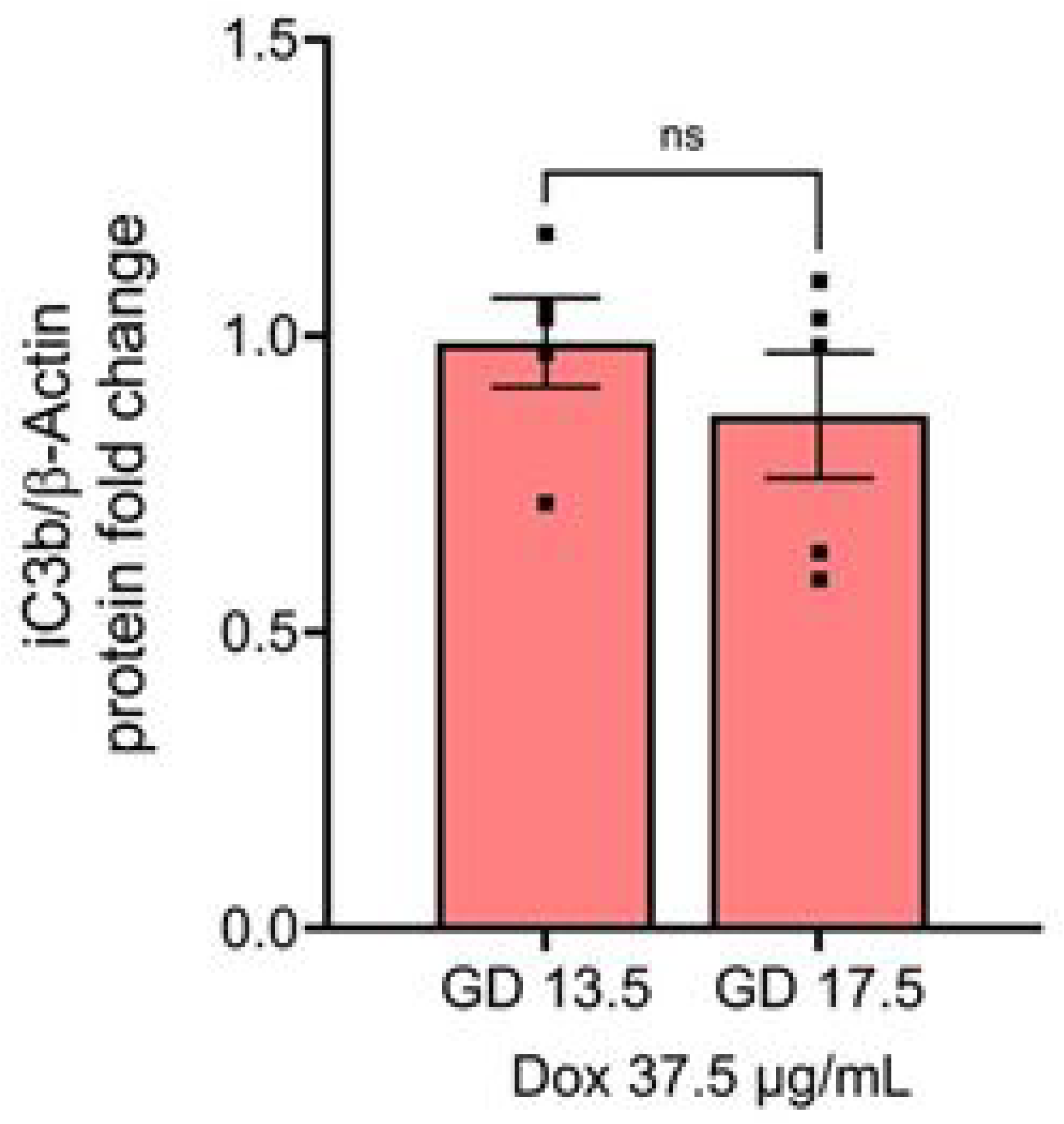
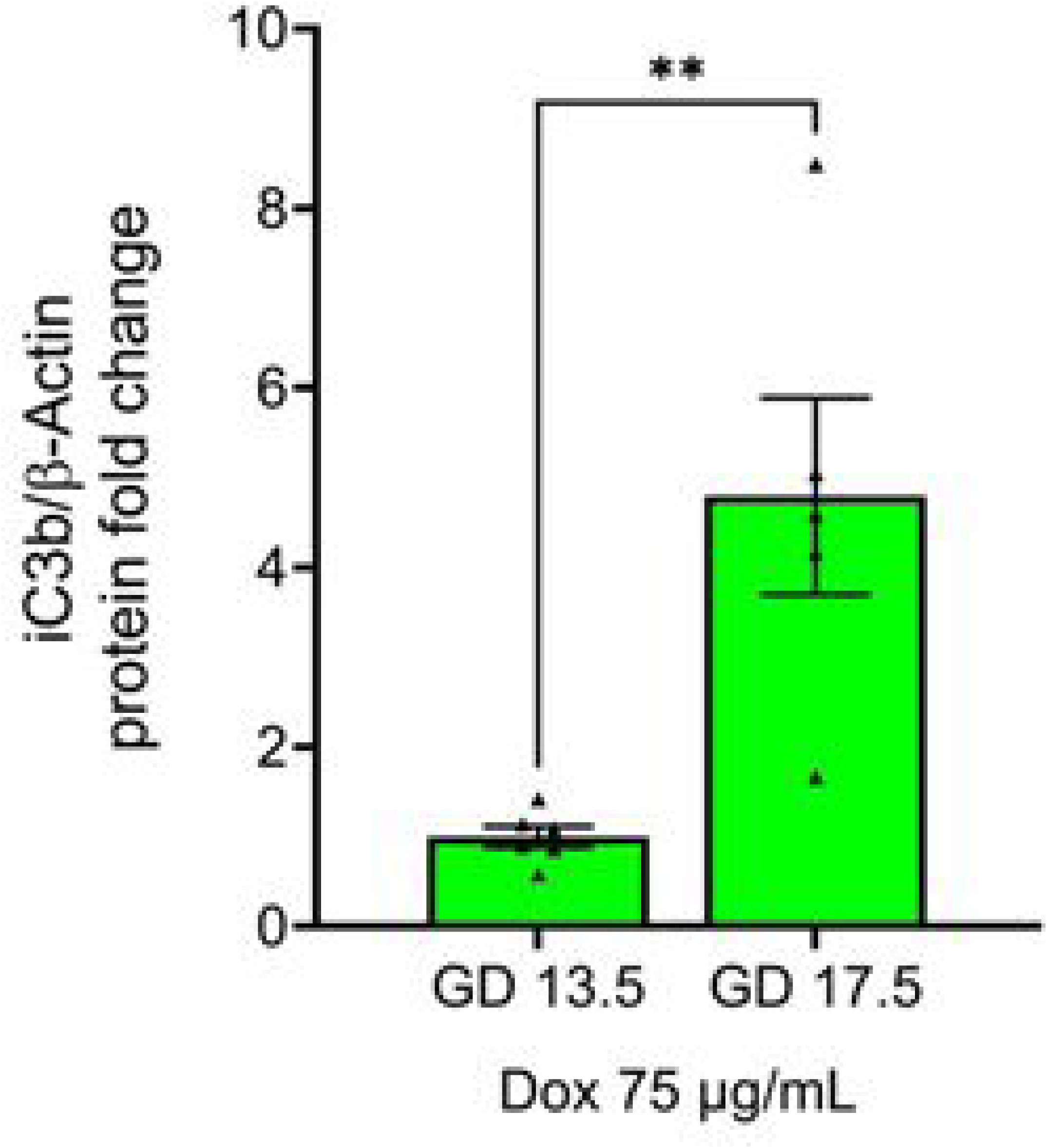
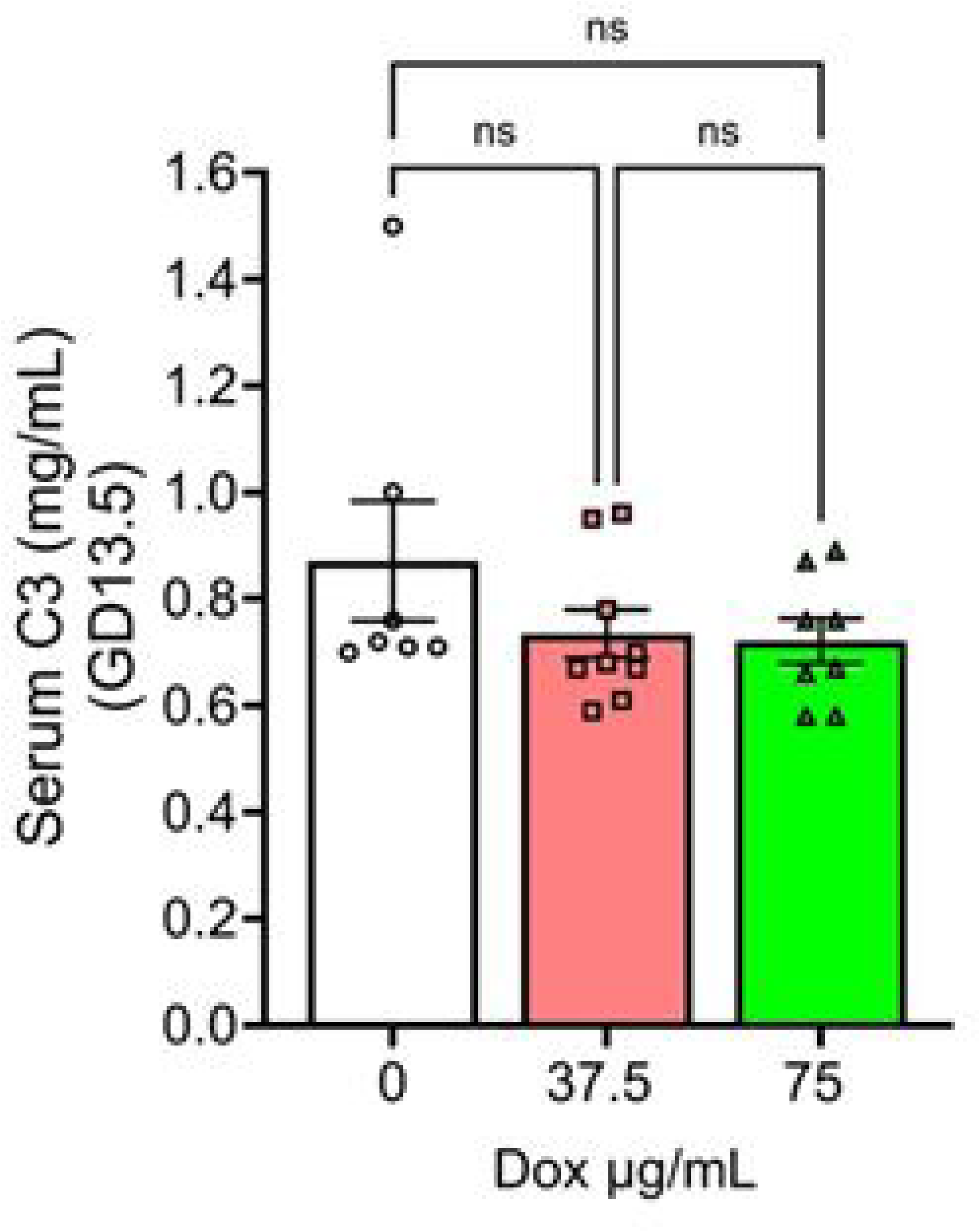
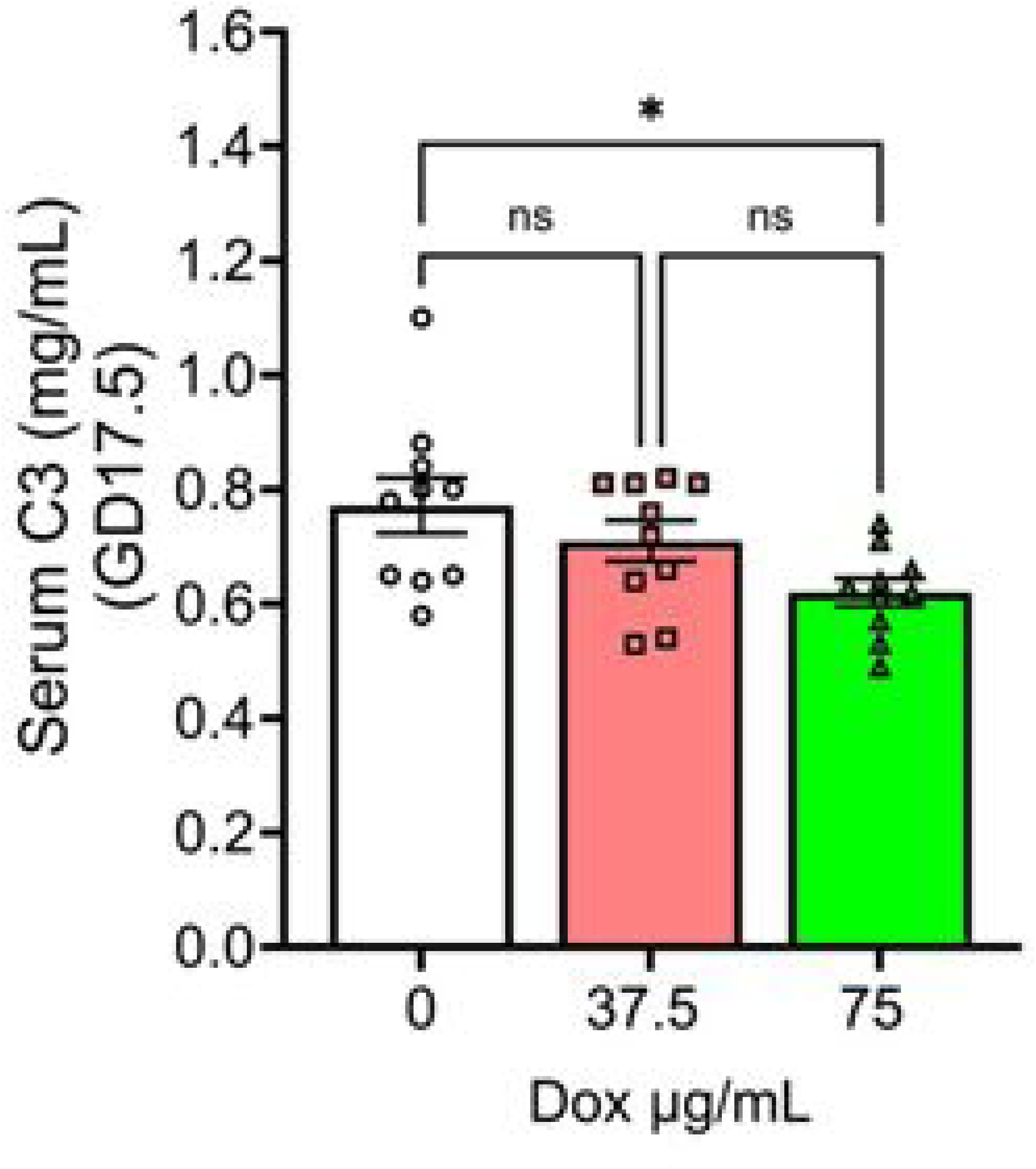
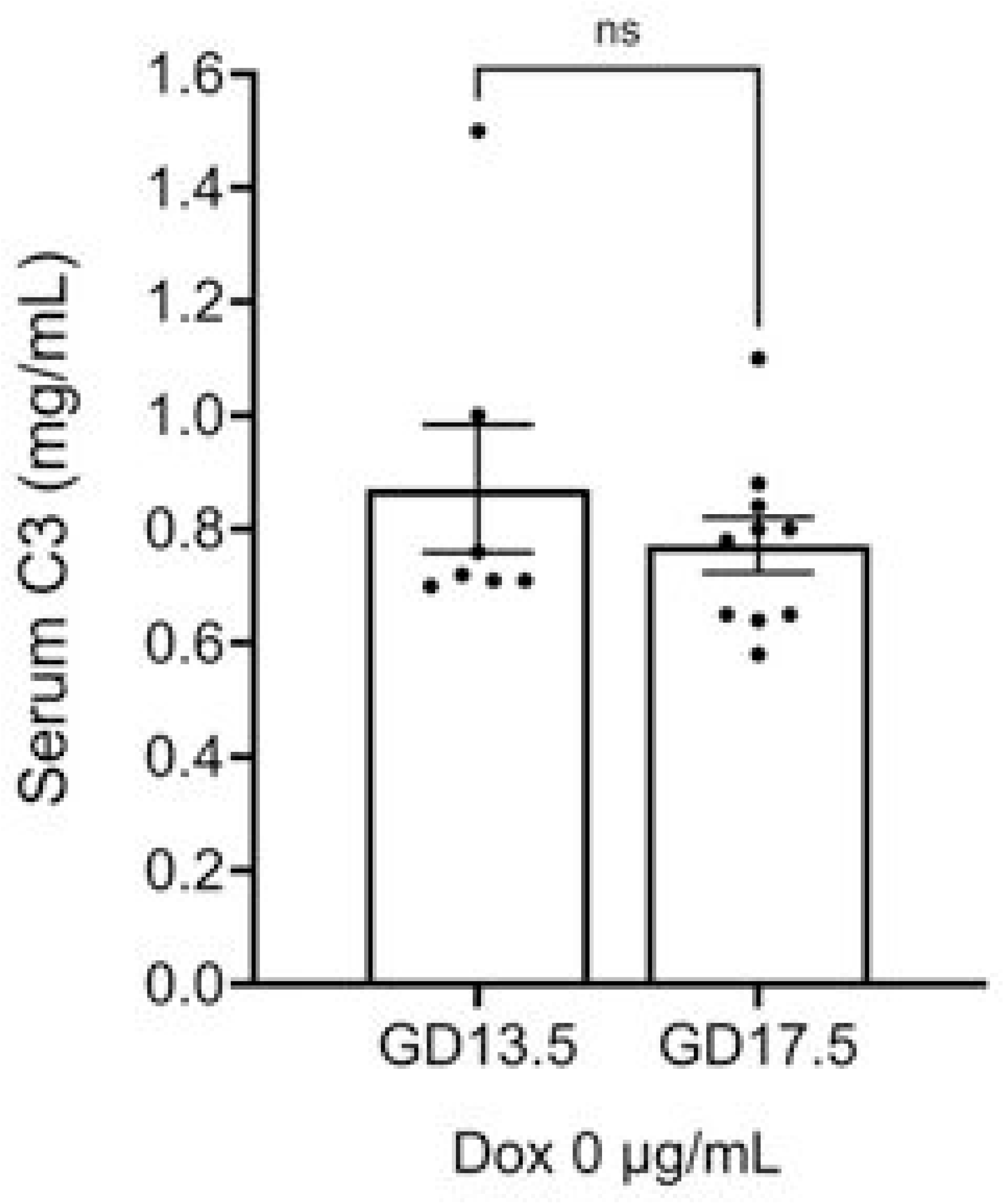
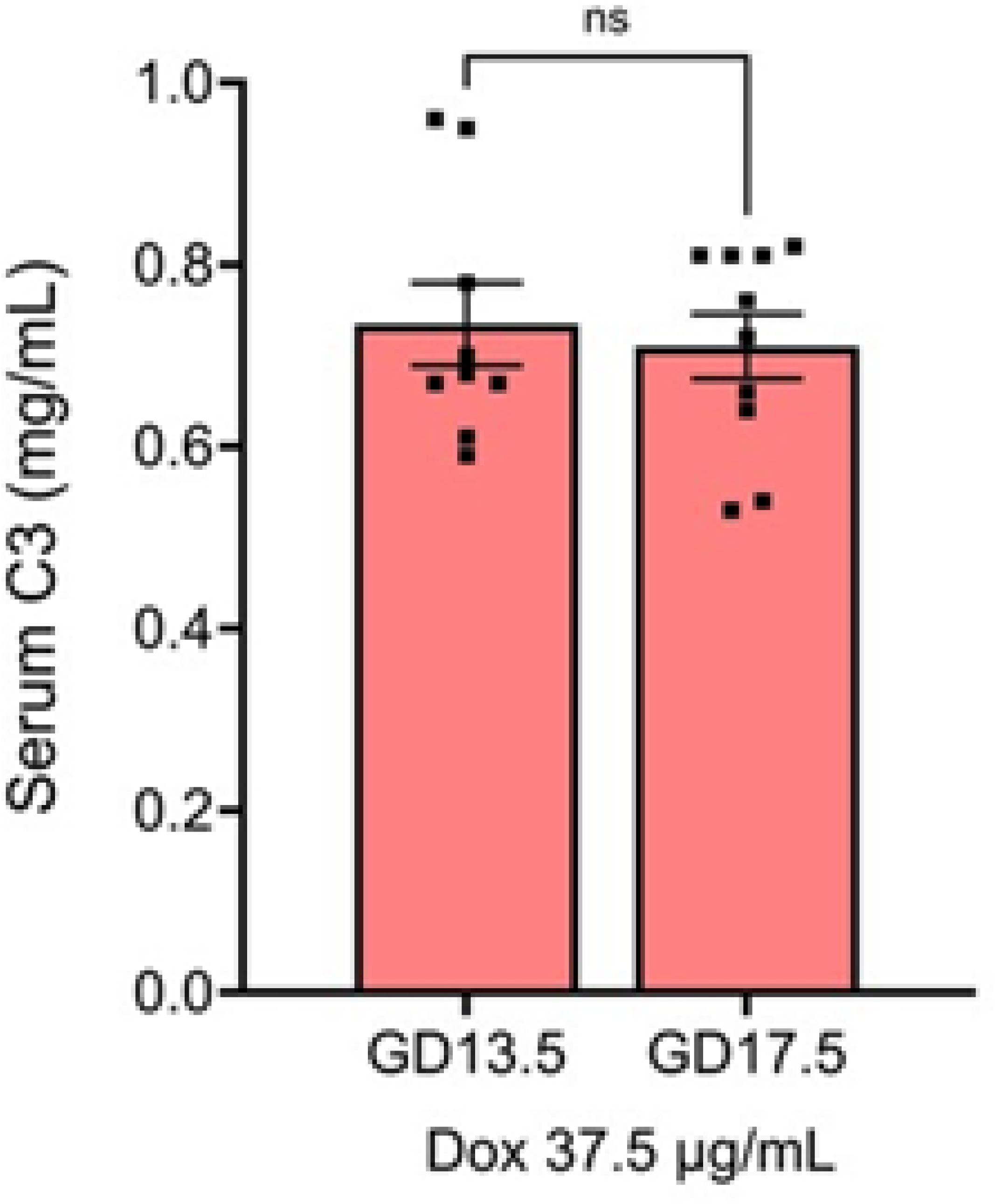
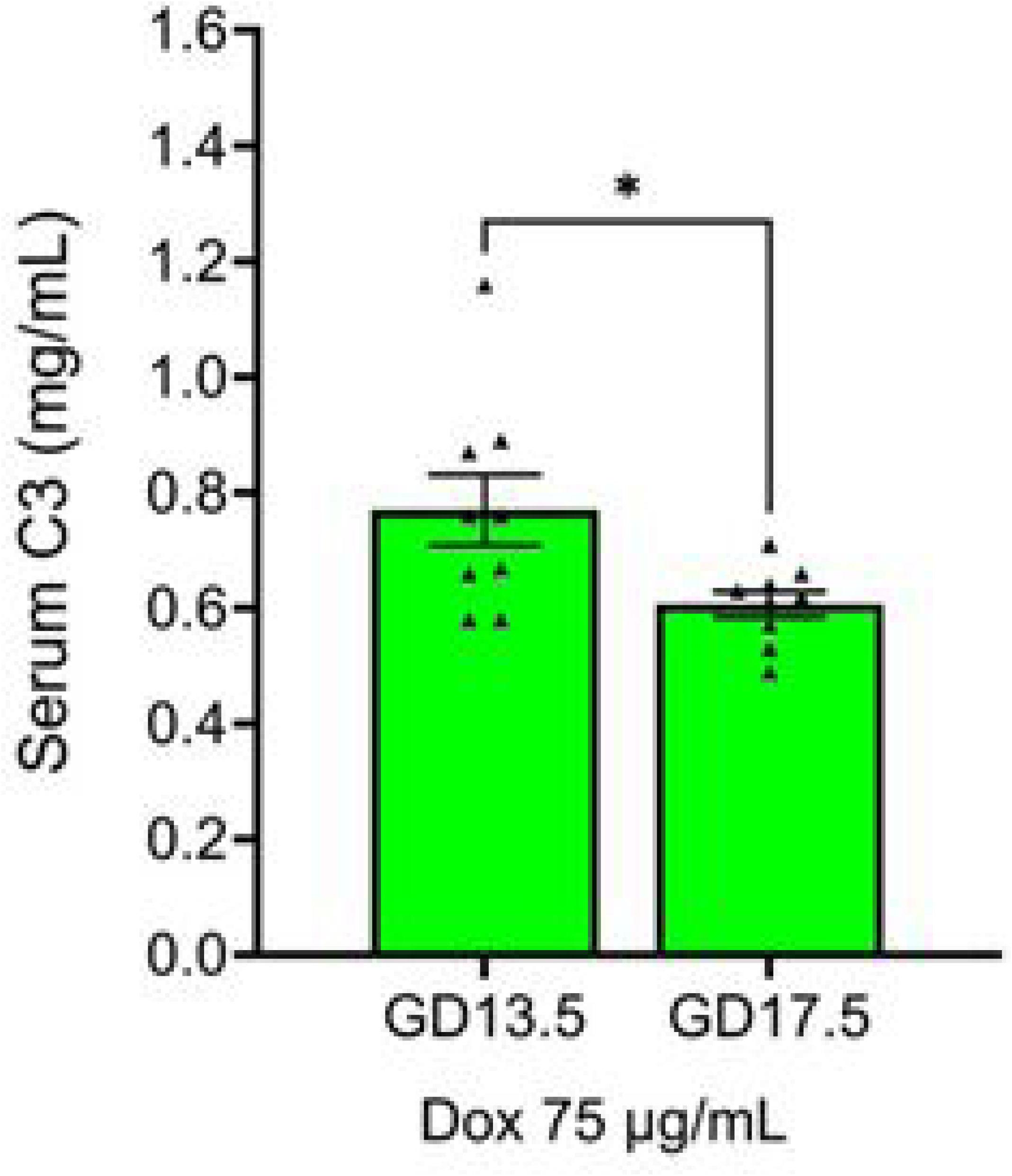
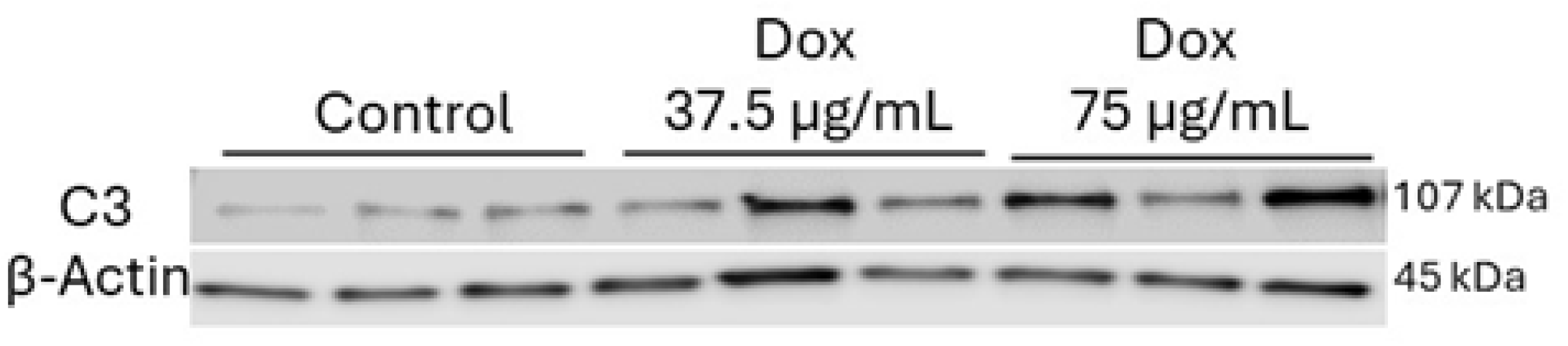
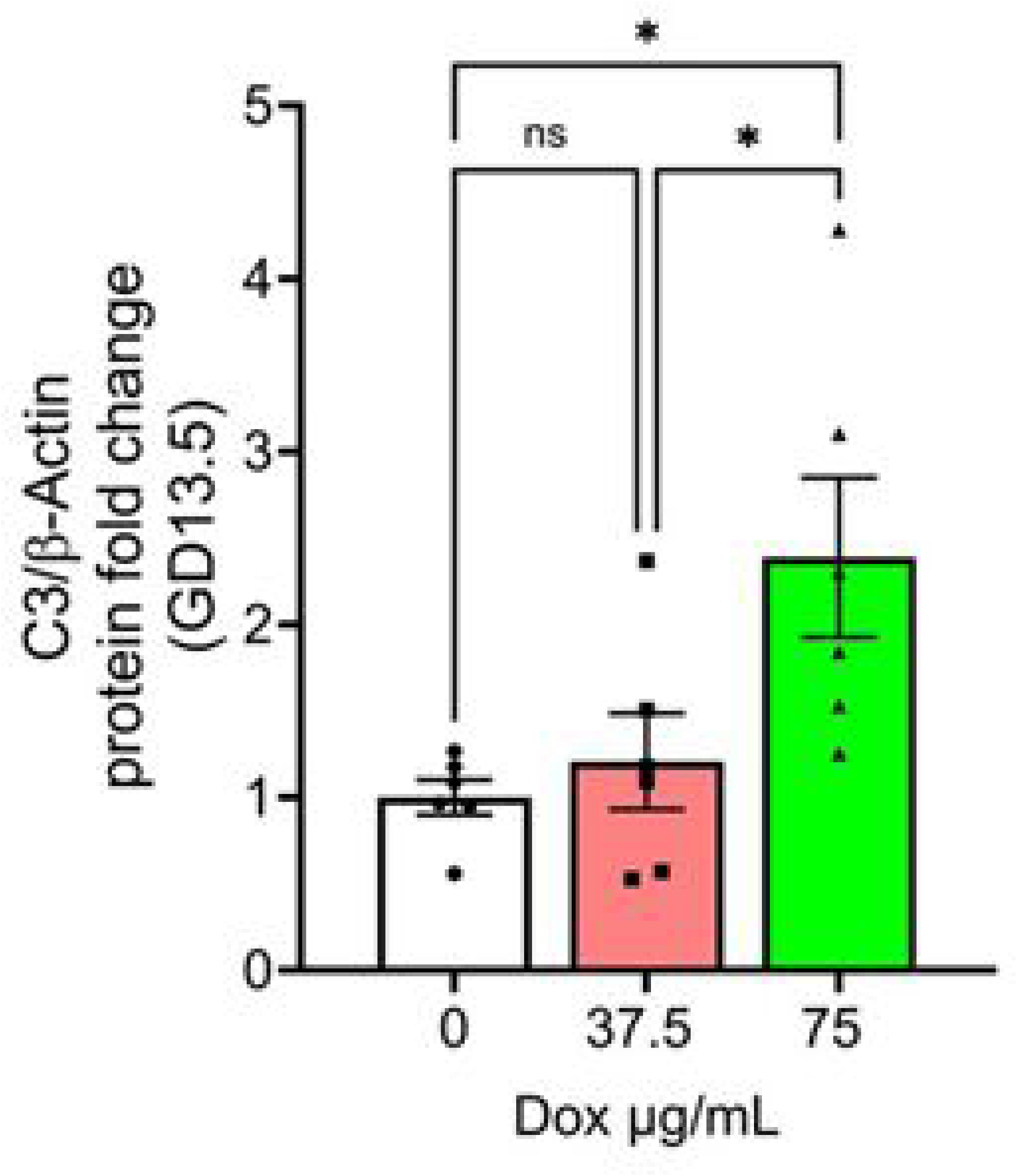
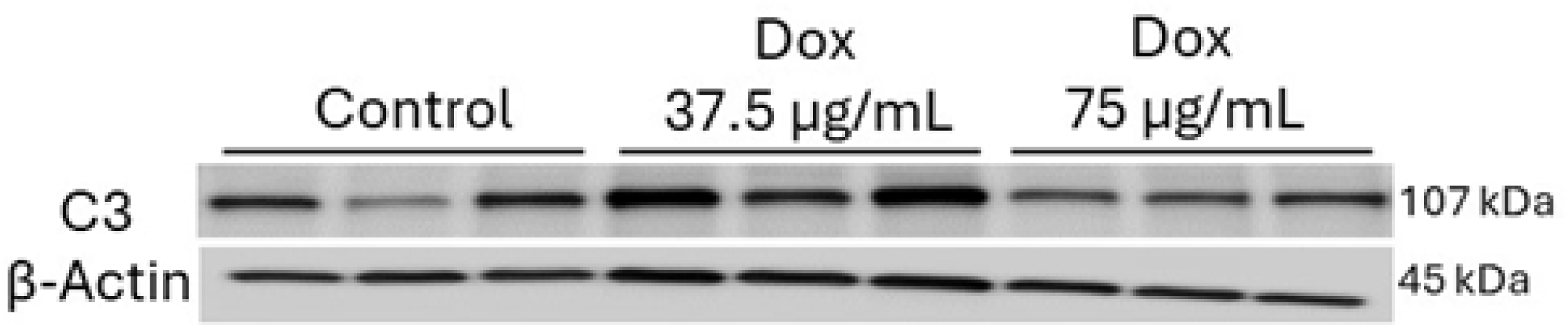
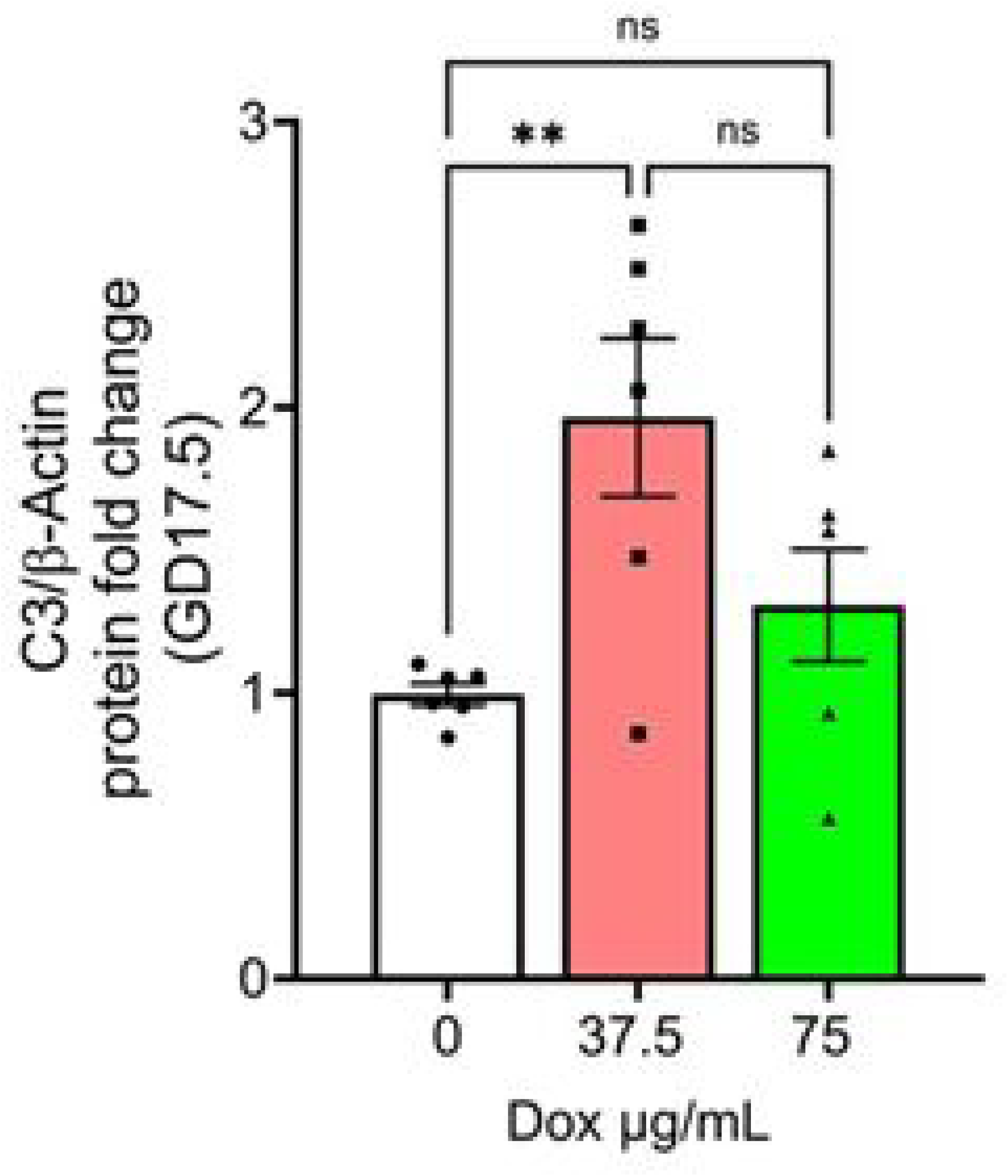
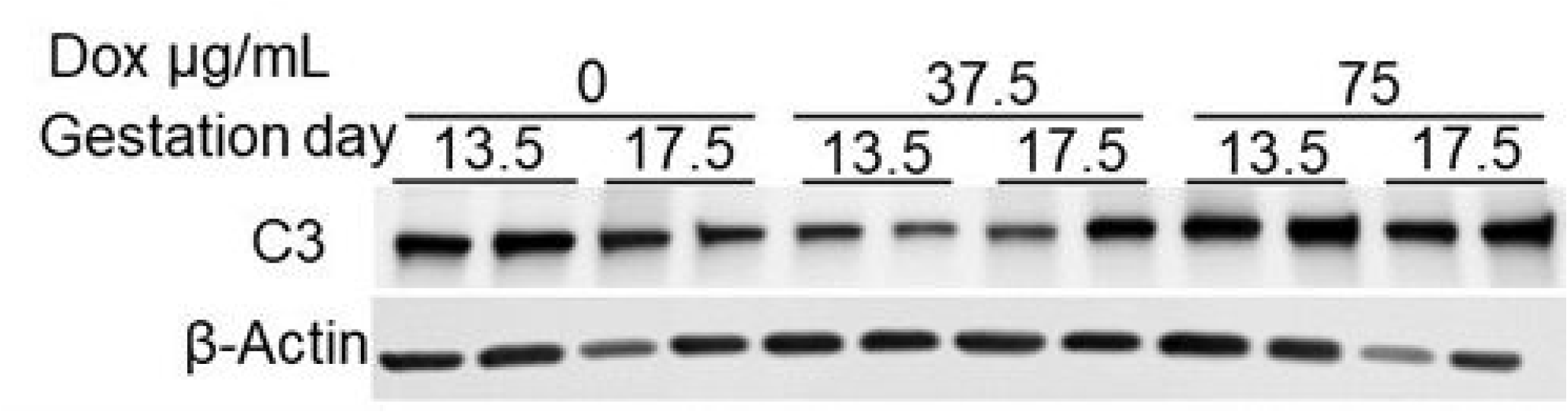
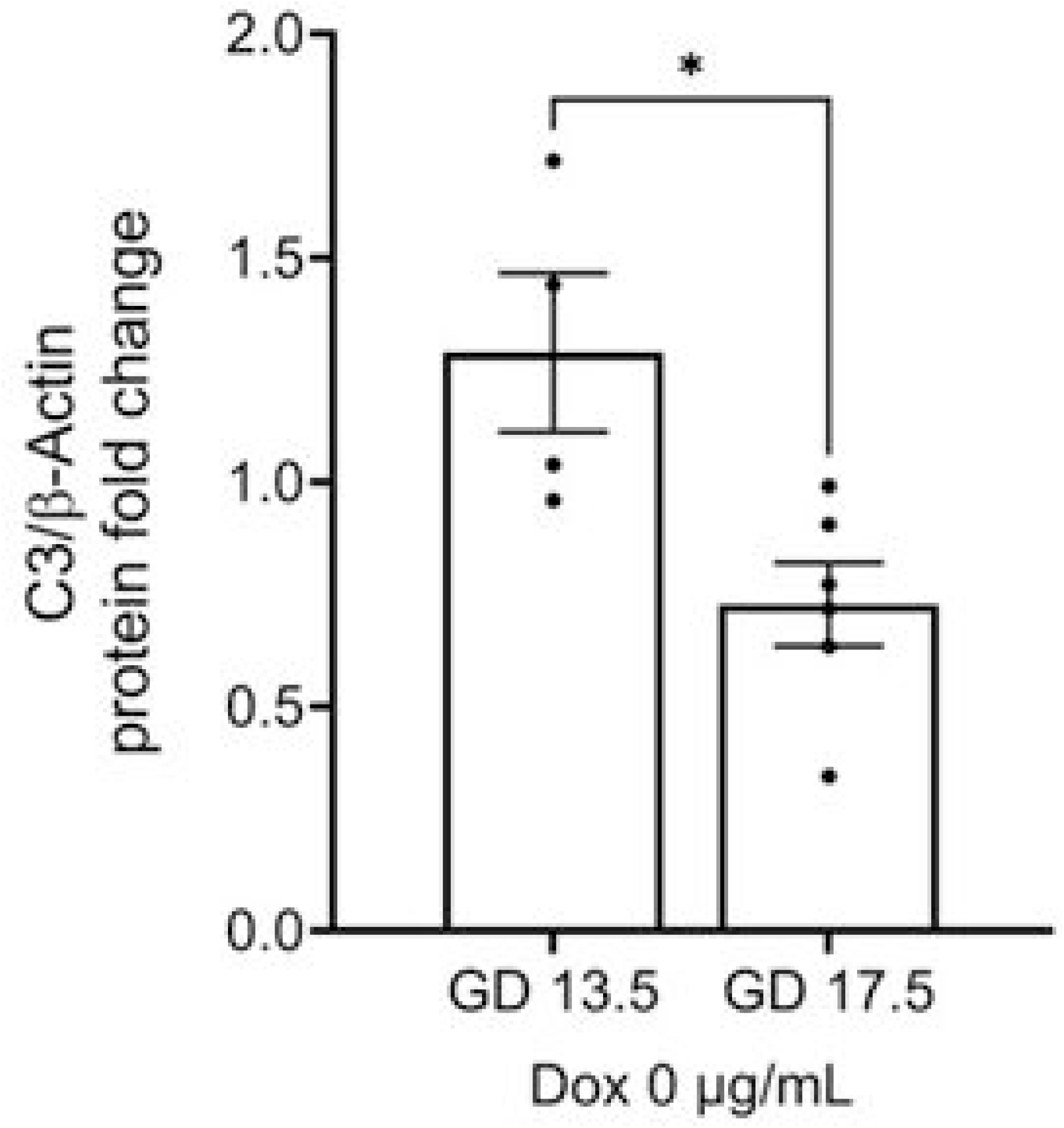
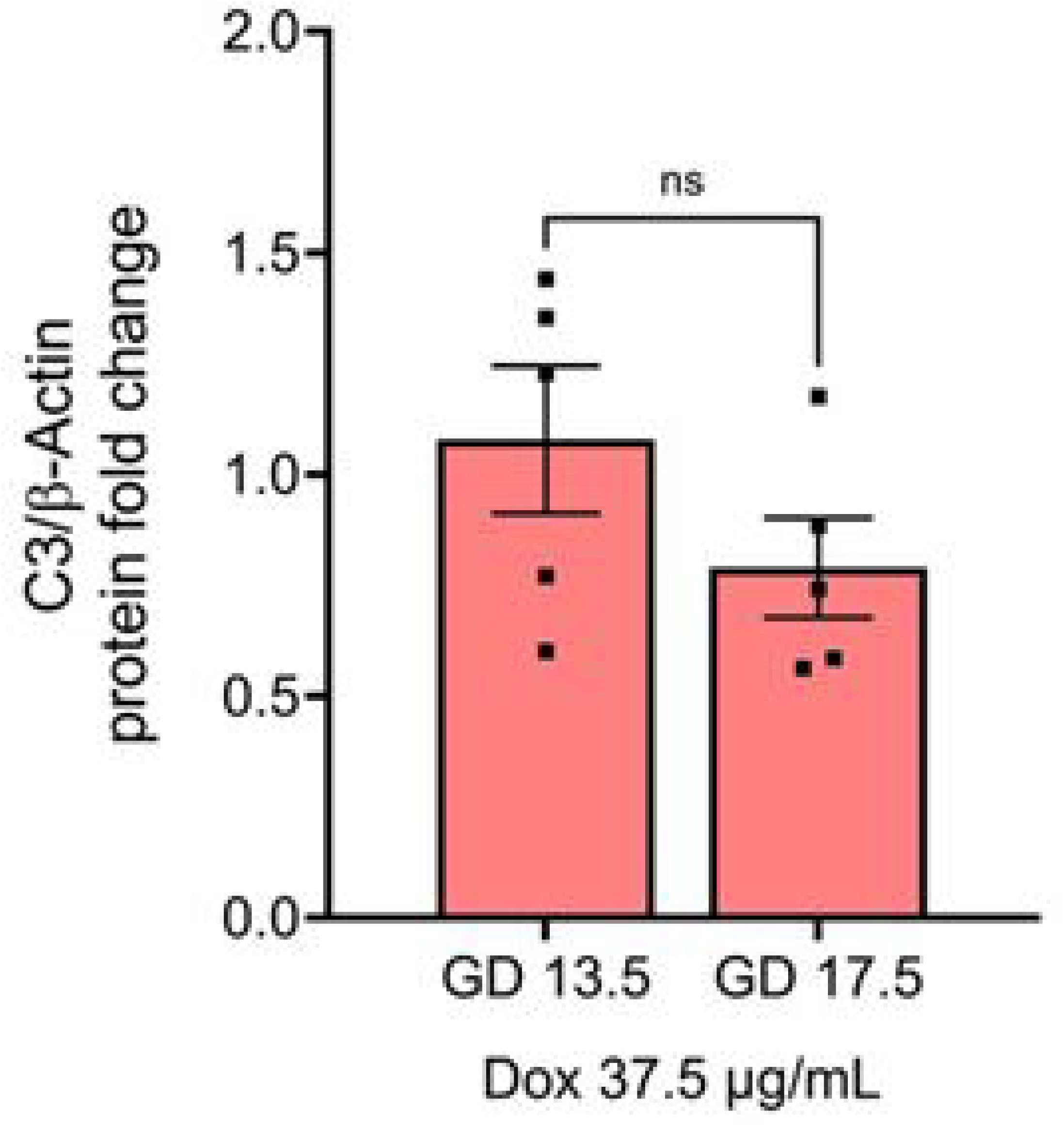
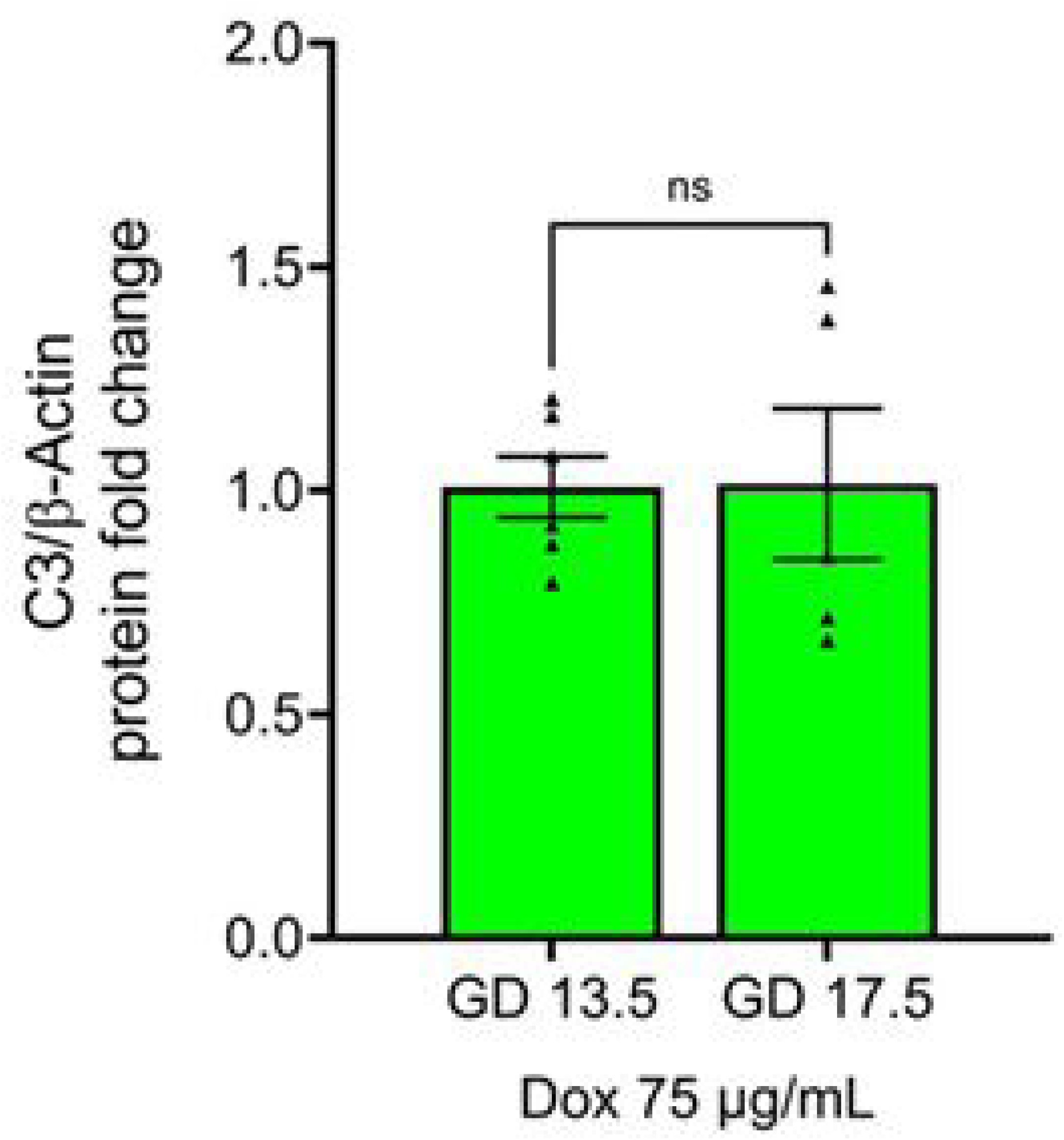
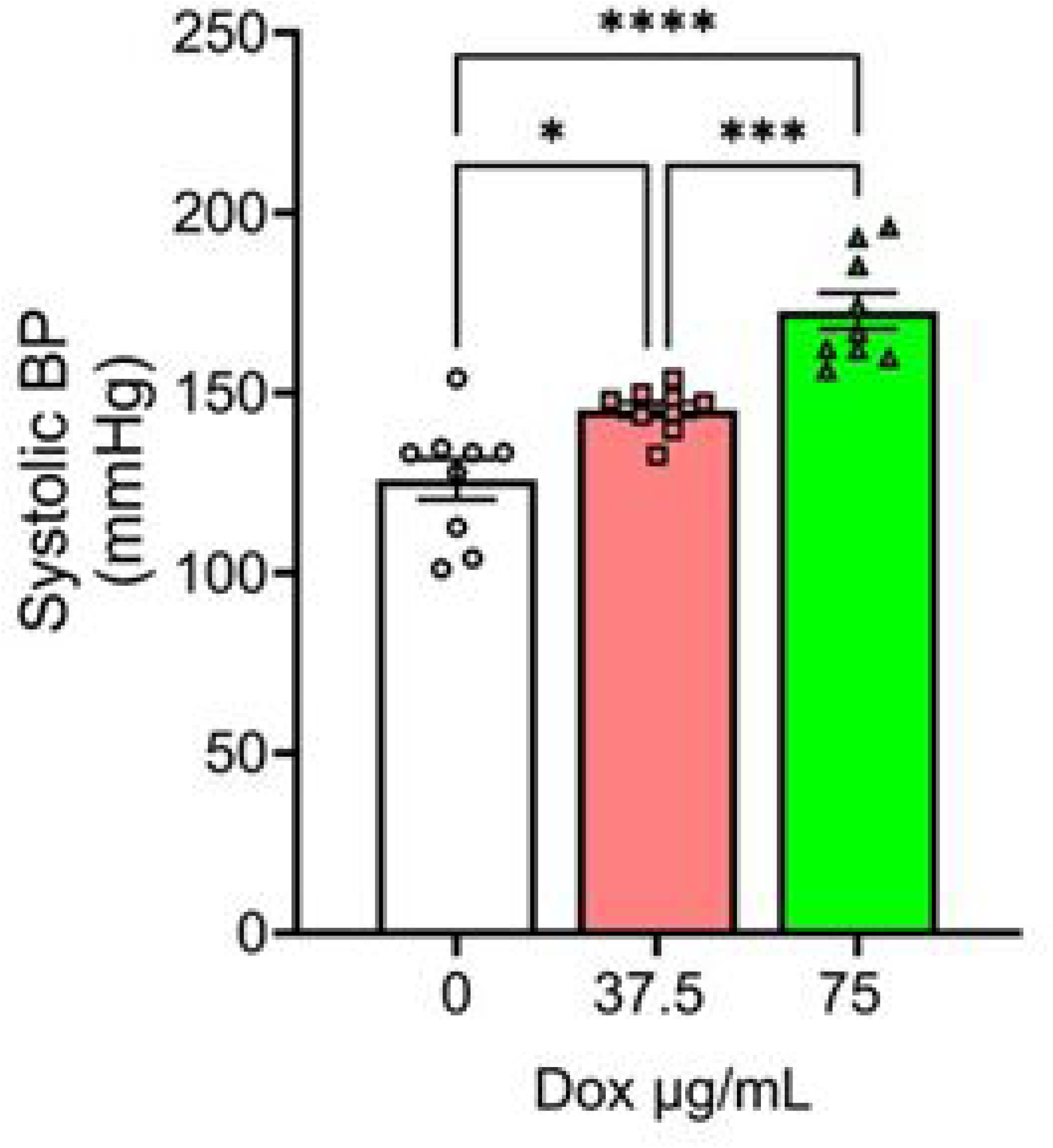
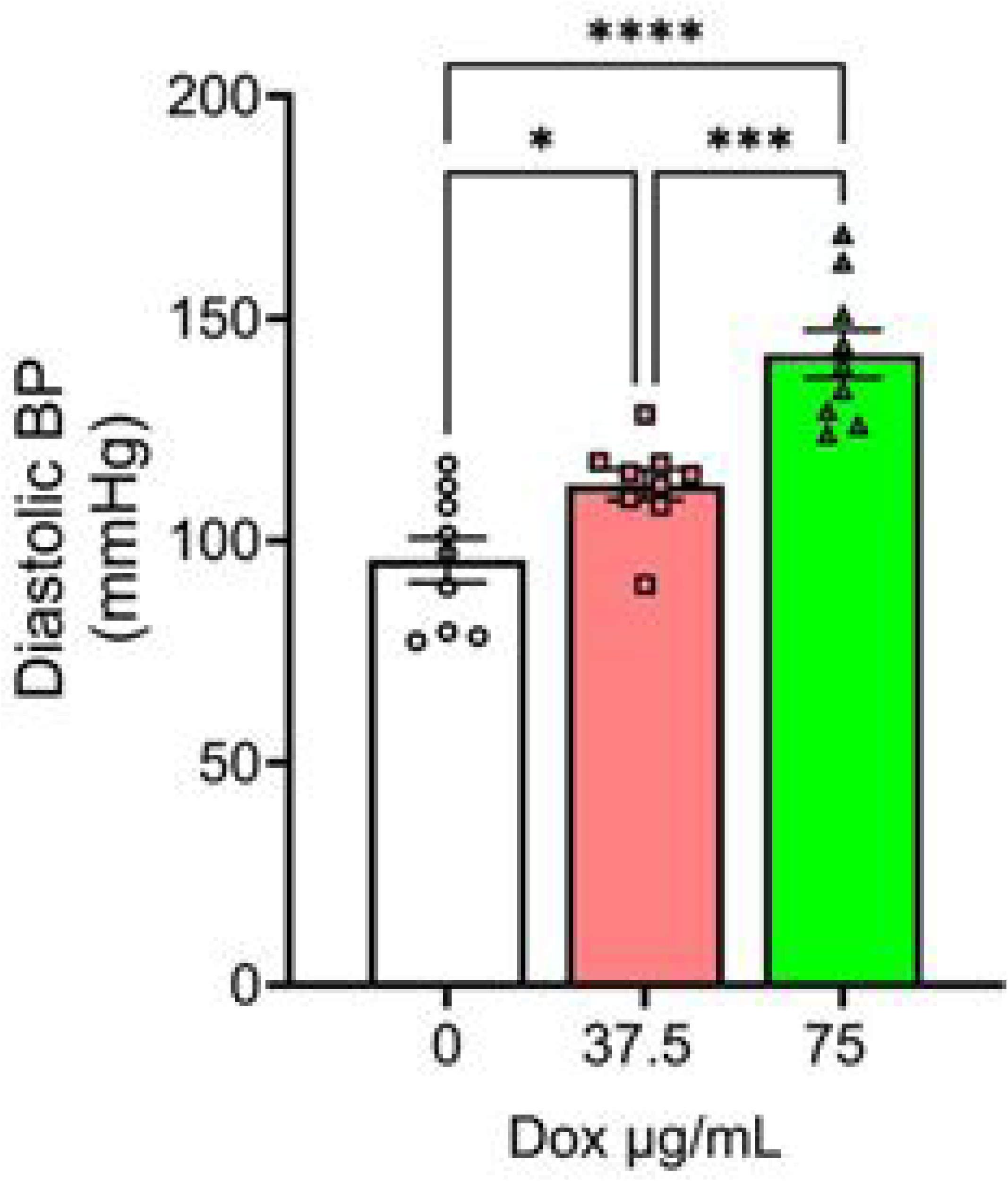
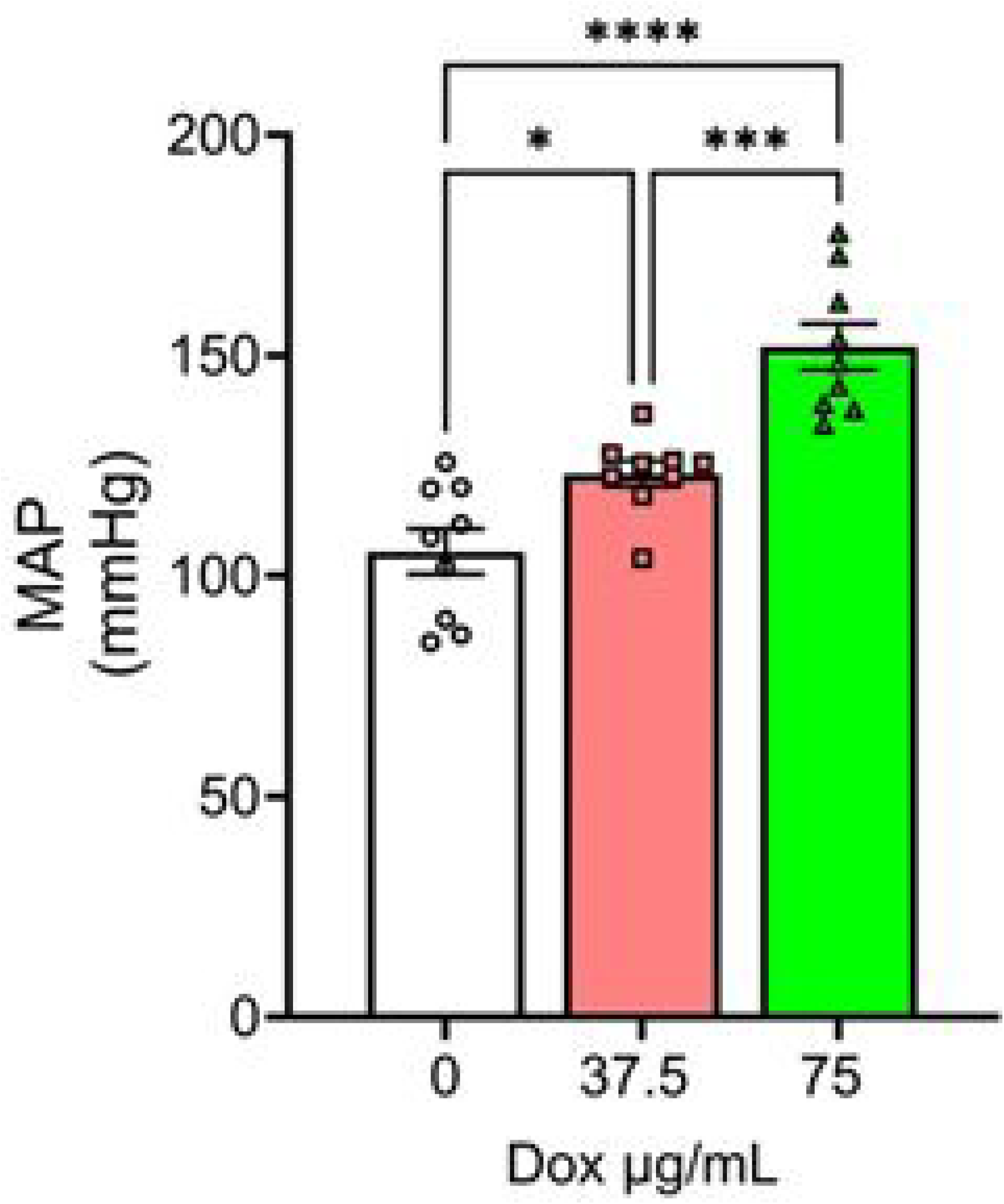
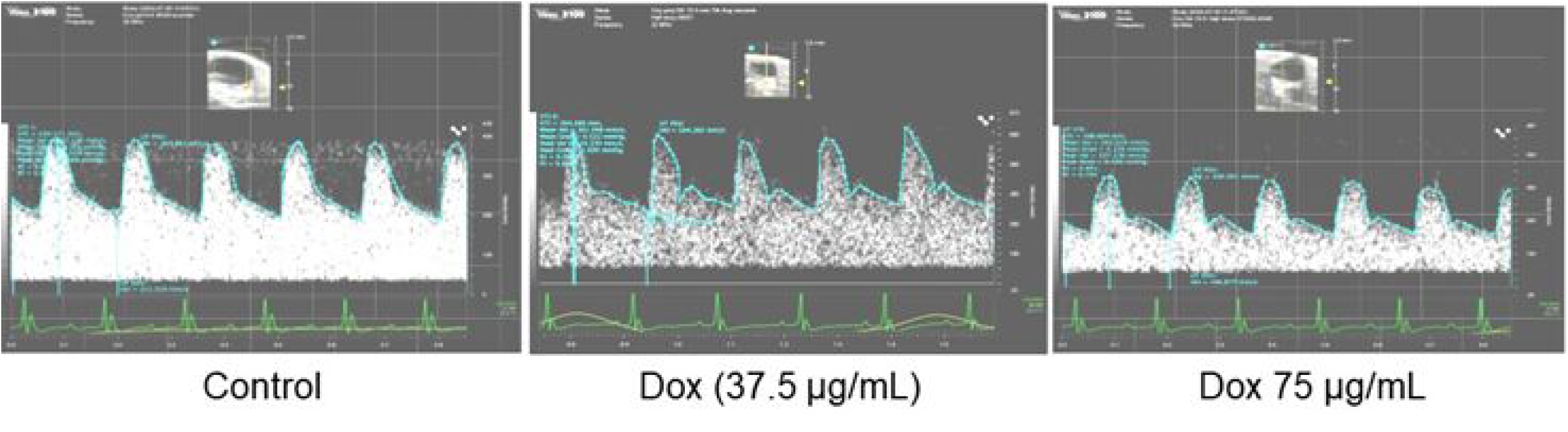
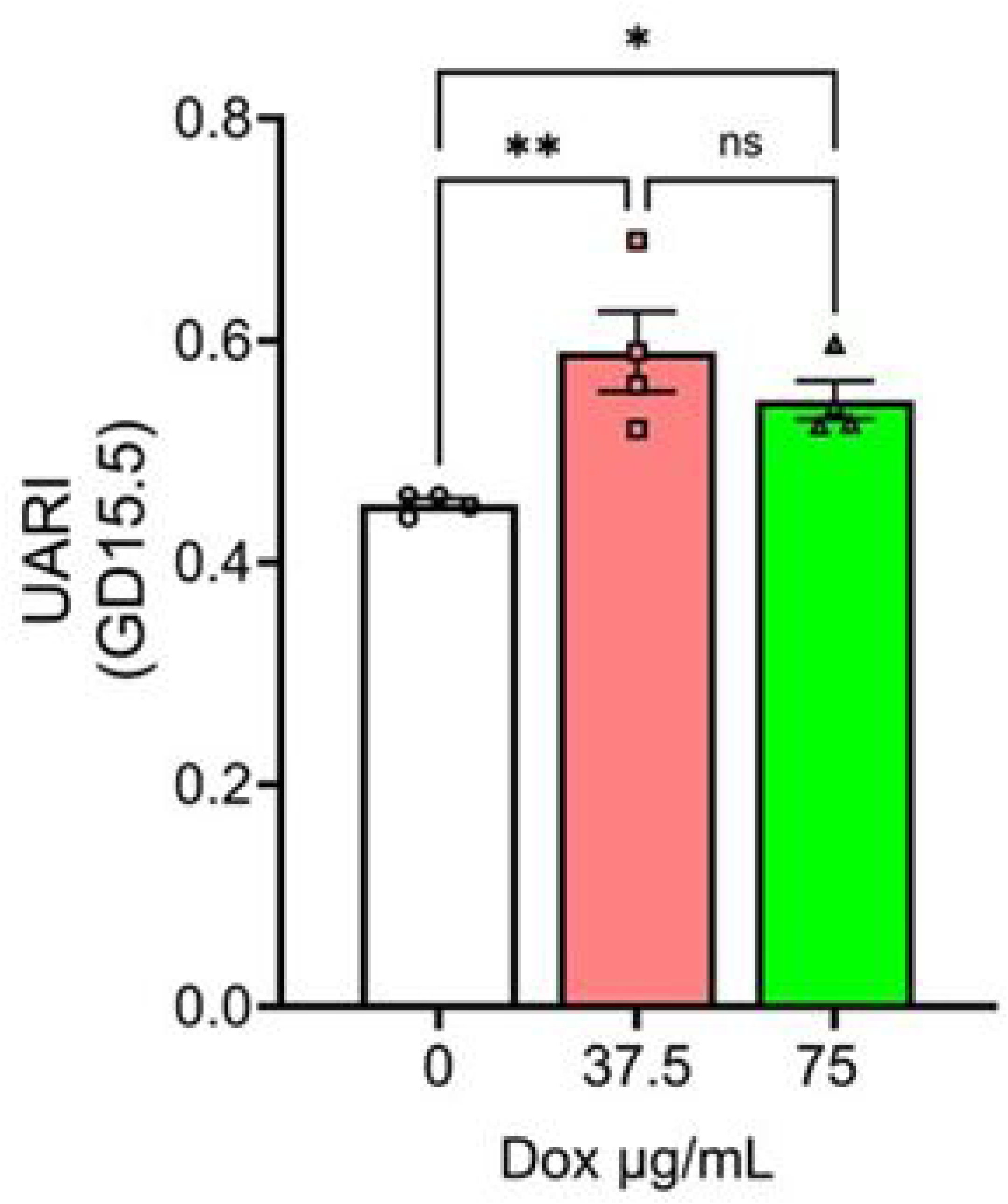
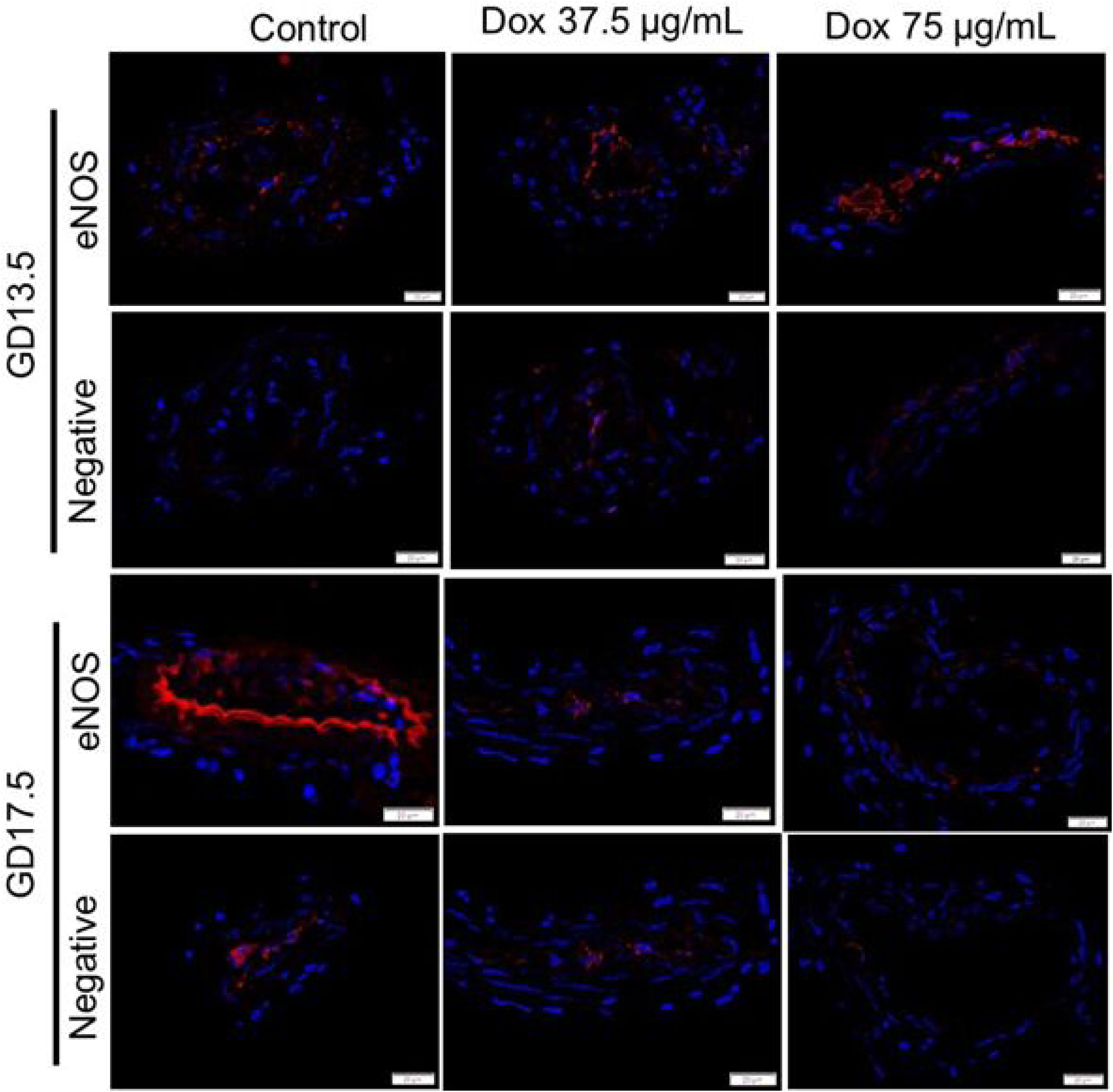
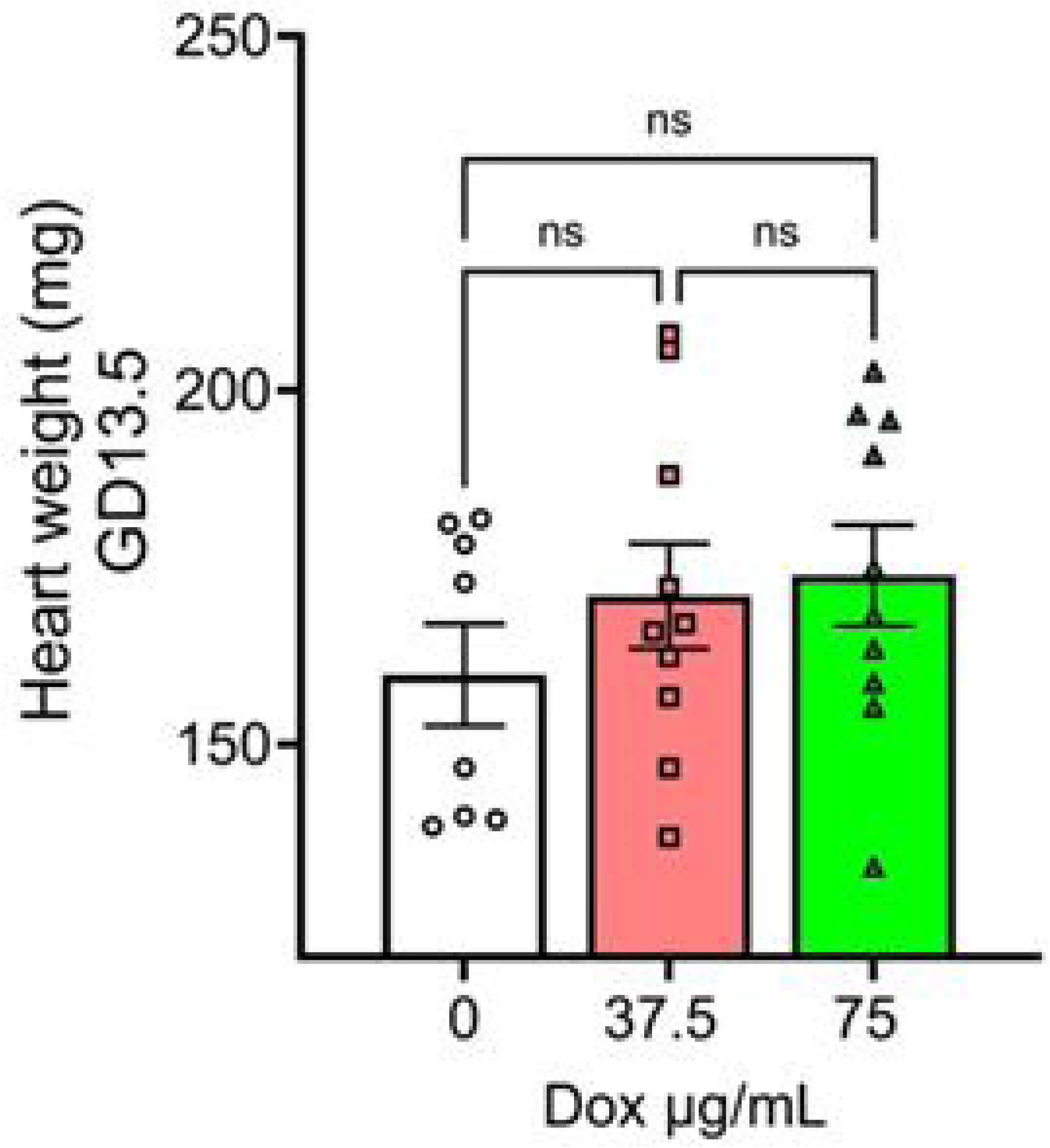
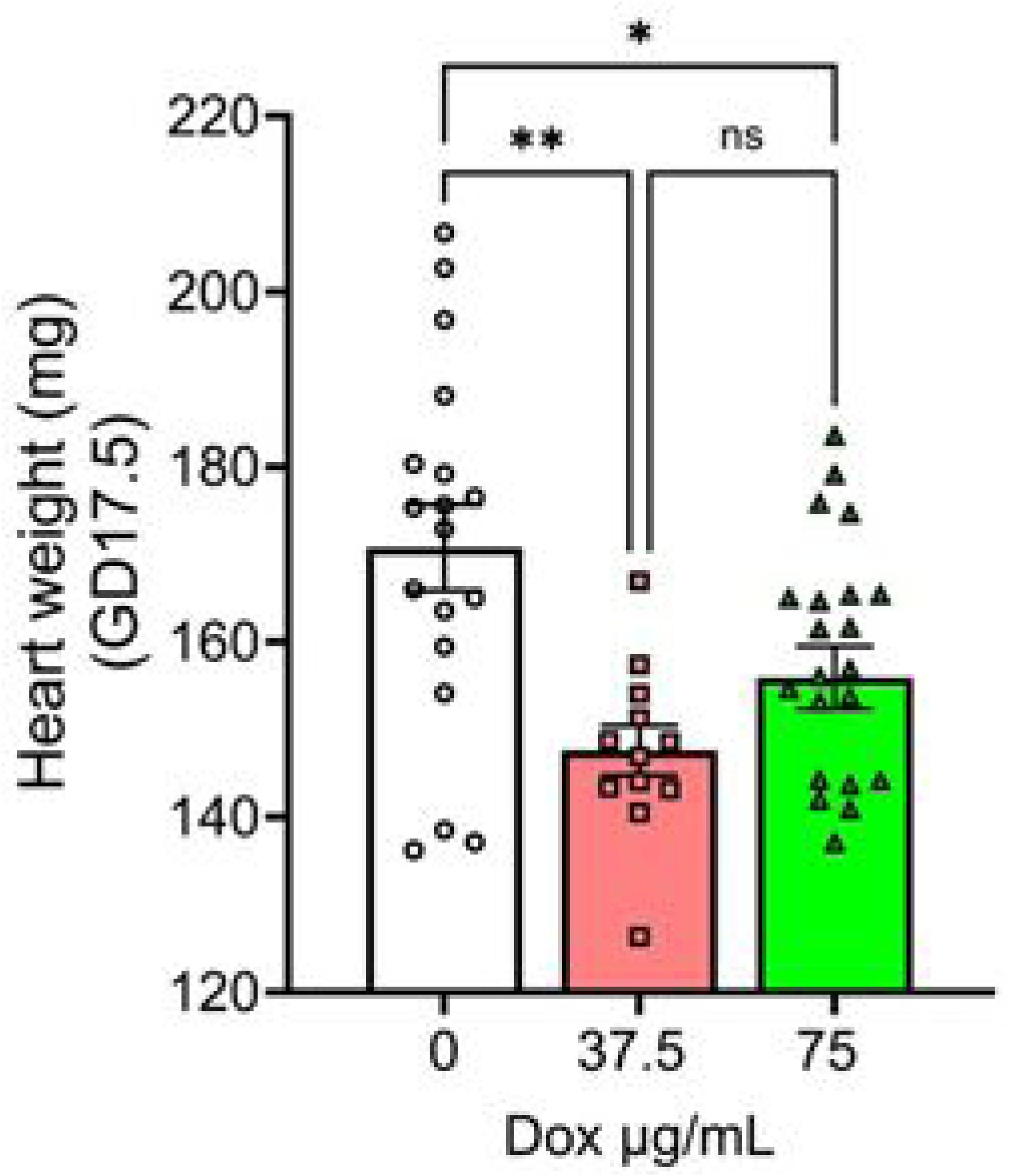
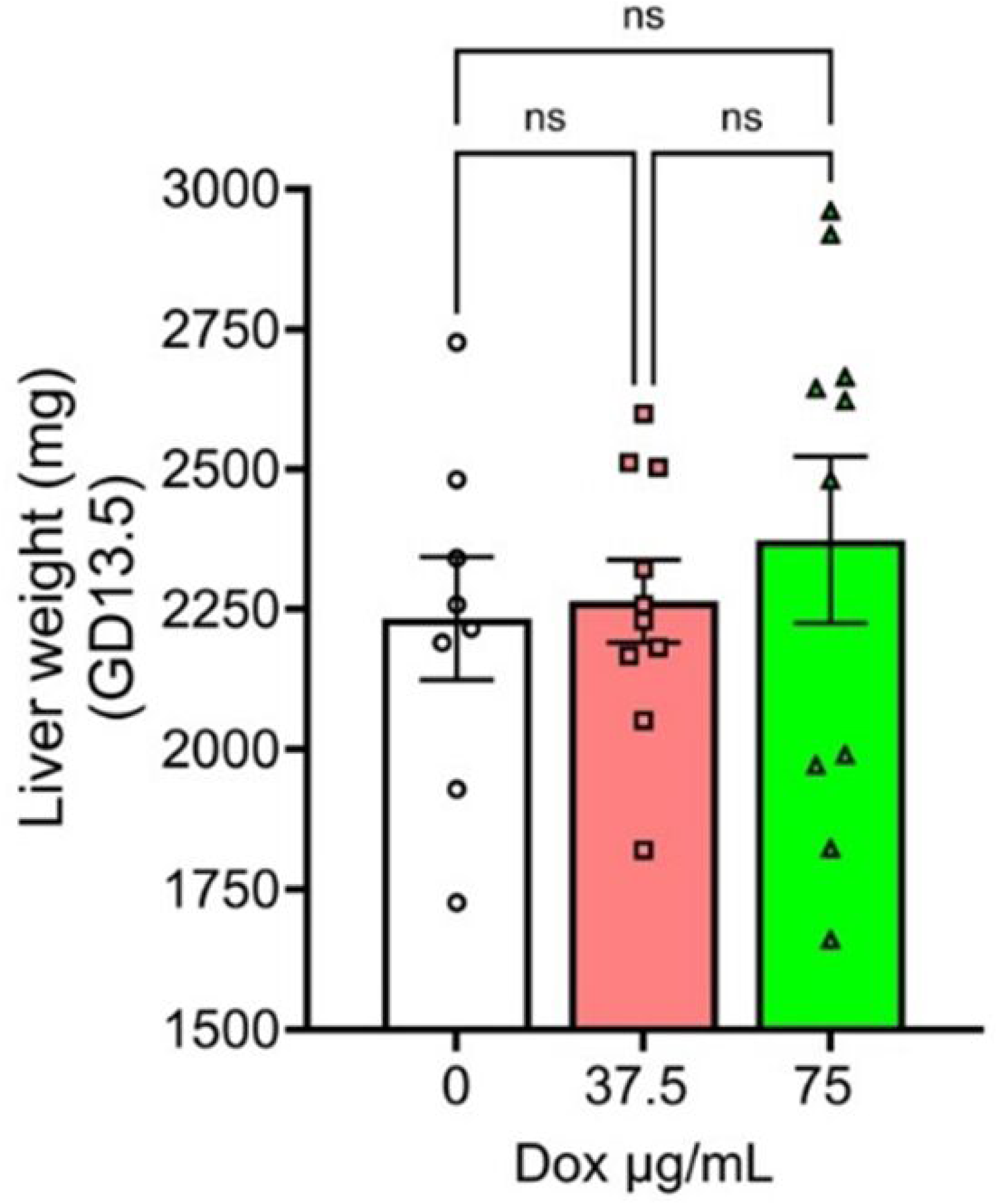
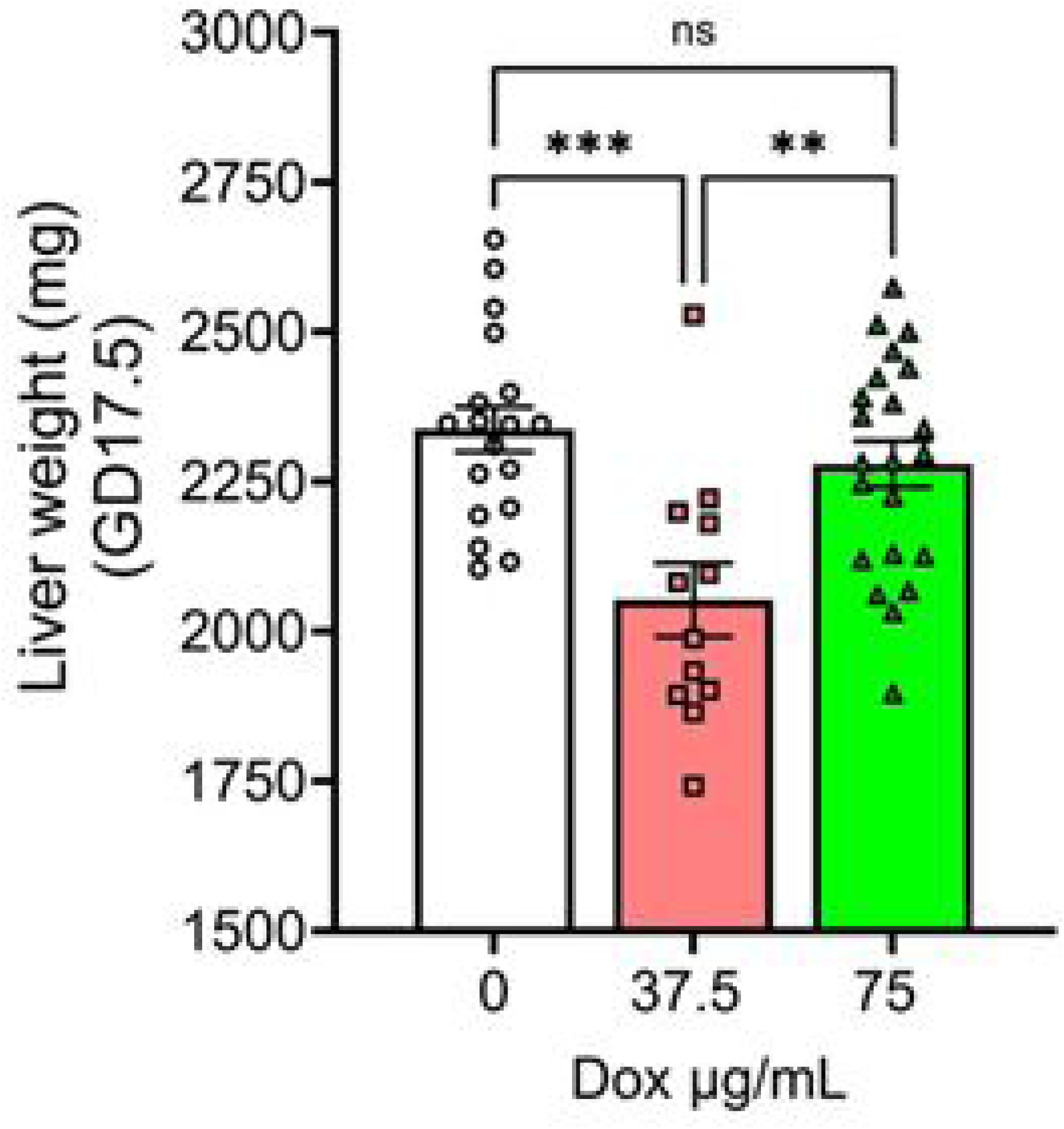
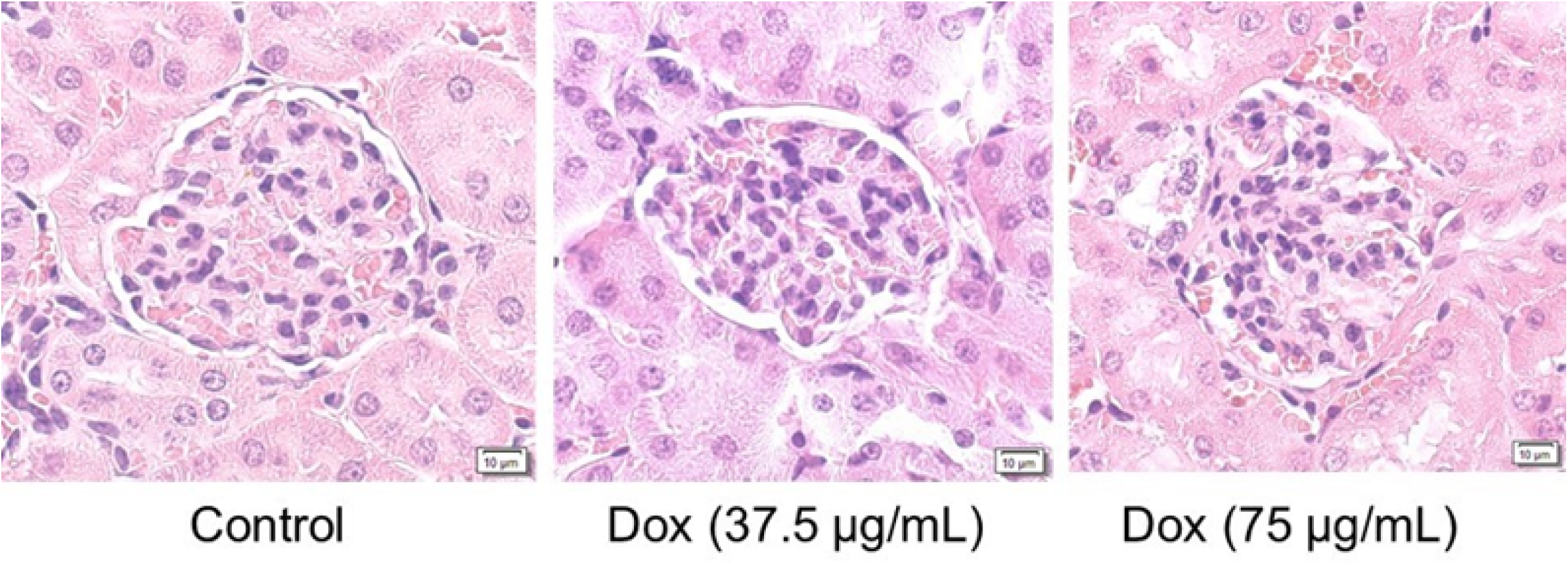
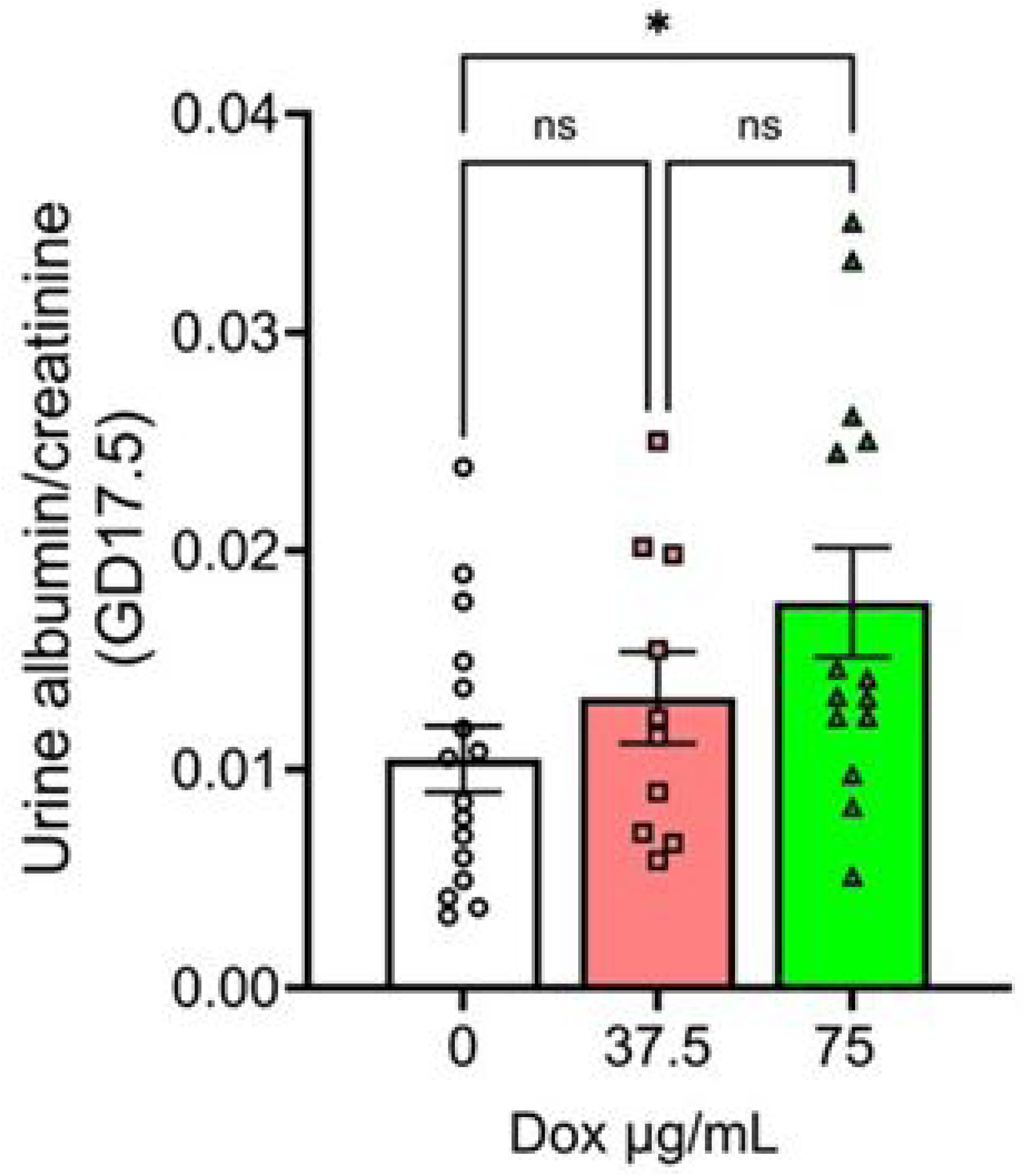
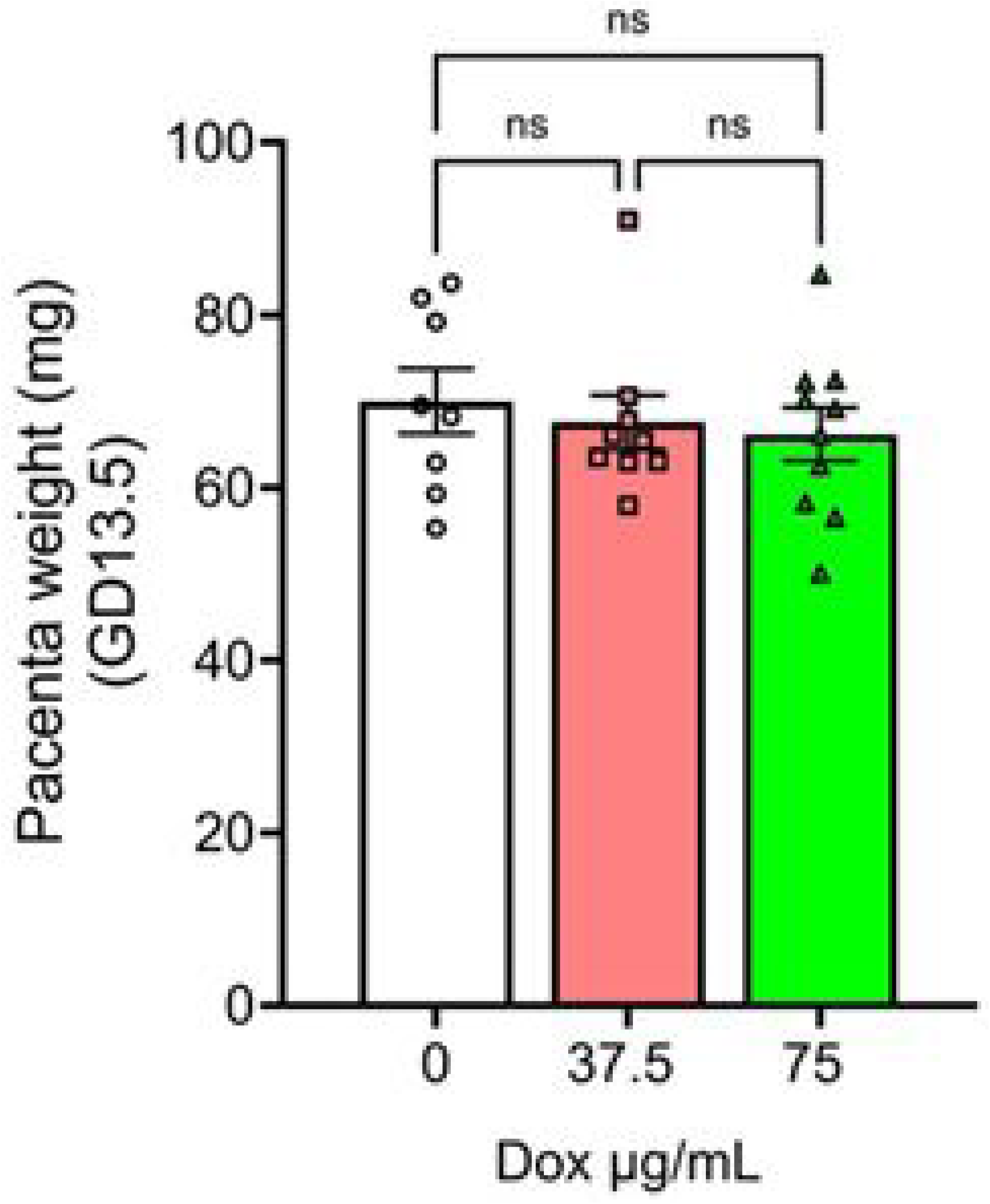
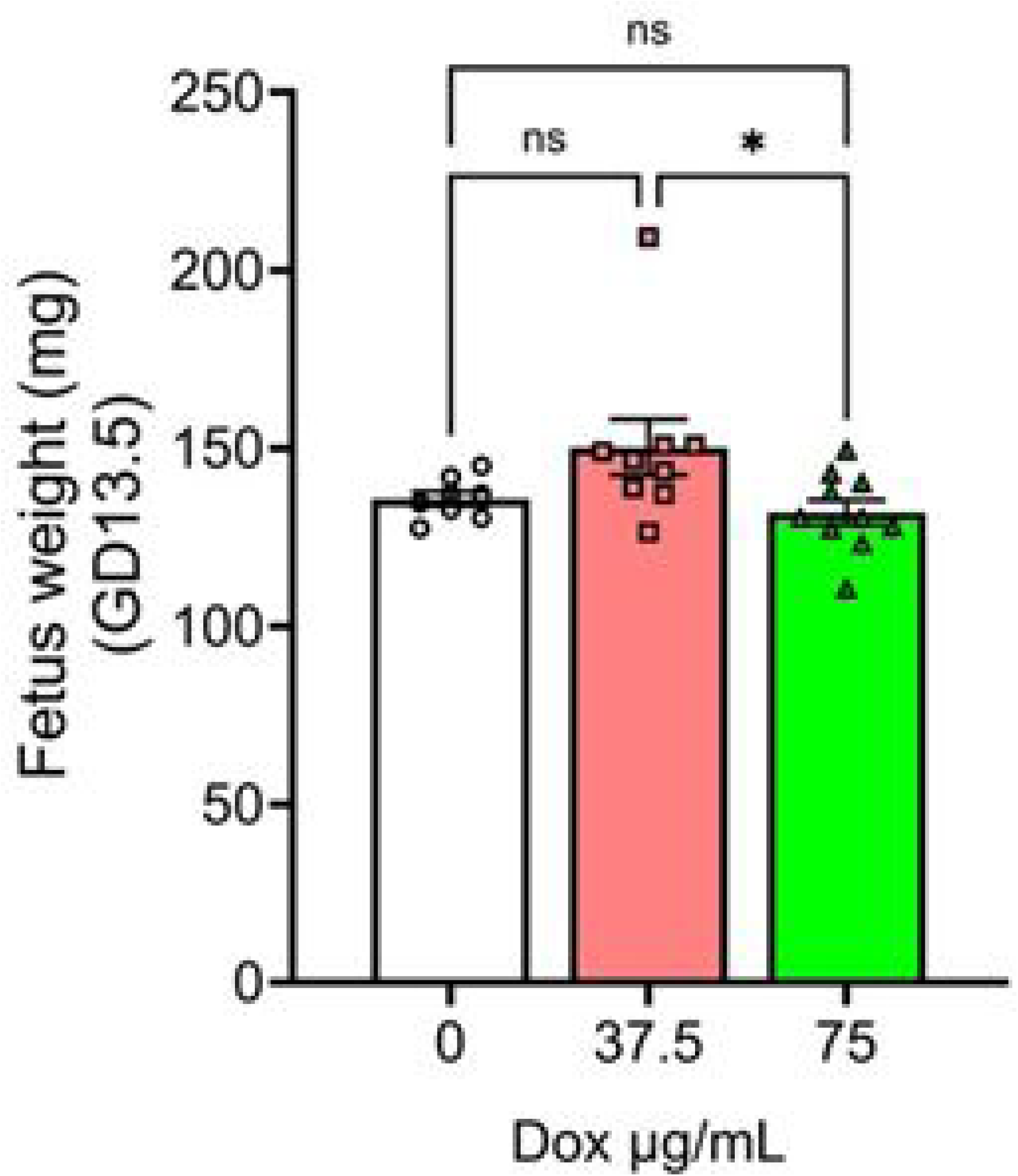
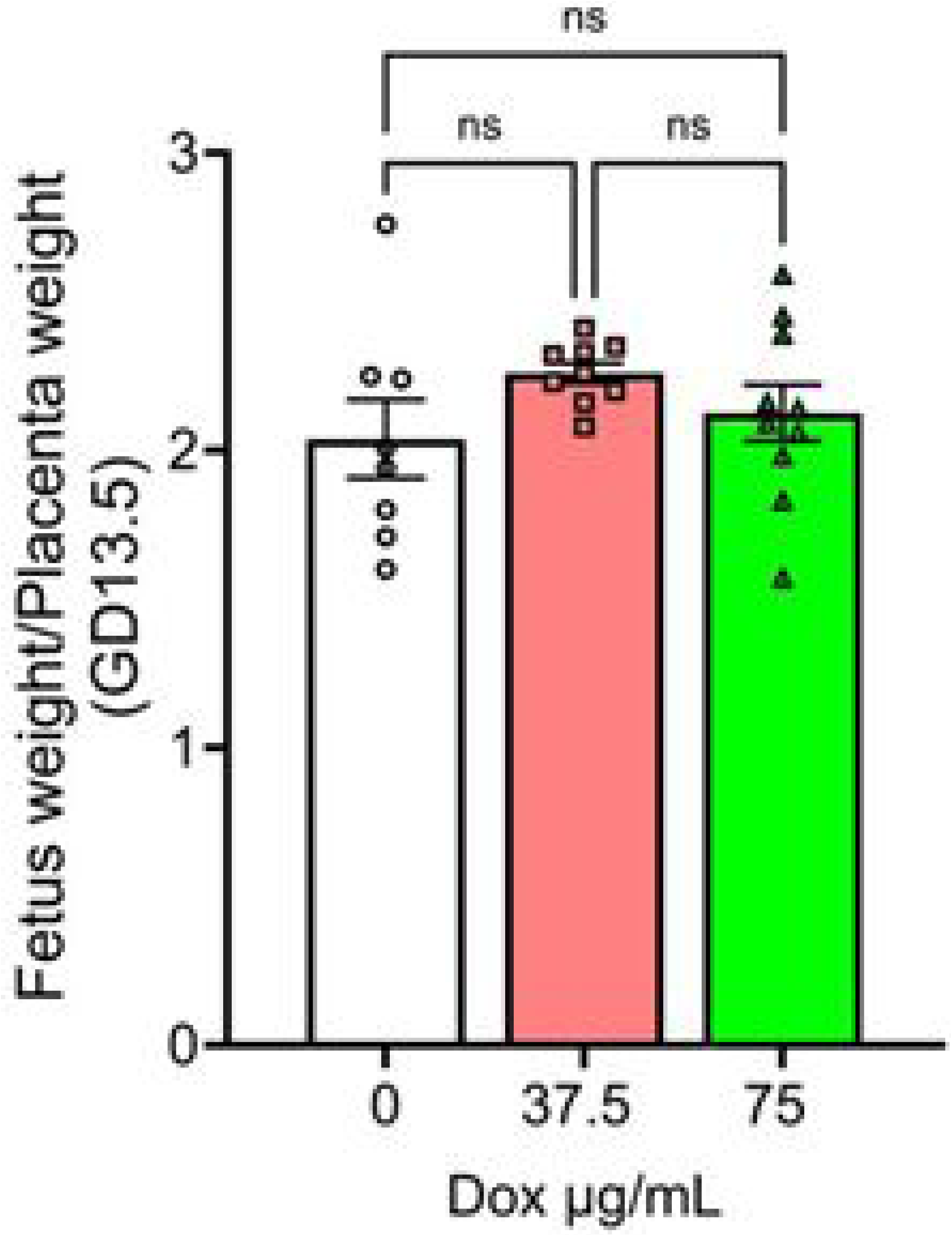
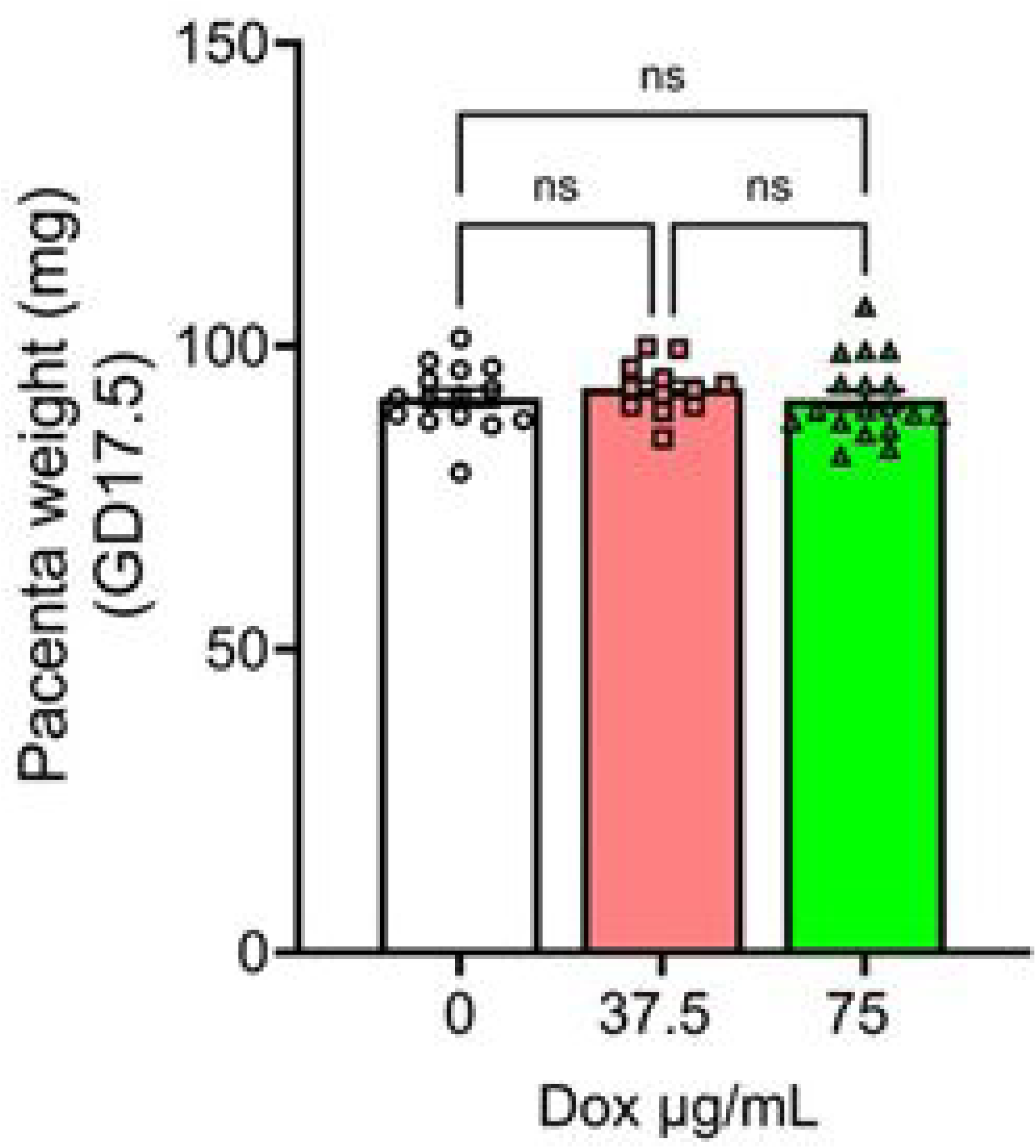
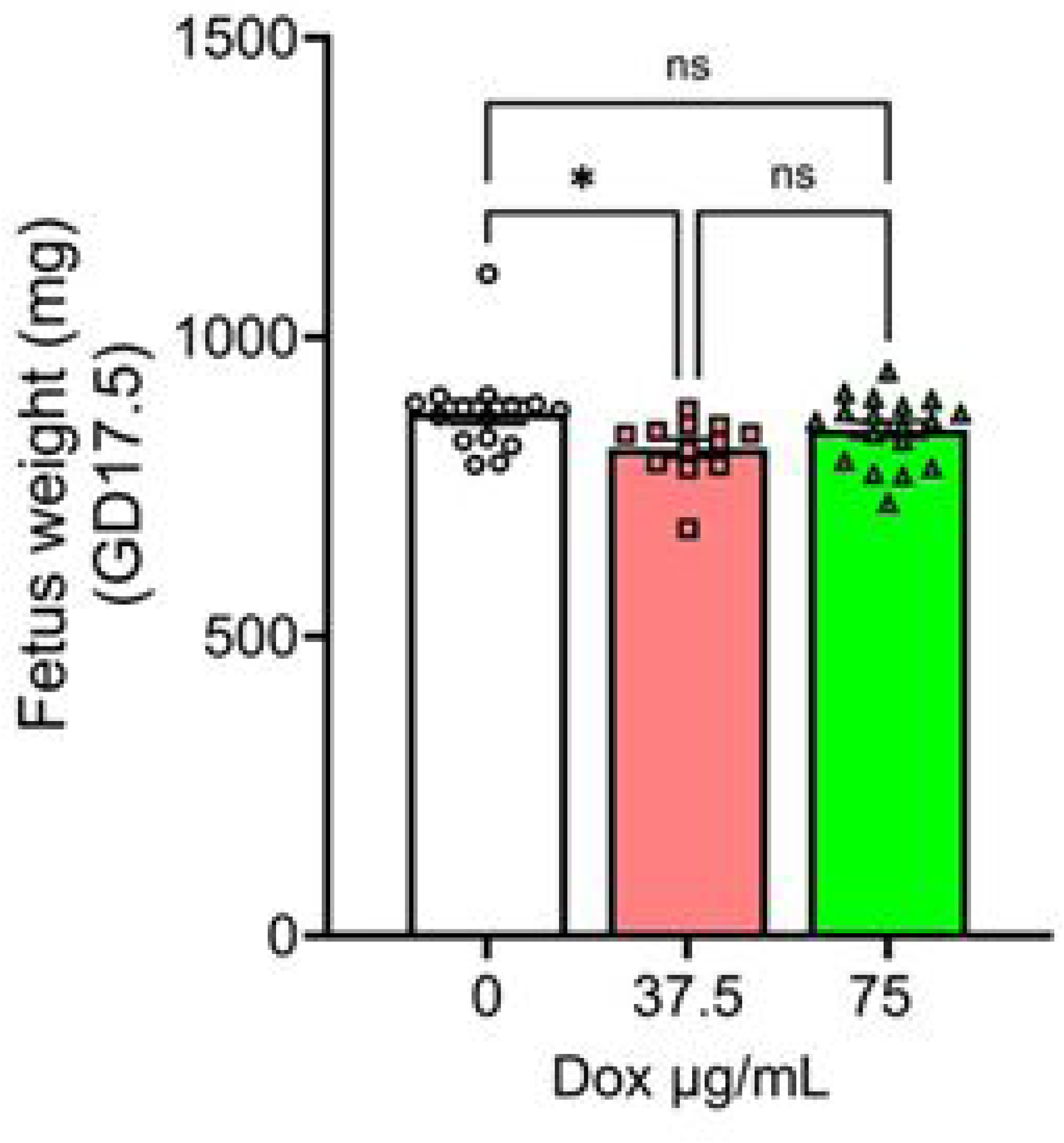
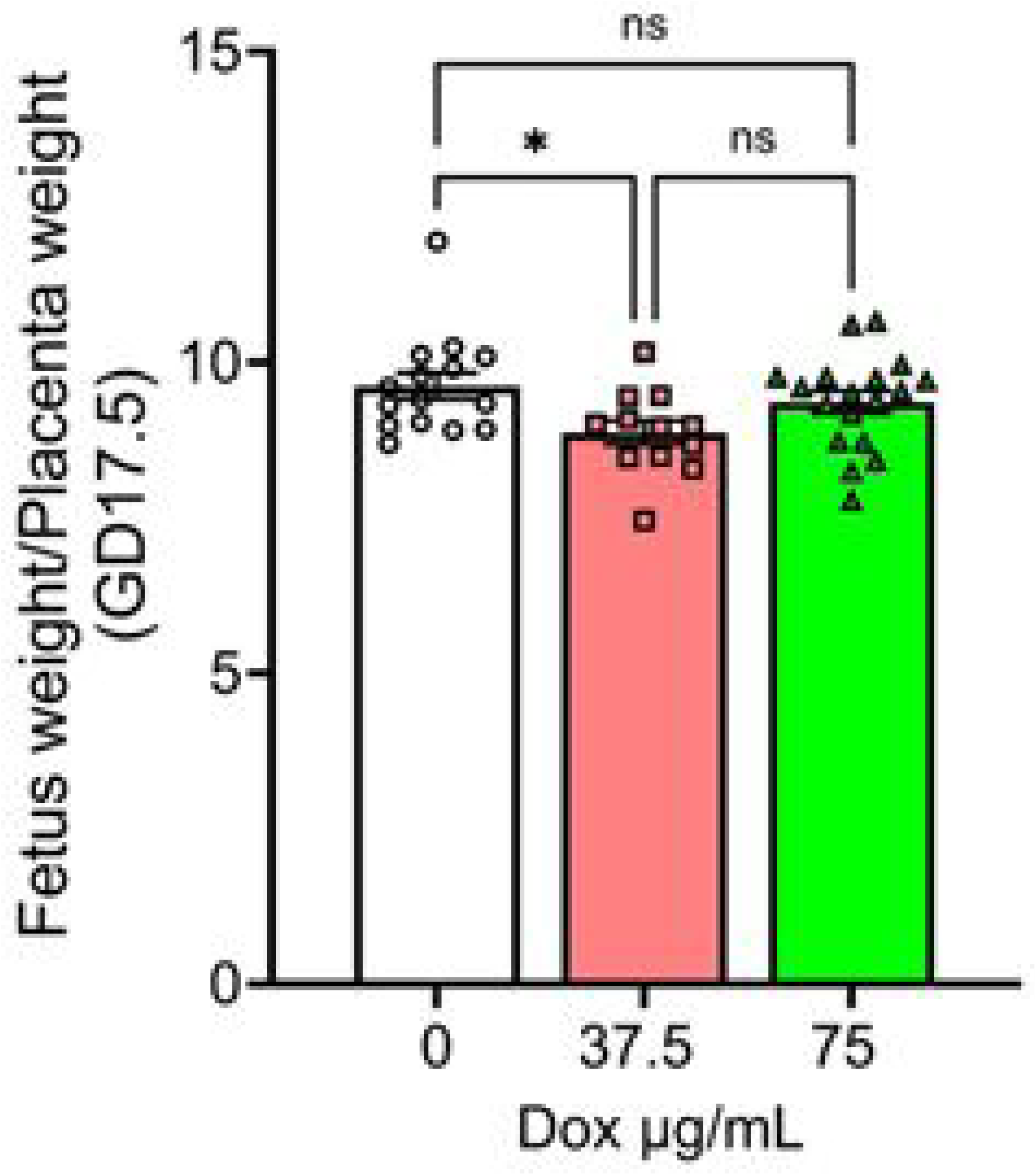
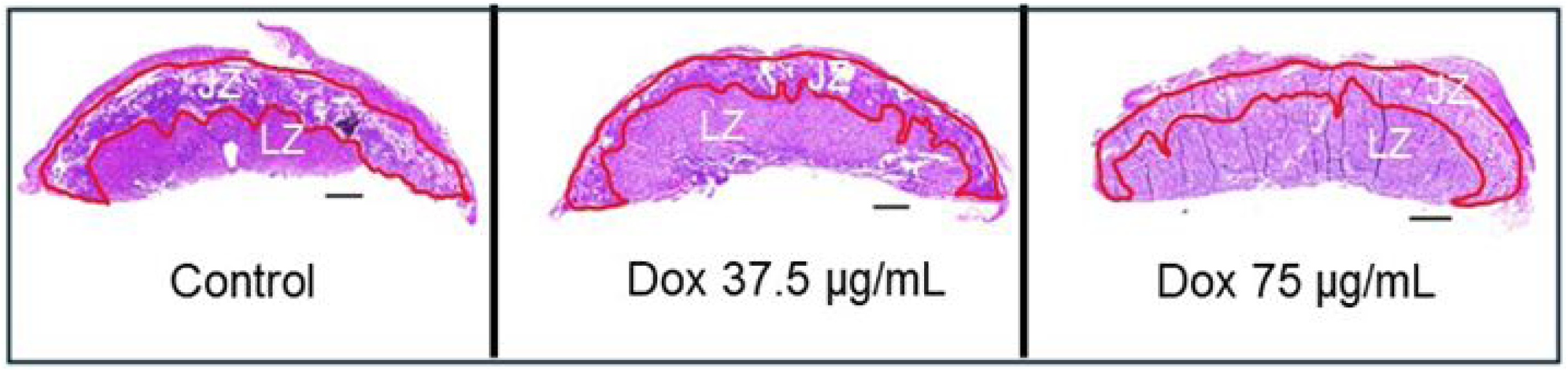
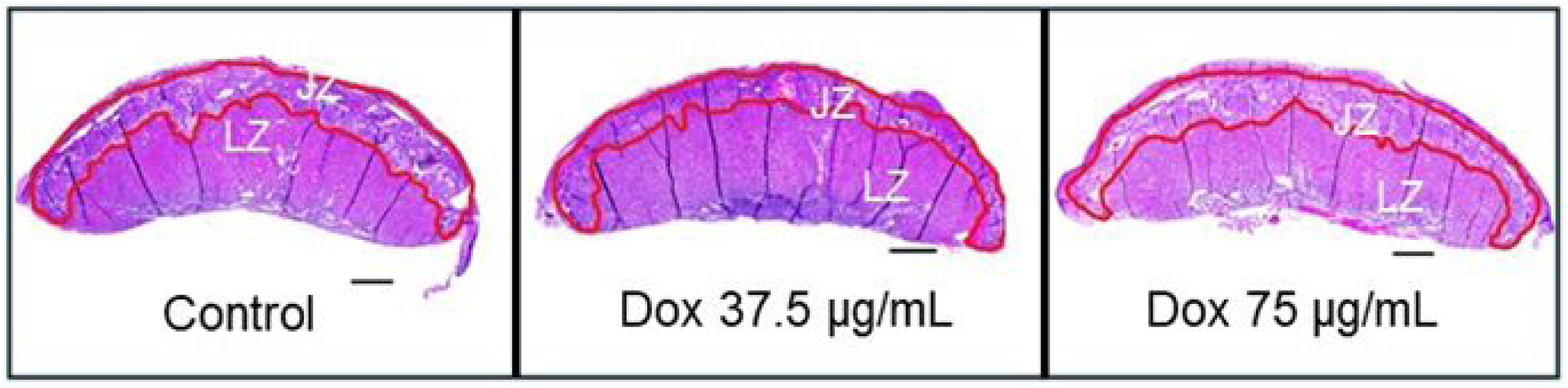
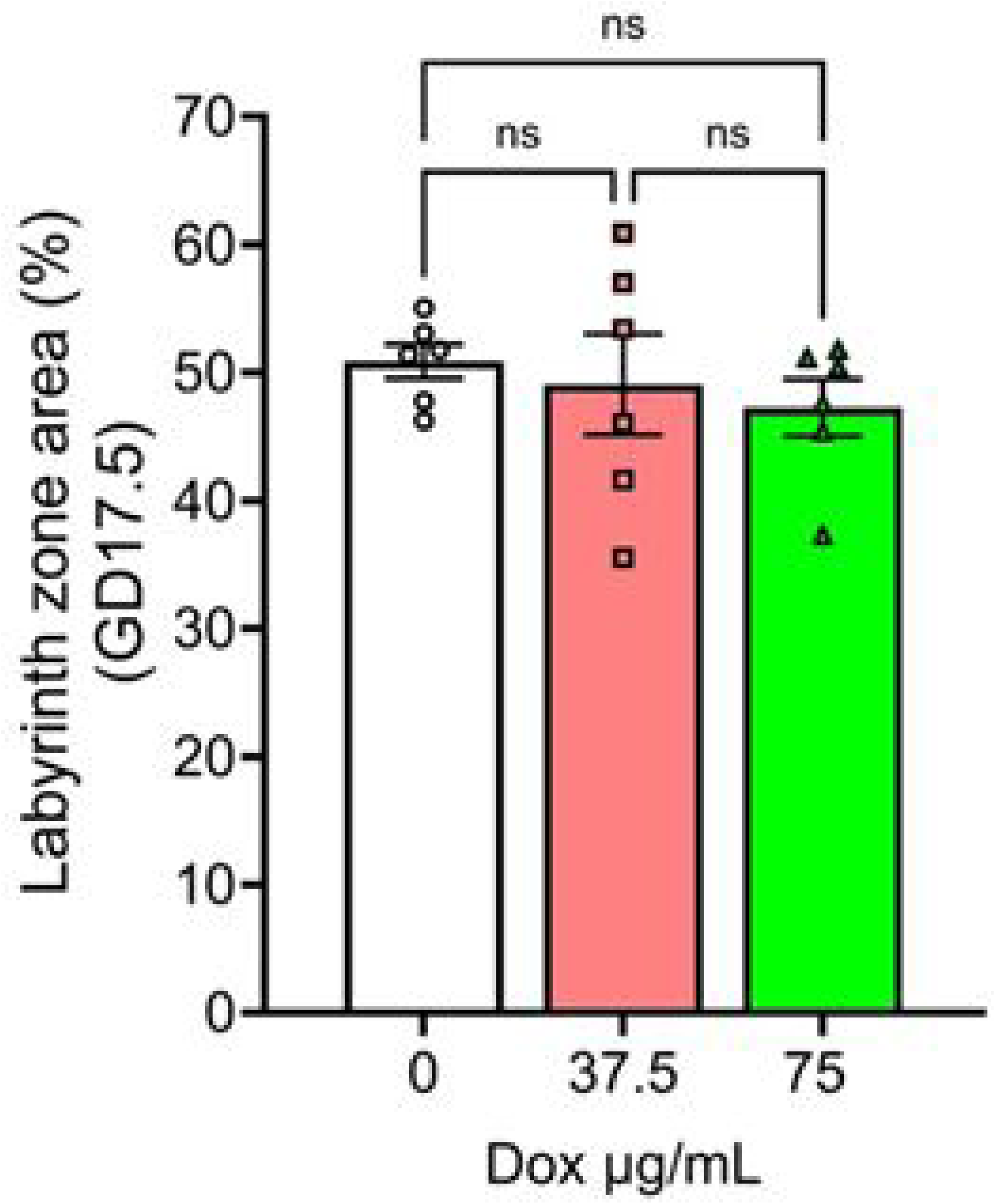
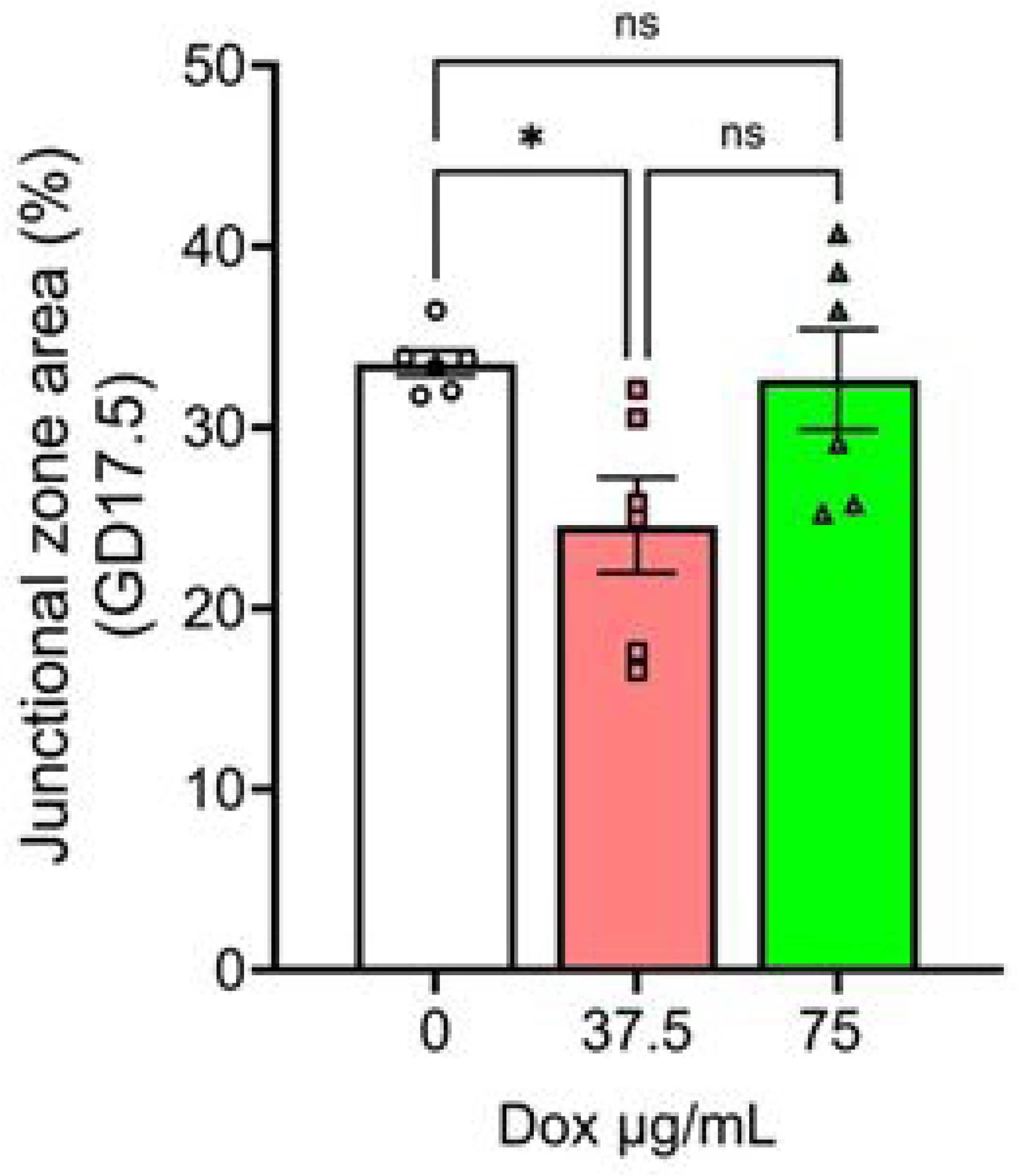
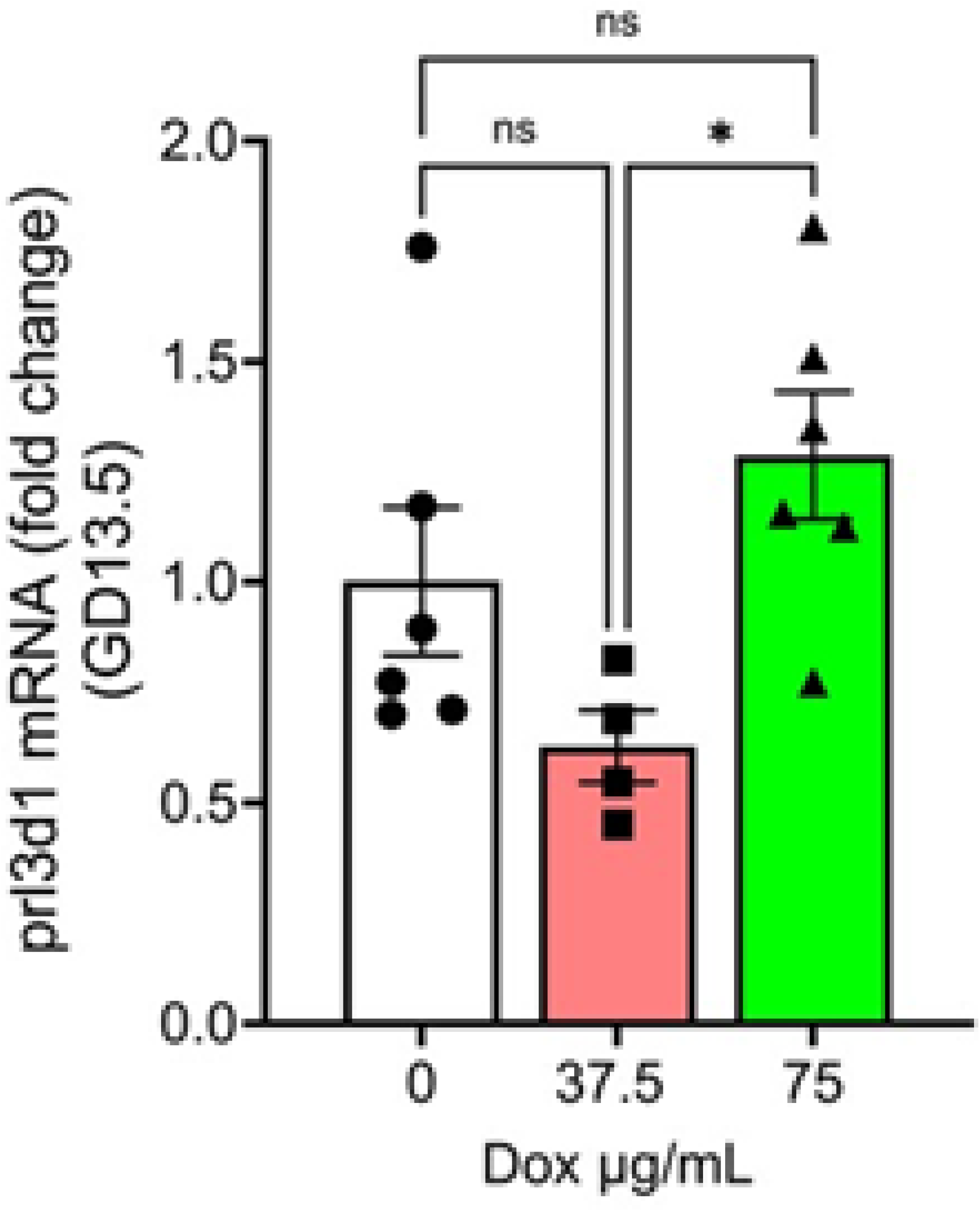
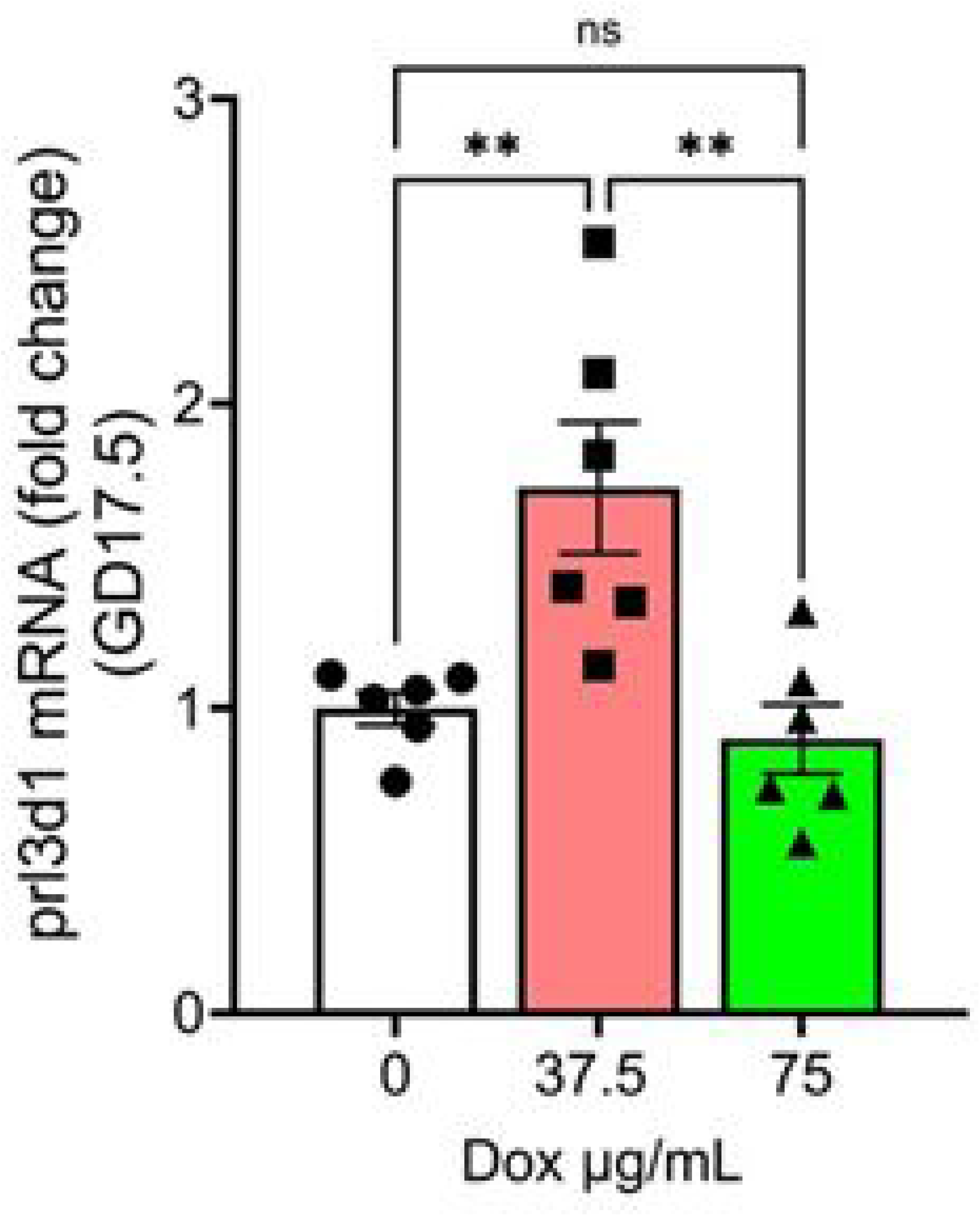
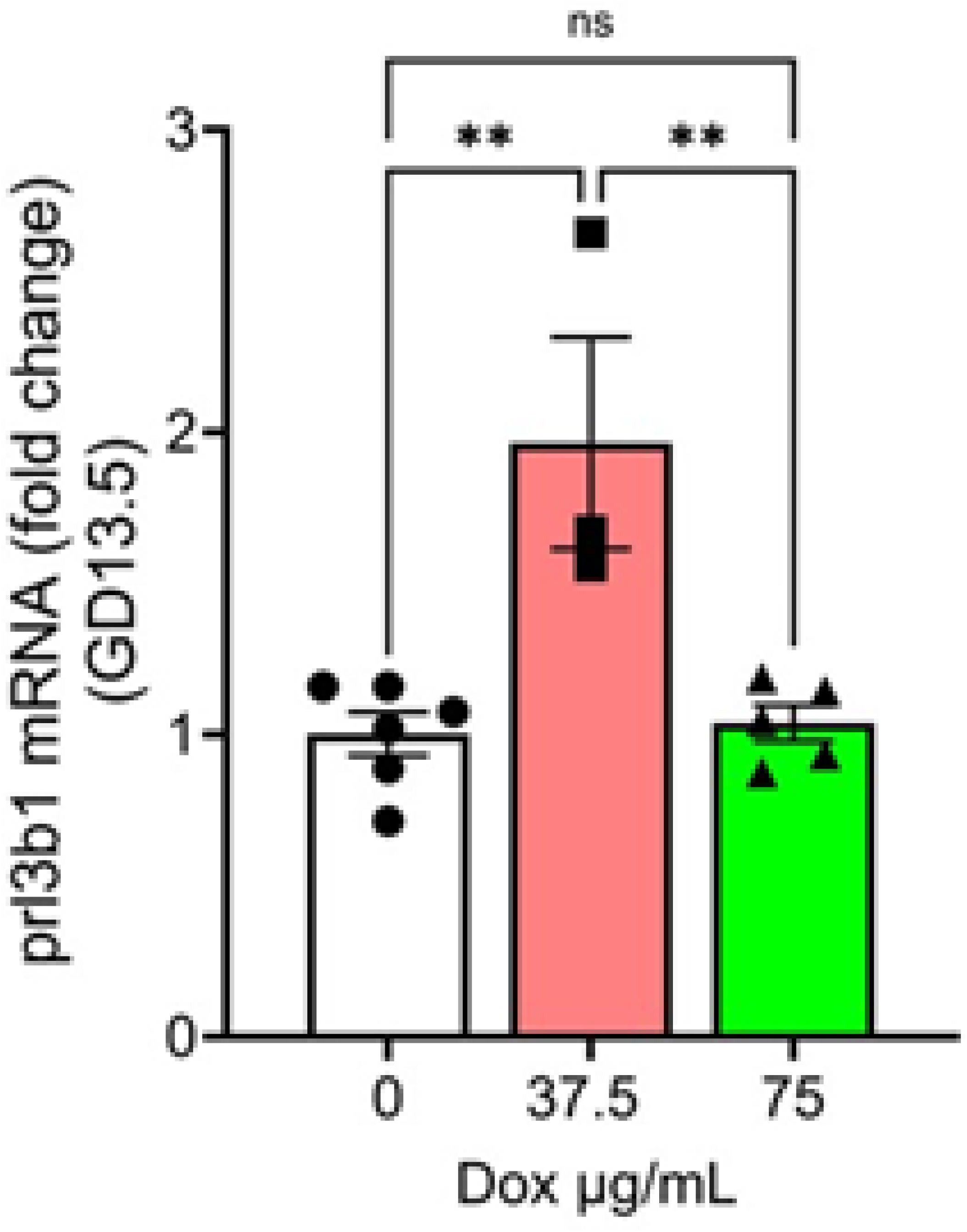
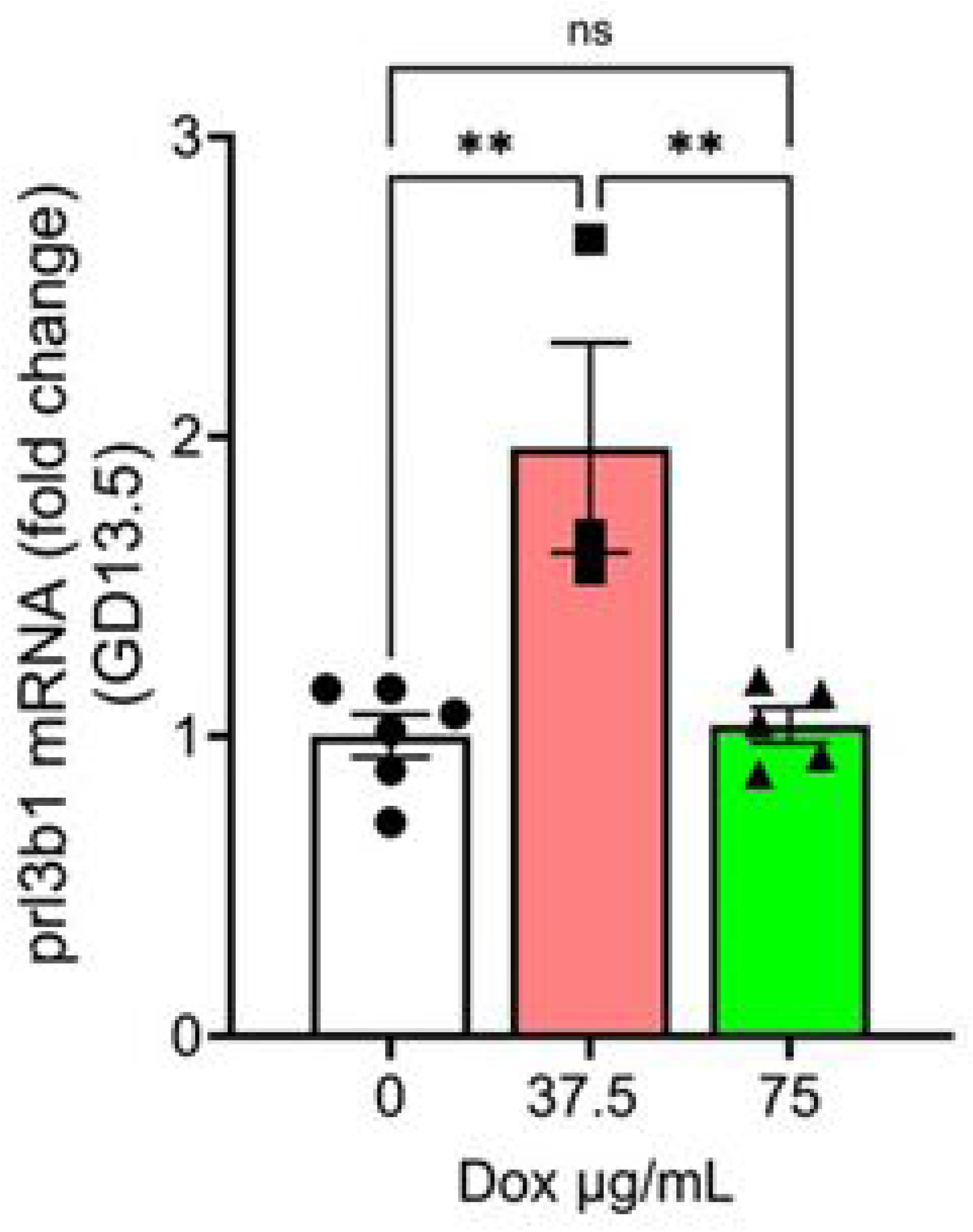
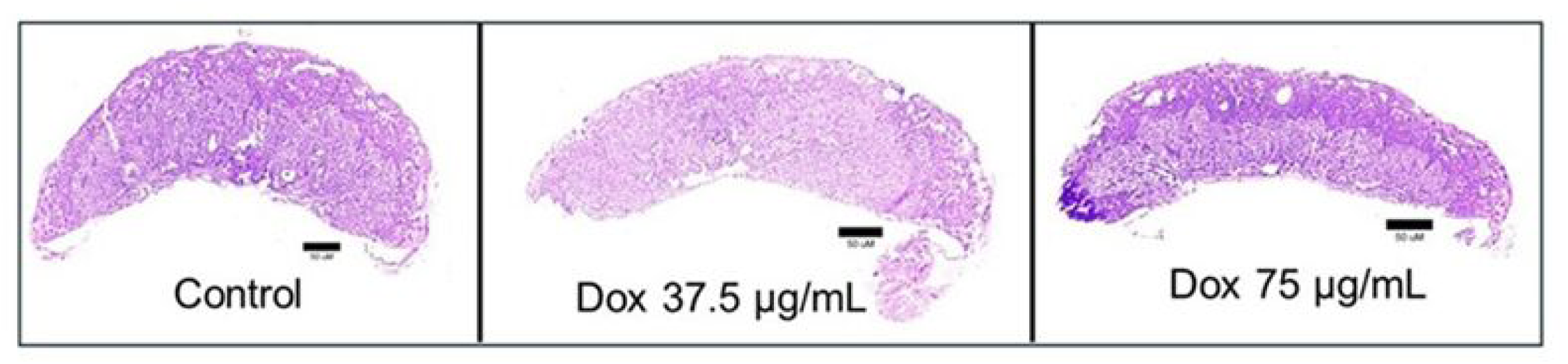
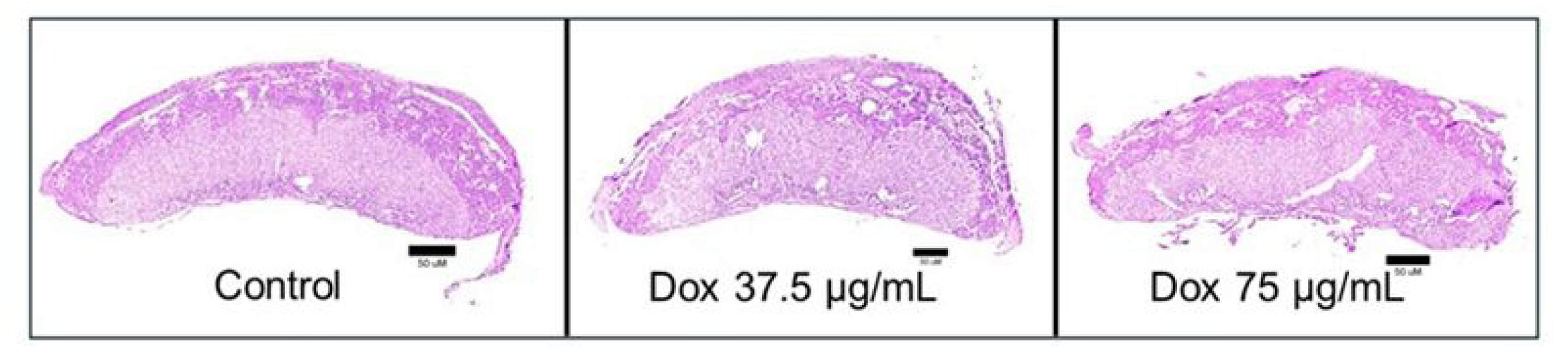
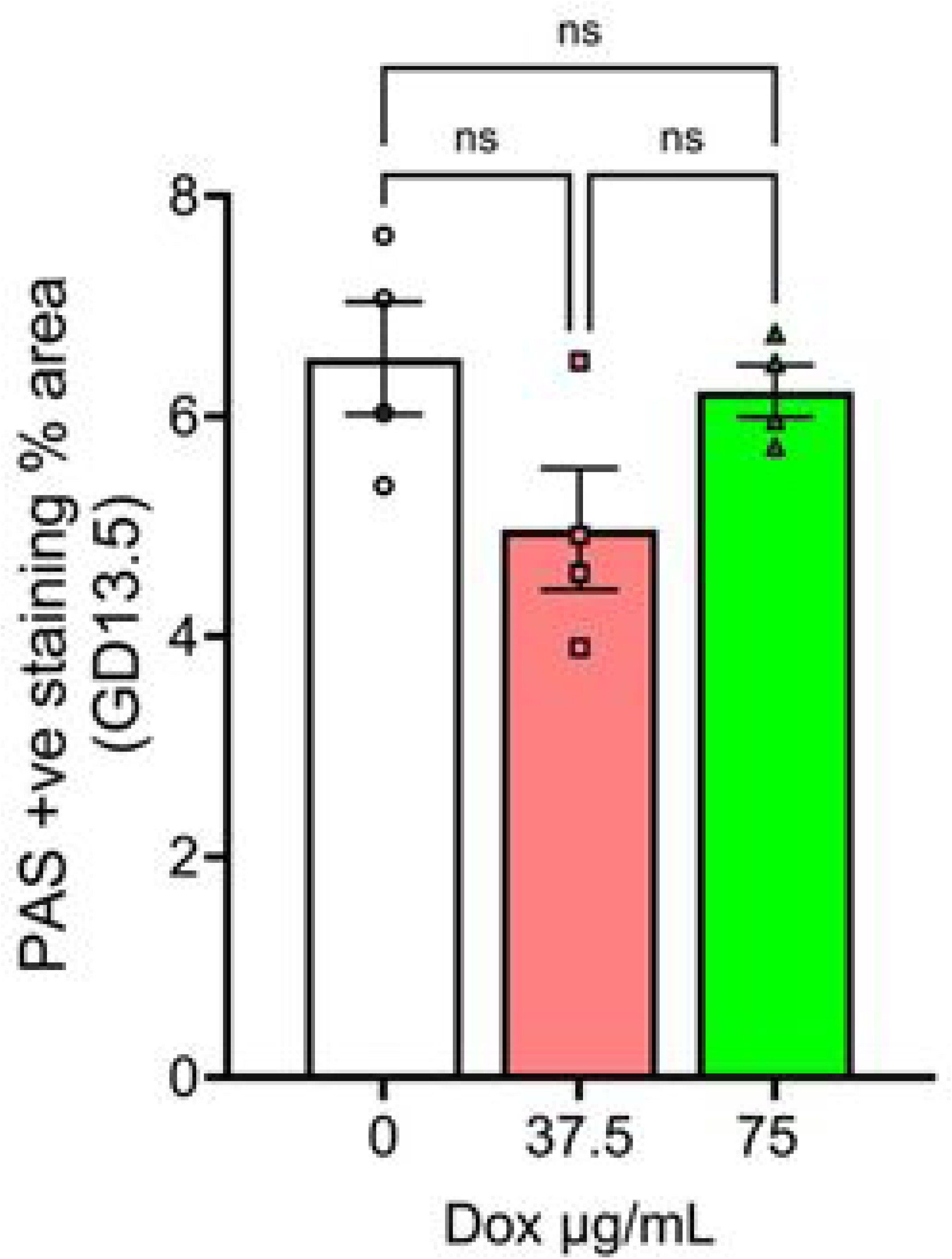
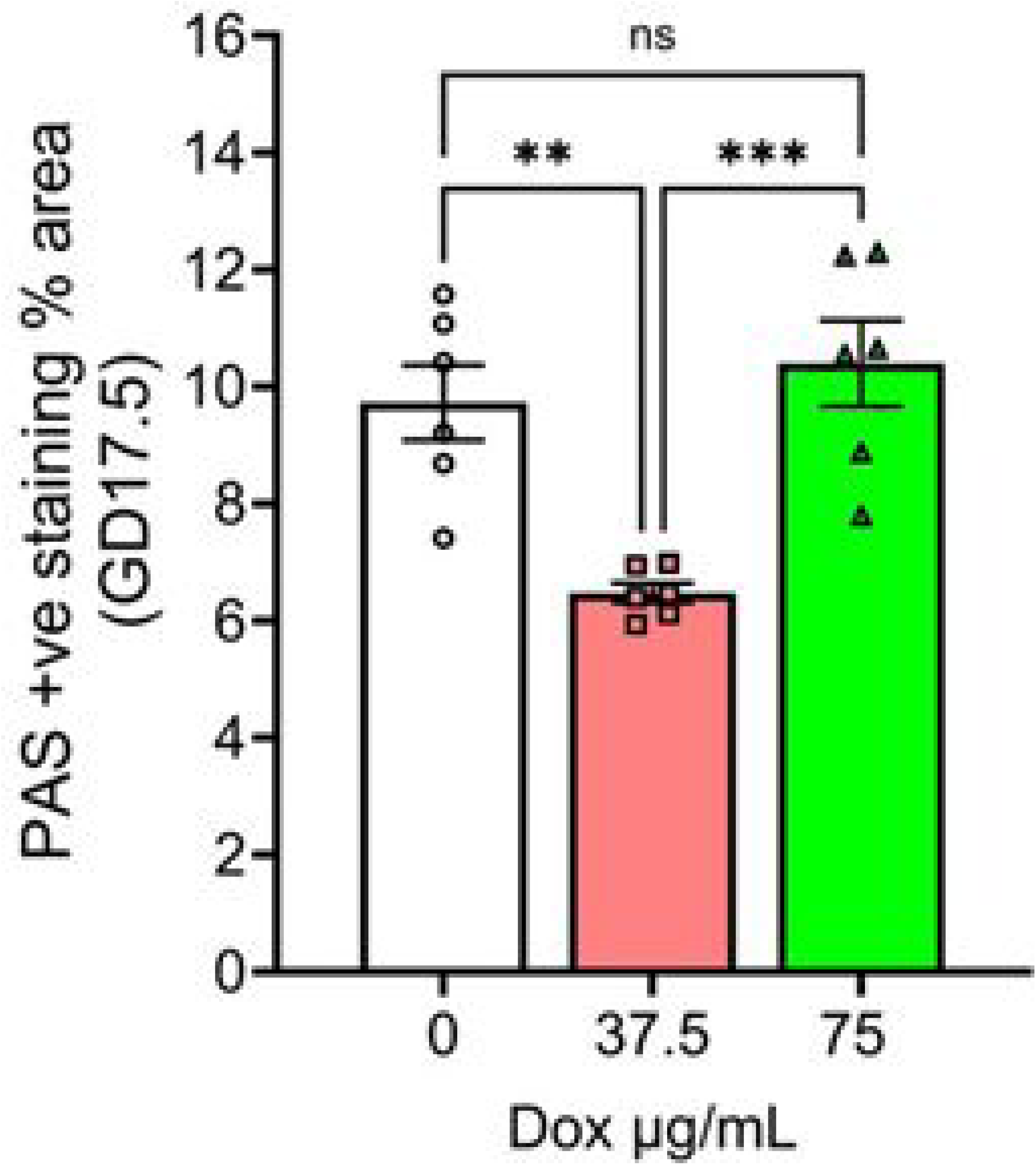
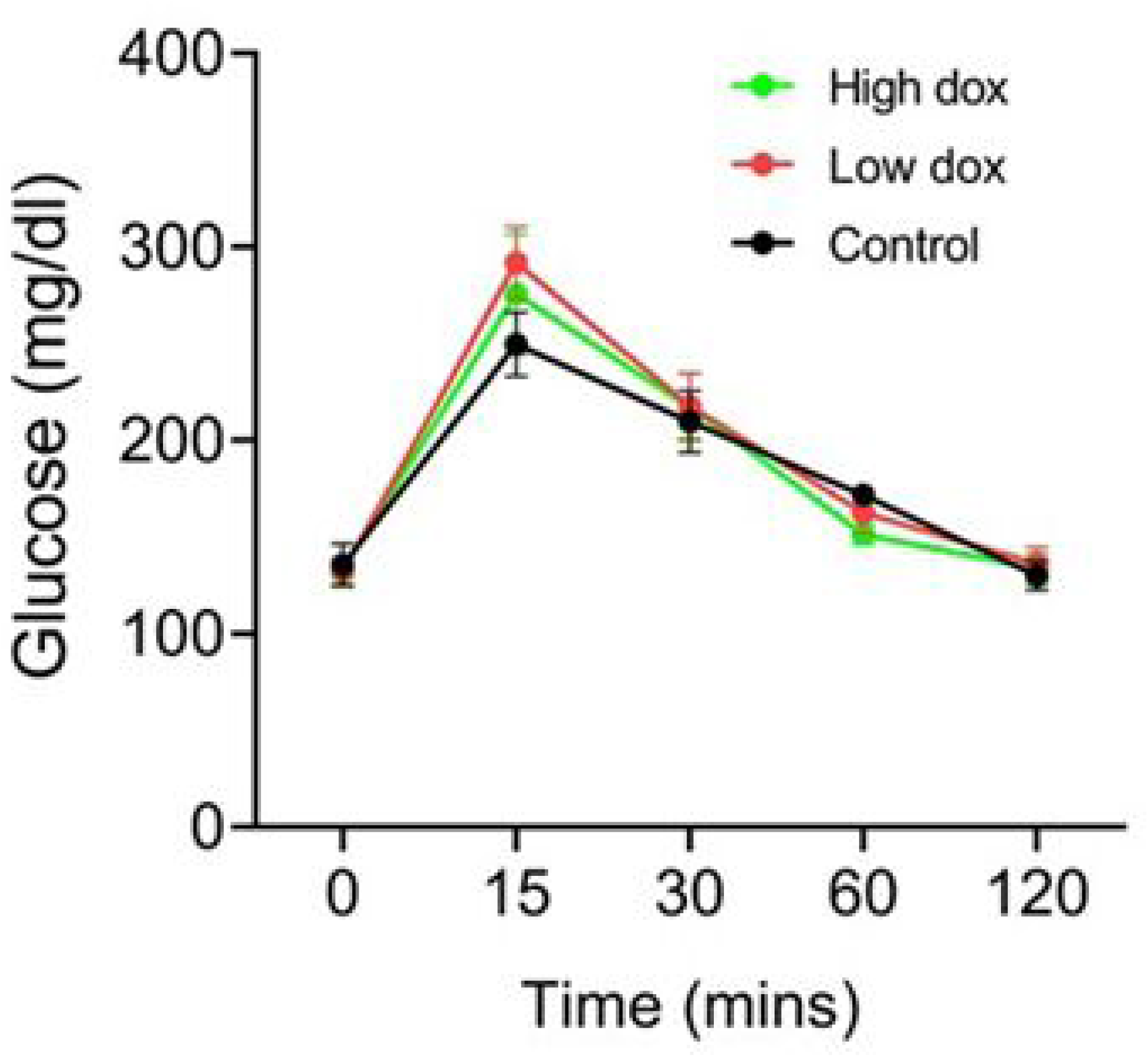
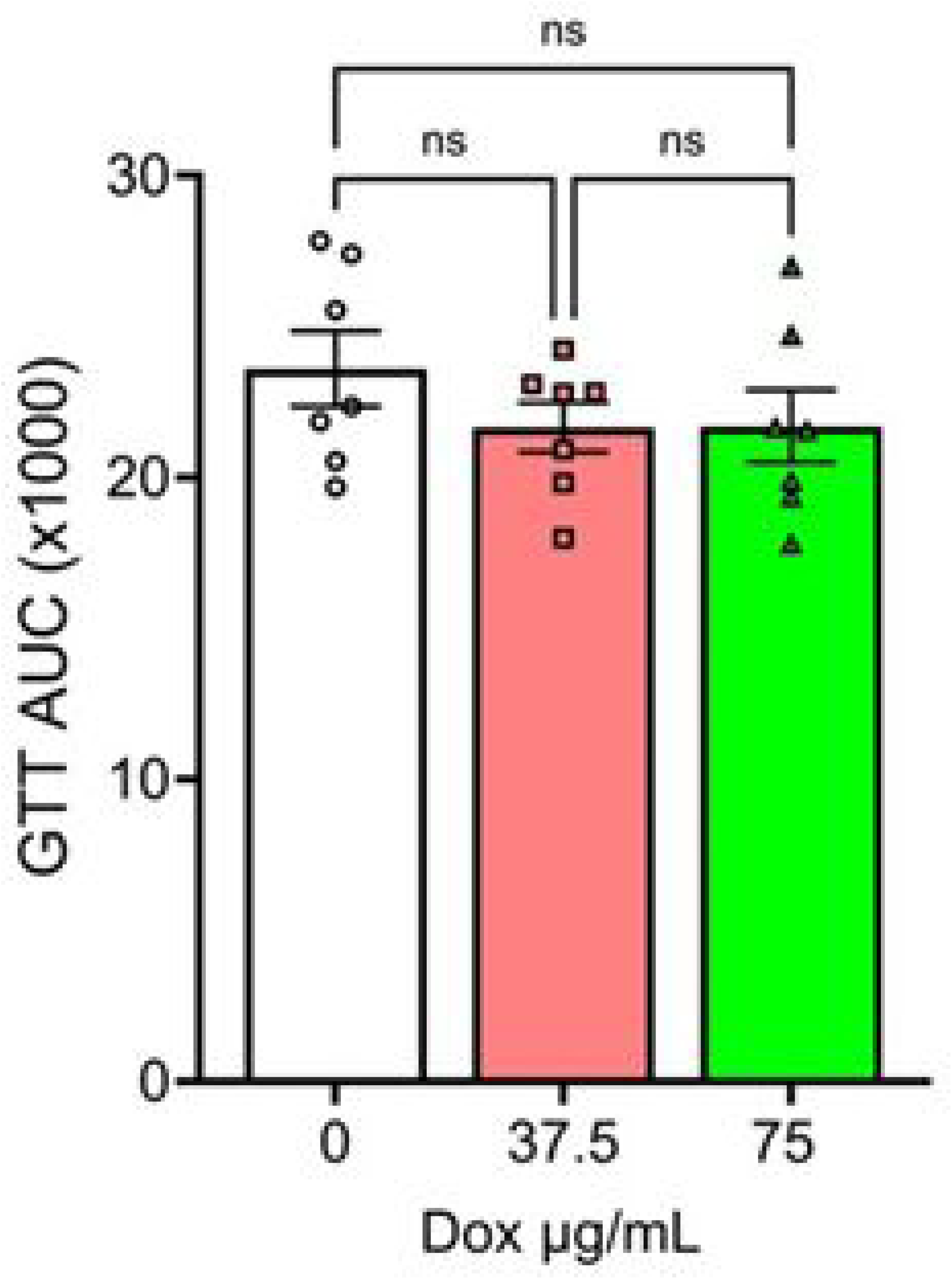
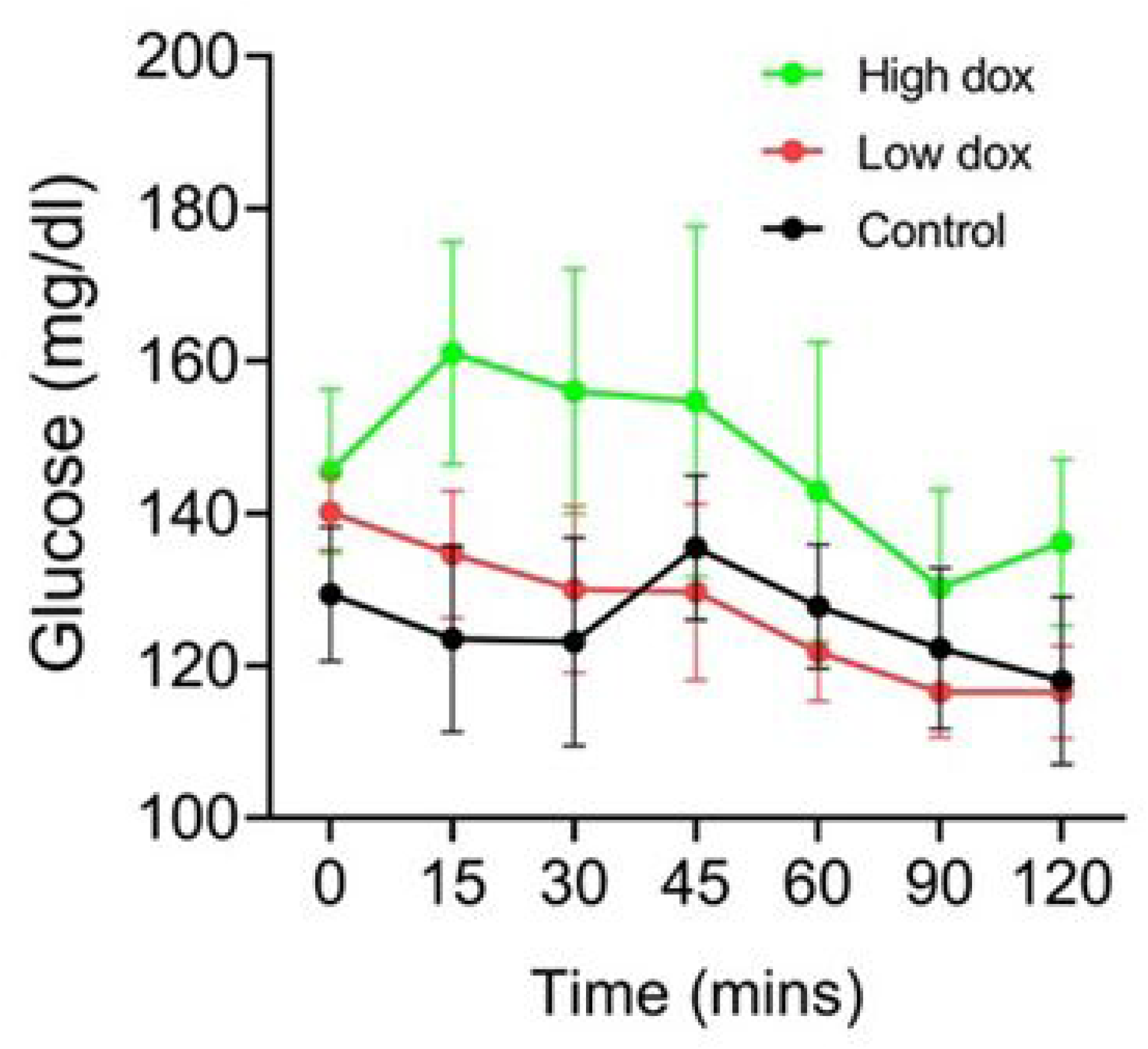
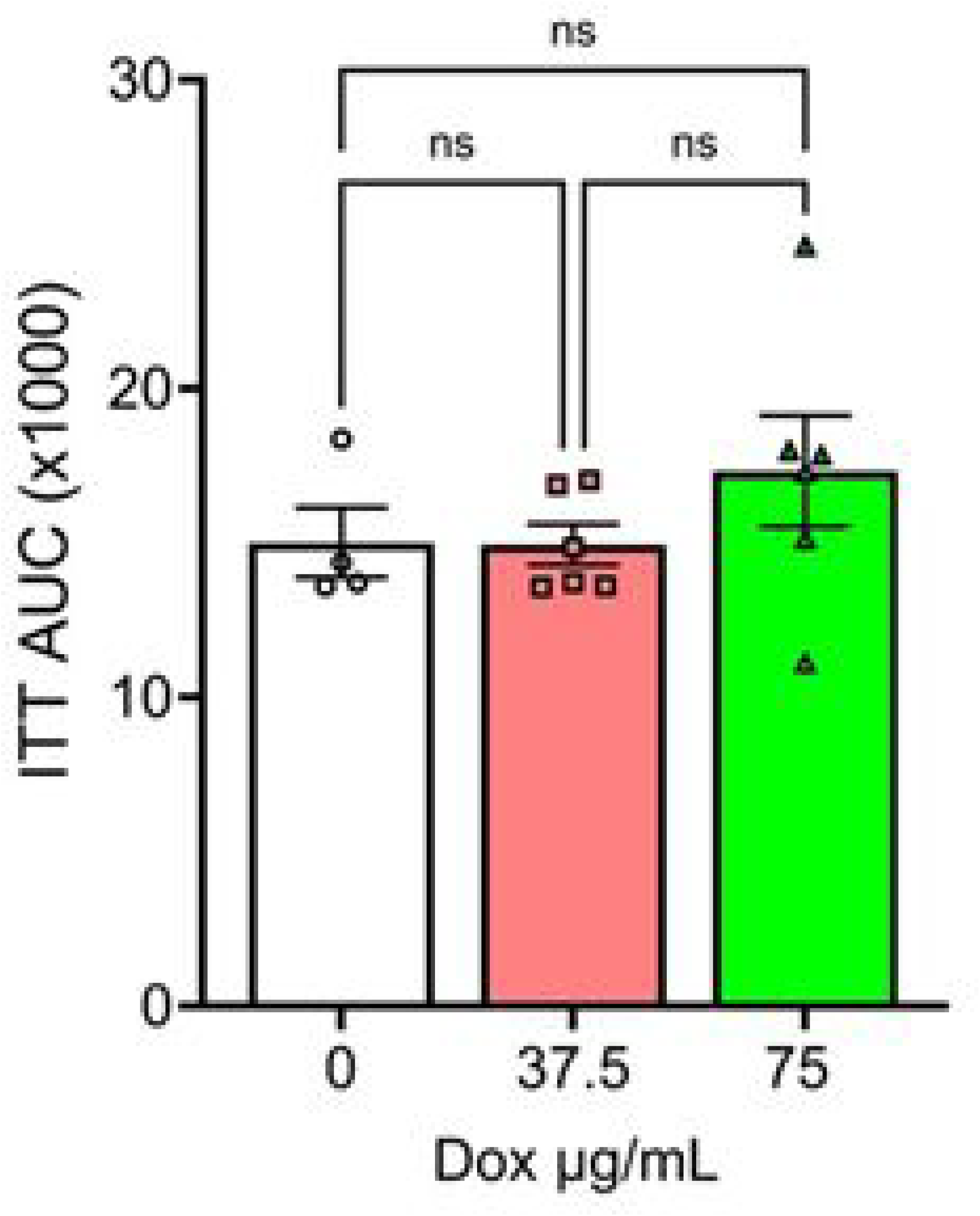
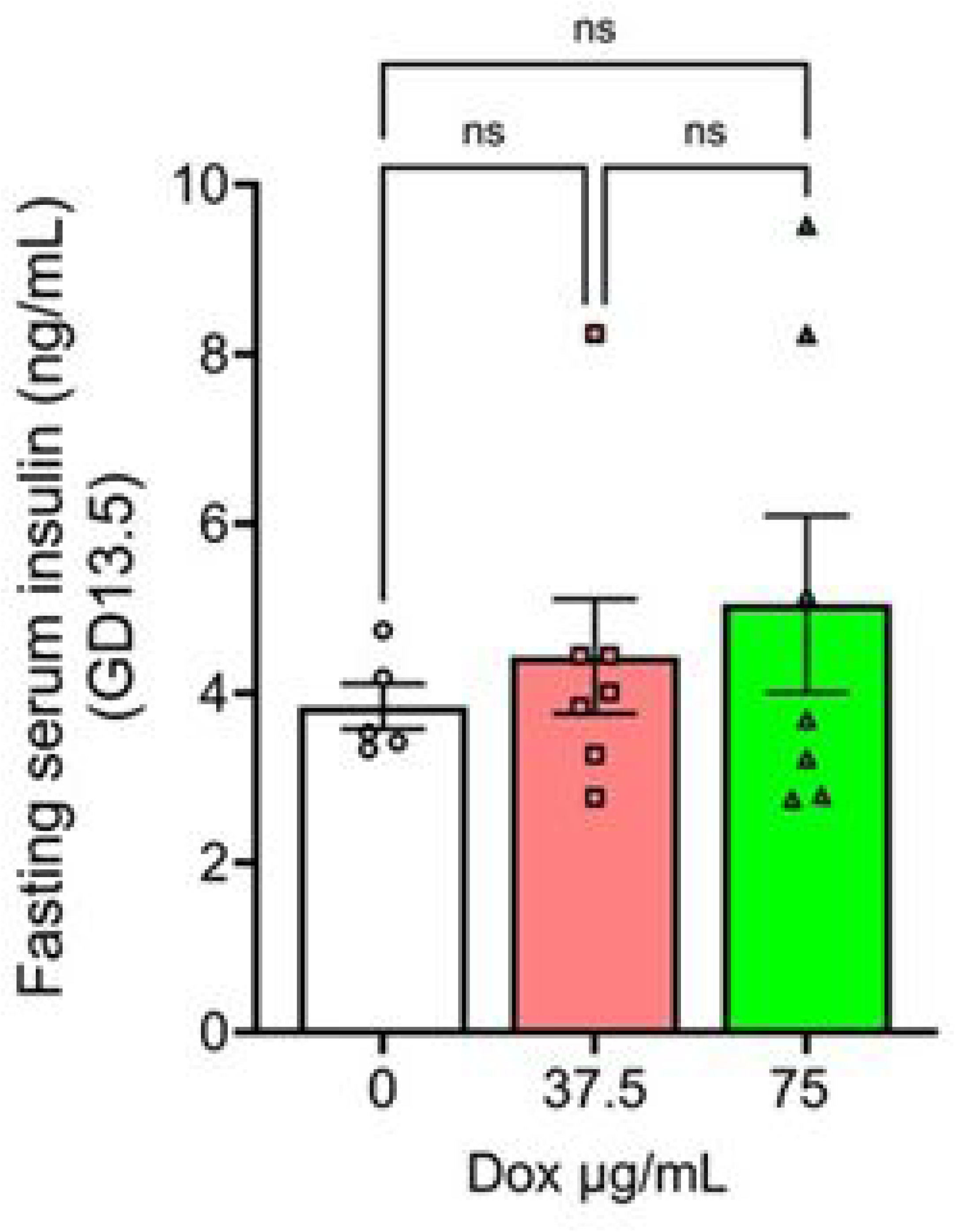
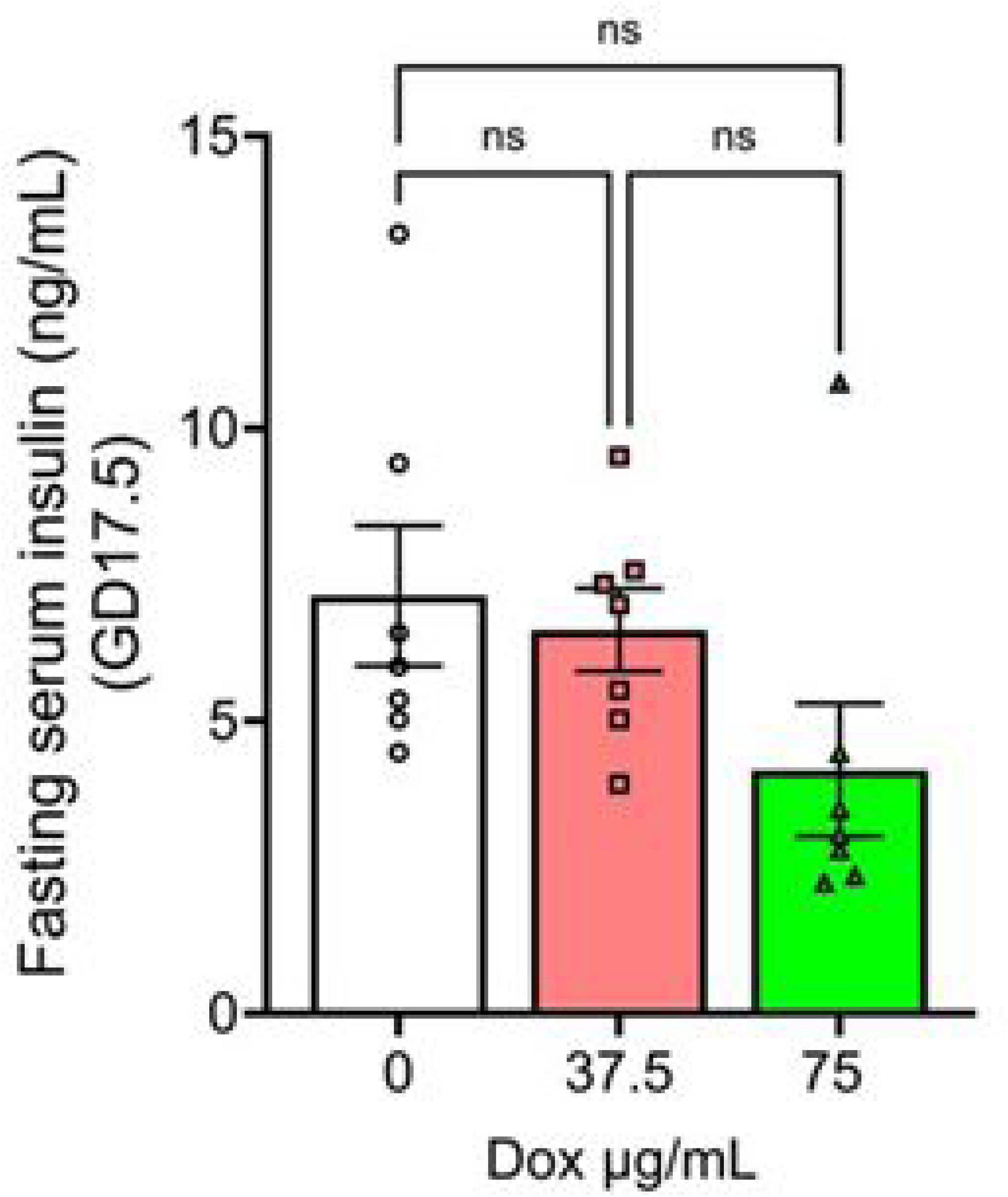
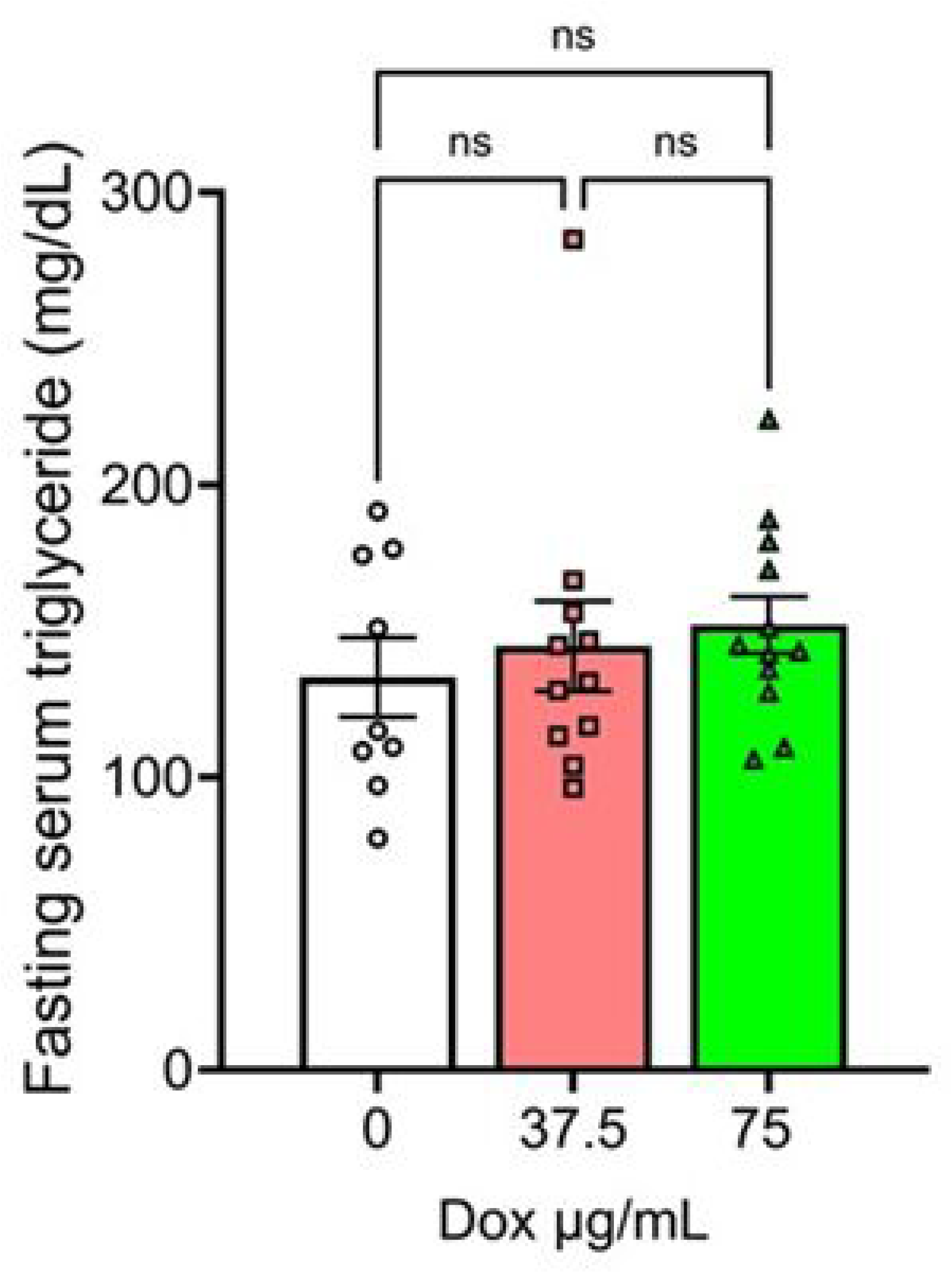
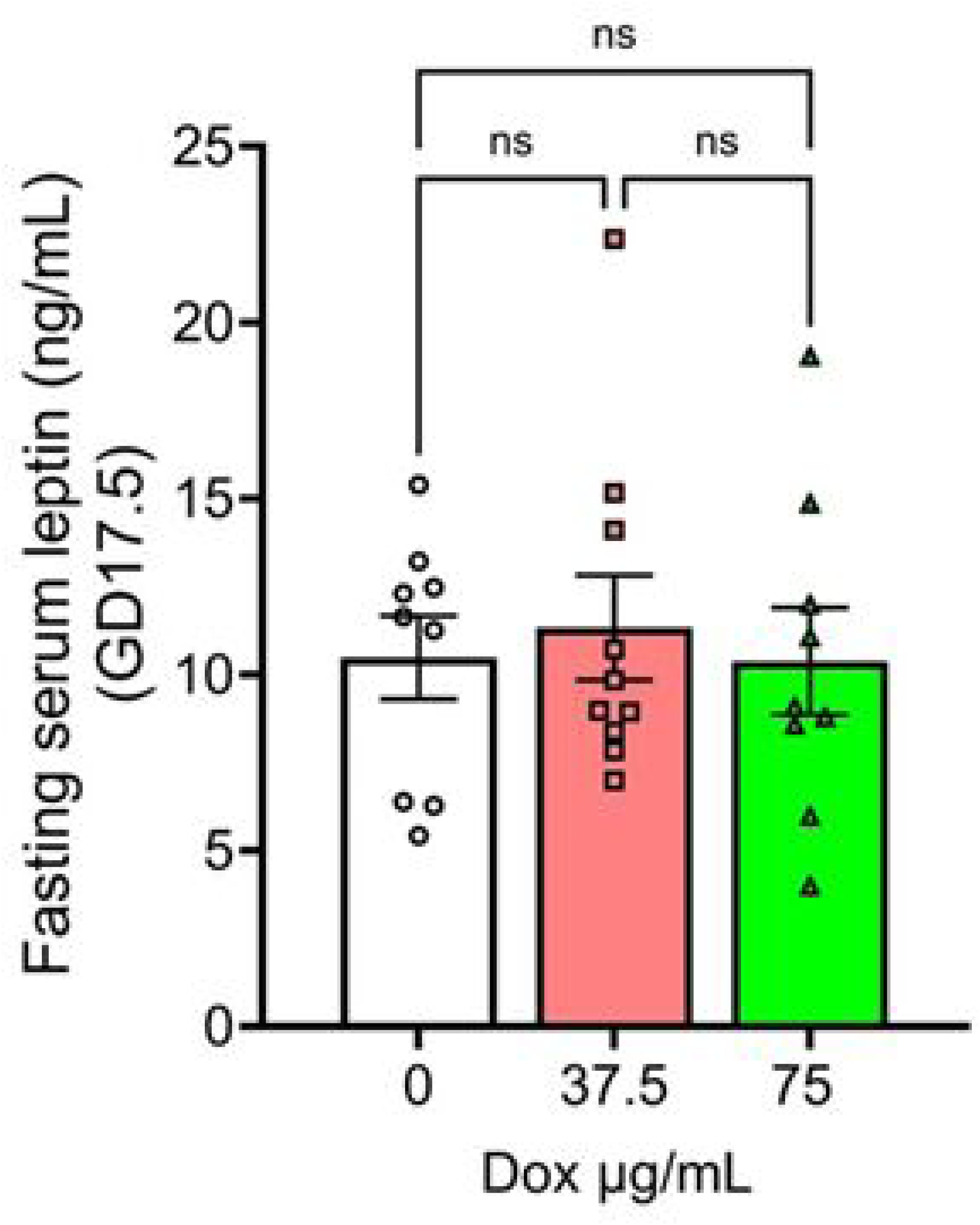
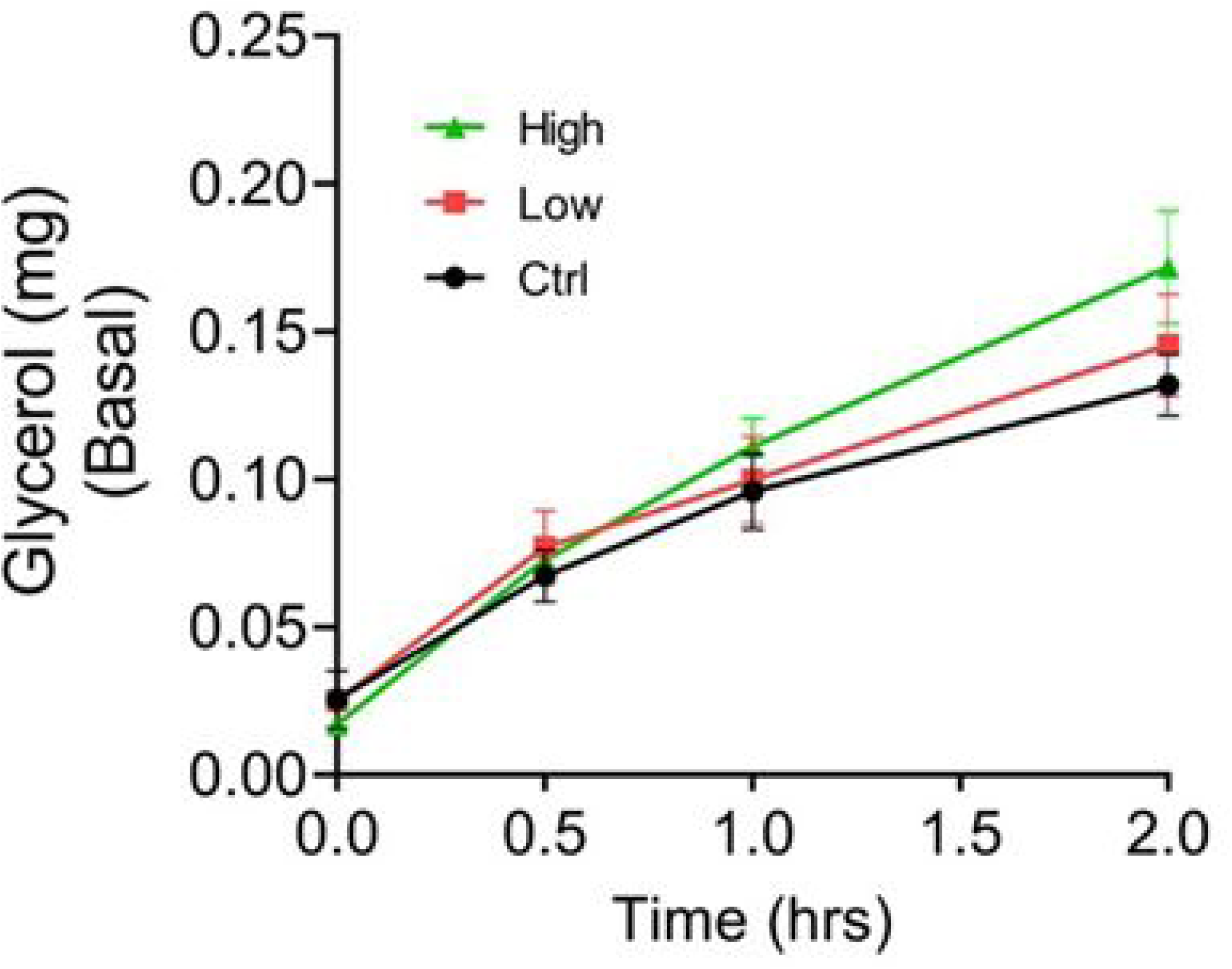
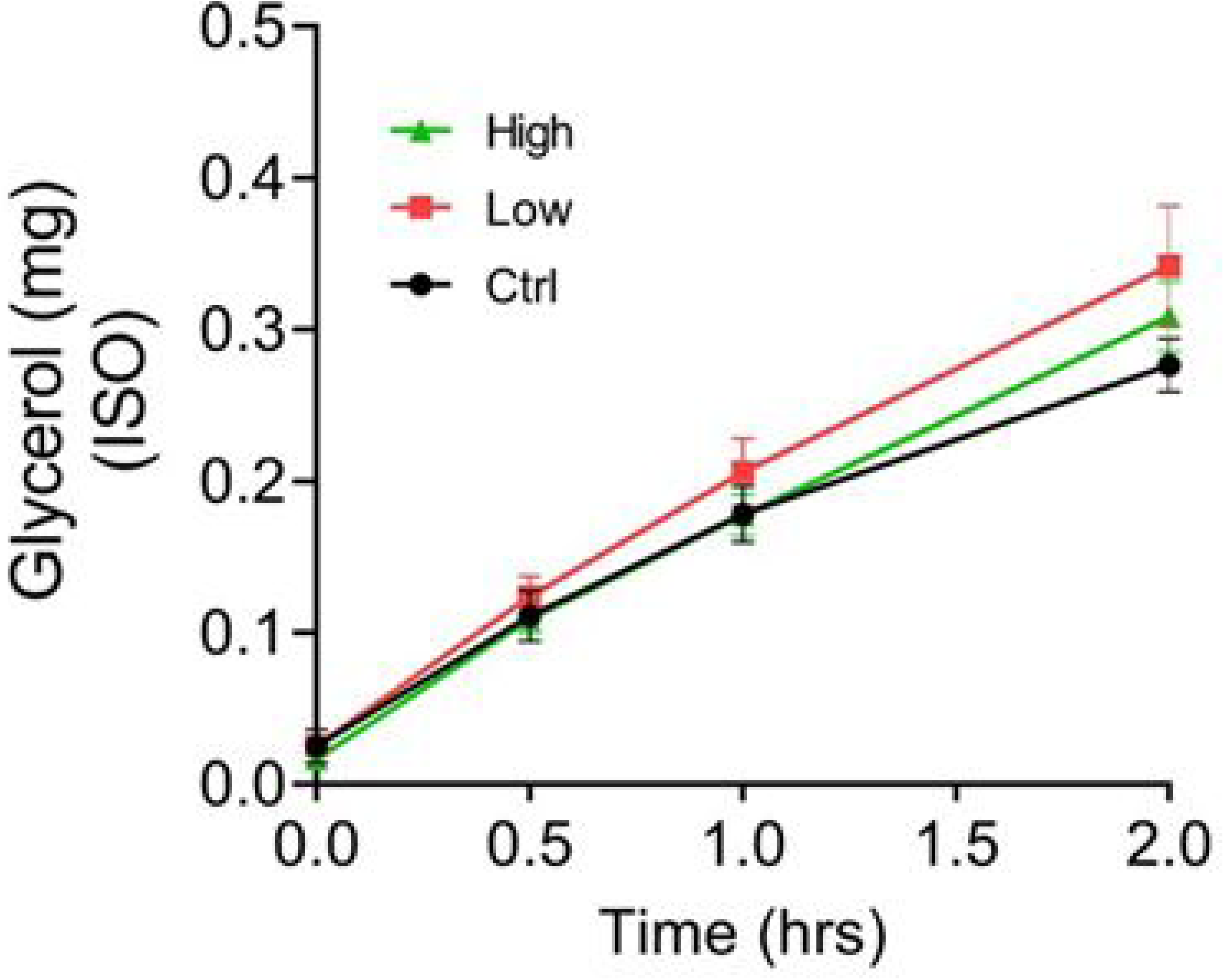
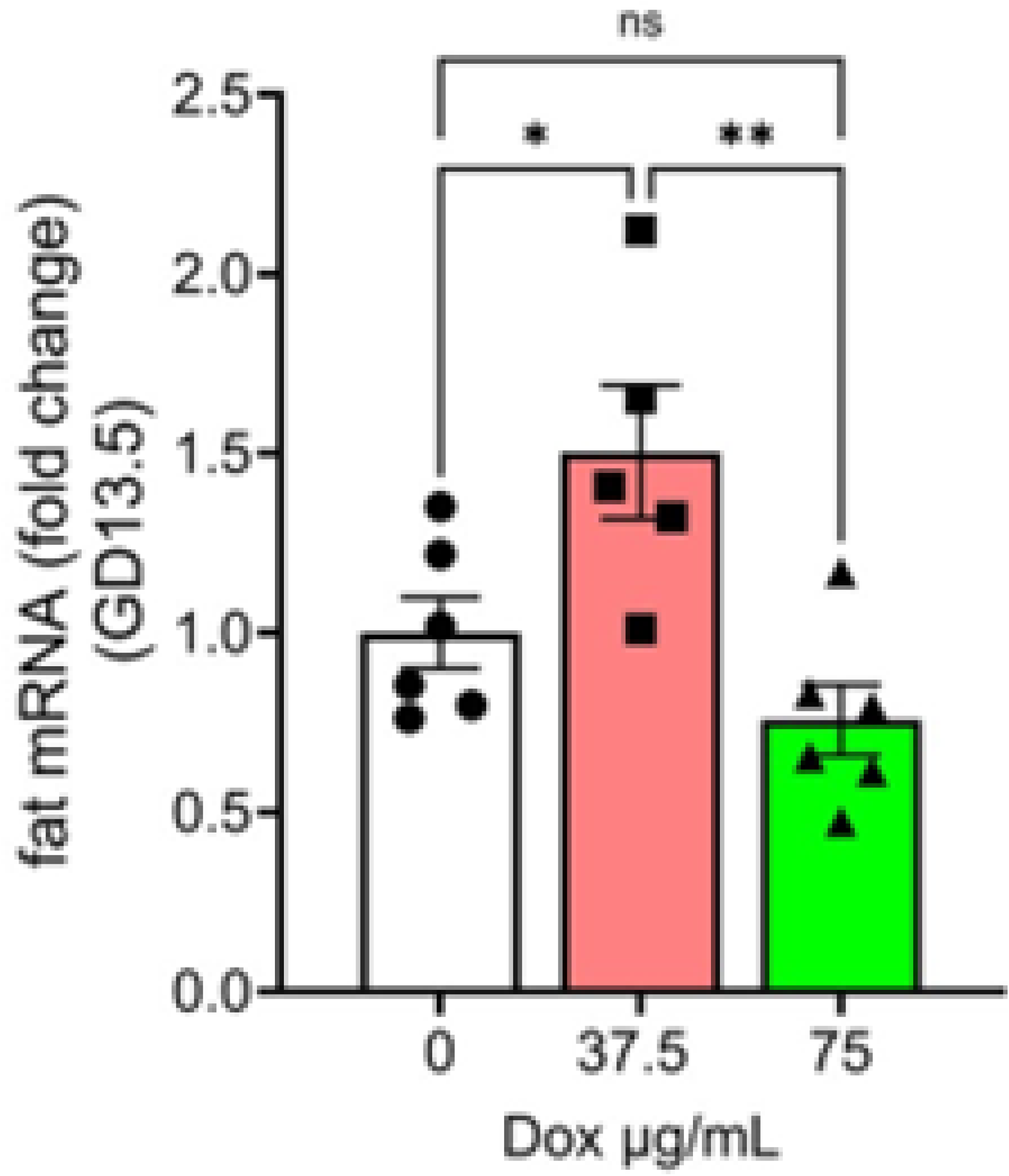
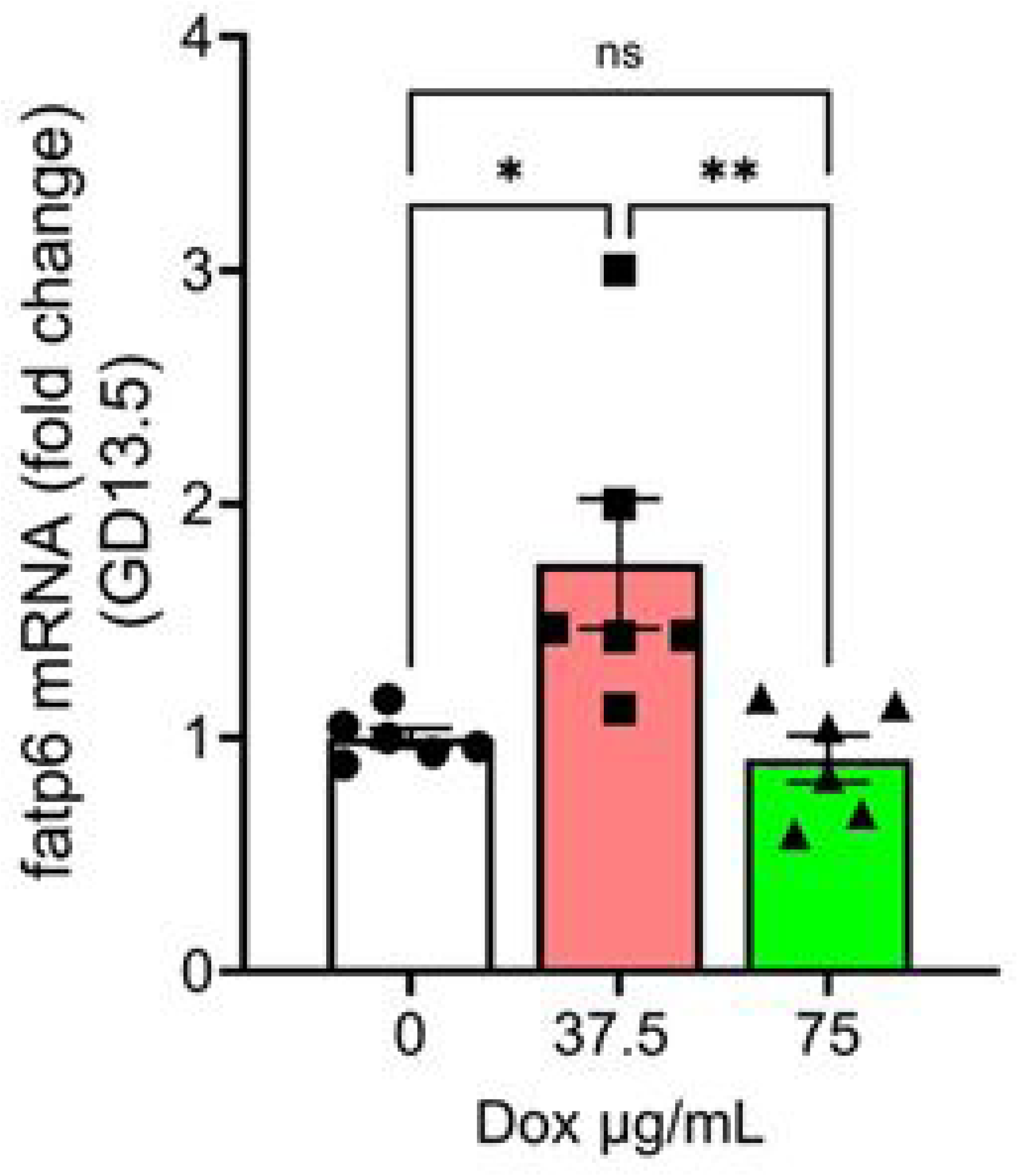
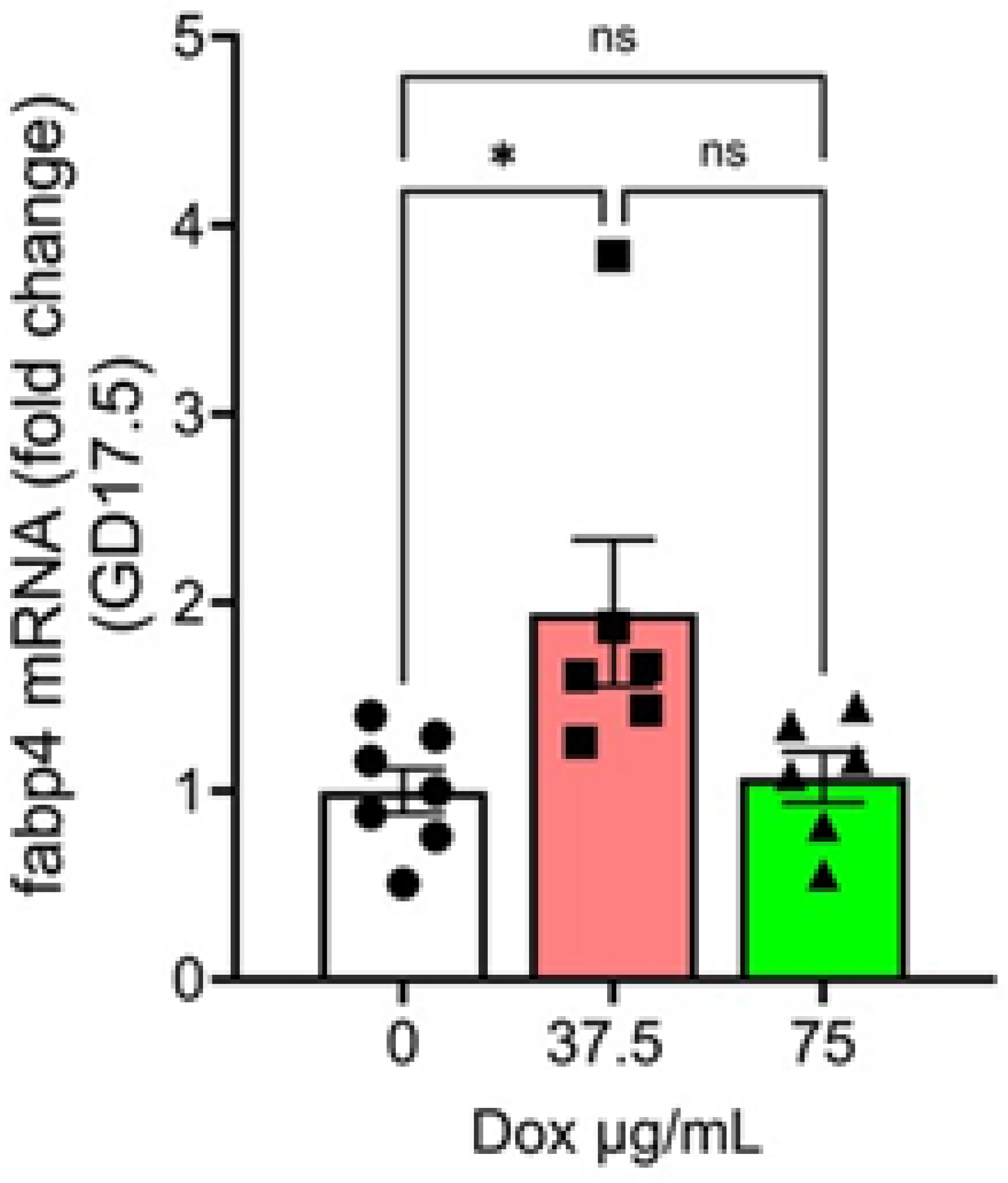
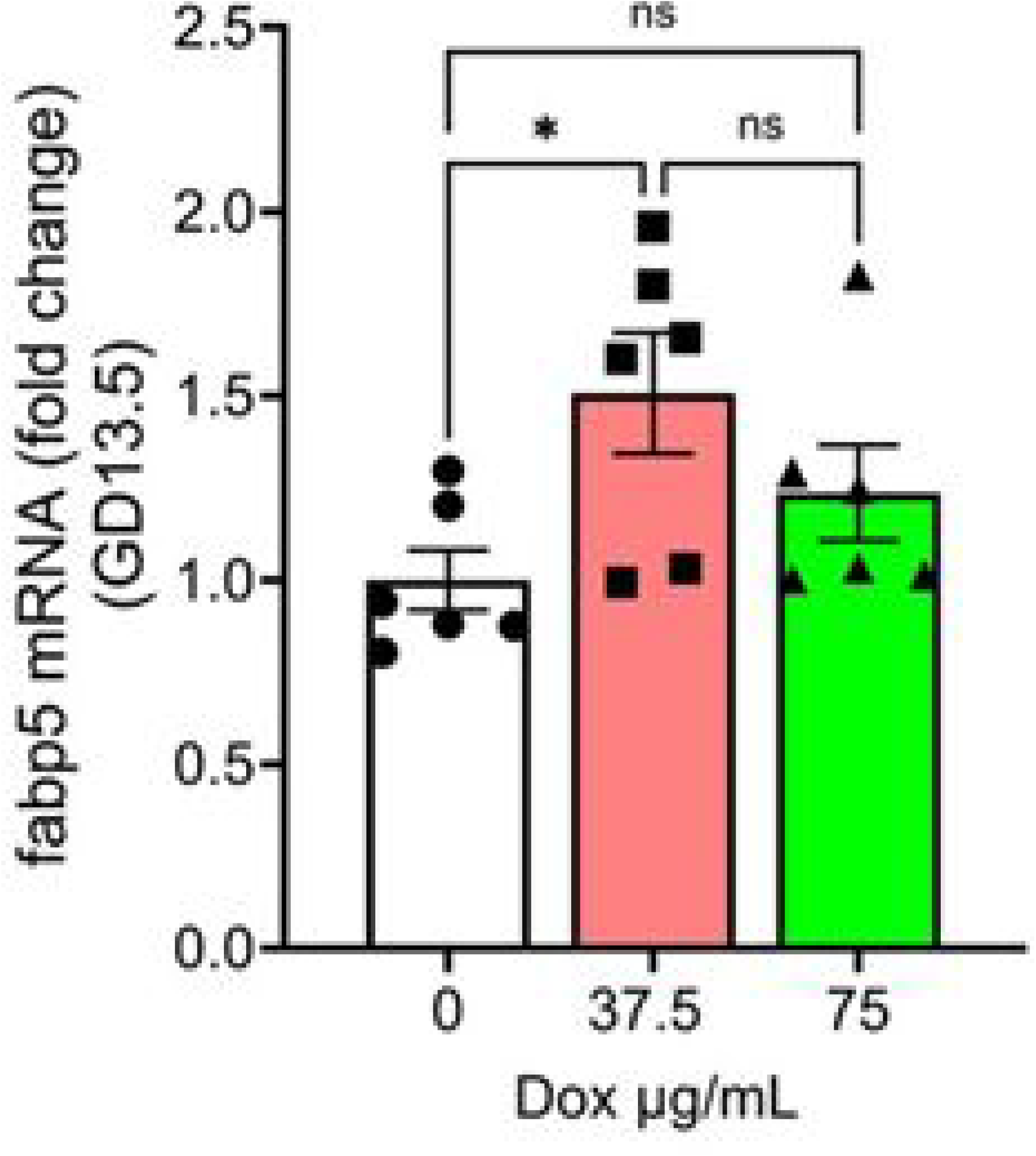
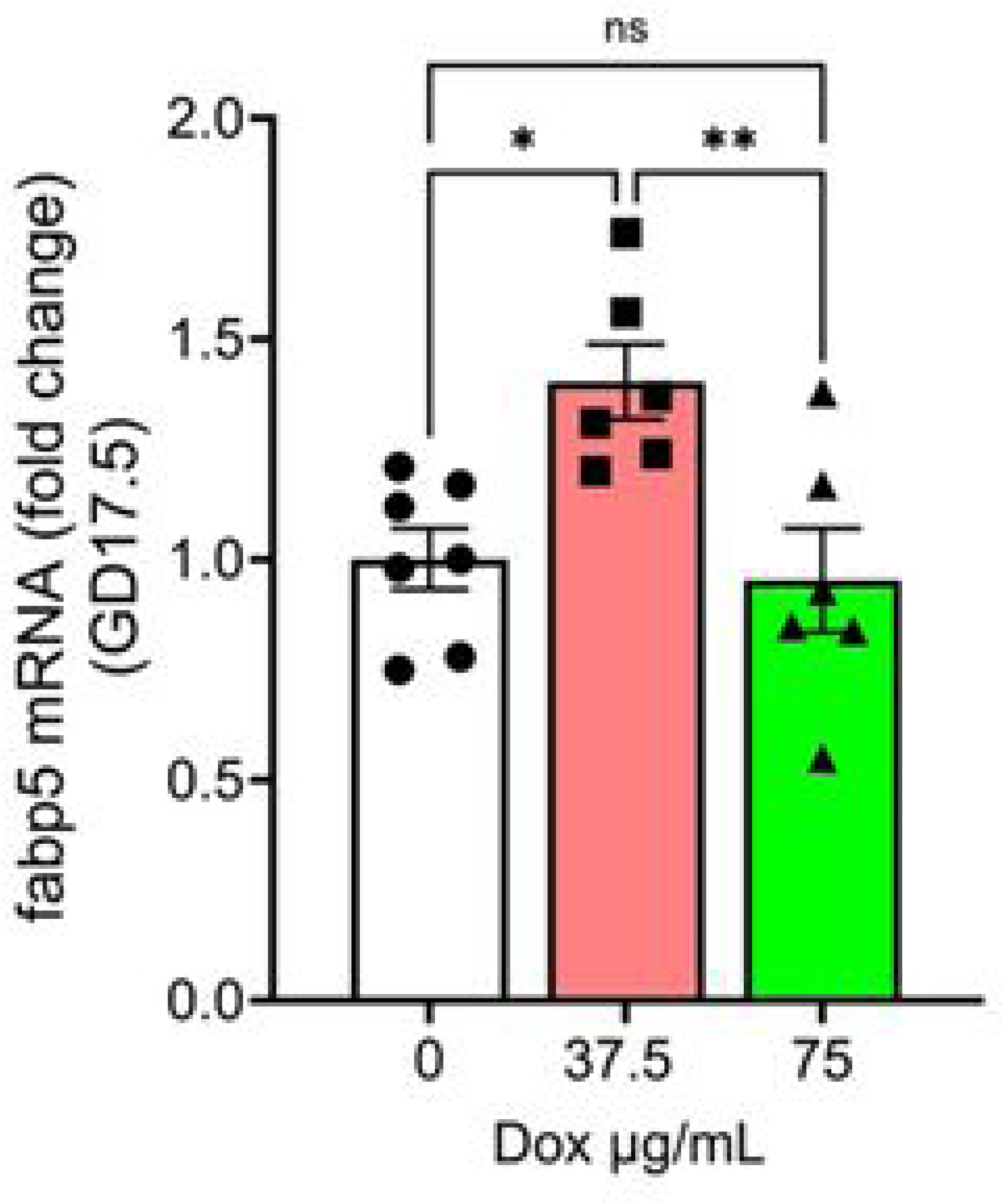

## Notes

### Competing Interest Statement

The authors have declared no competing interest.

